# Green alleys in Quebec provide variable biodiversity support and ecosystem services

**DOI:** 10.1101/2025.07.11.664403

**Authors:** Isabella C Richmond, Kayleigh Hutt-Taylor, Johanna Arnet, Lauren Bianco, Antonia Vieira Zanella, François Bérubé, Paola Faddoul, Étienne Perreault-Mandeville, Nathalie Boucher, Thi Thanh Hiên Pham, Carly D Ziter

## Abstract

Green infrastructure is increasing in popularity in cities globally because of its potential to improve urban sustainability and resident quality of life. In this paper, we studied green alleys in two Quebec cities, one with a resident-led green alley program and one with a municipally led program. Green alleys are conceptualized and promoted as green infrastructure that provide many benefits for urban residents. Using mixed social and ecological methods, we assessed 53 green alleys’ capacity to support biodiversity and provide ecosystem services, alongside 23 grey alleys and 76 streets. We interviewed residents to select the ecosystem services that were most relevant to people living around green alleys and then measured indicators of ecosystem service capacity with traditional ecological techniques, harnessing an interdisciplinary approach to ecosystem service assessment. Green alleys provided more biodiversity support than grey alleys and adjacent street segments but did not consistently increase the capacity for ecosystem services. Vegetative complexity and proportion of native tree species are both higher in green alleys than traditional grey alleys and adjacent streets. The proportion of flowering trees was one indicator of ecosystem services that was consistently higher in green alleys. Resident-led vs municipally led creation and management of green alleys resulted in different results, where resident-led alleys were more able to target the needs of residents but resulted in high levels of variation in both support for biodiversity and ecosystem services. We recommend ongoing funding paired with technical expert support to increase the impact of green alleys.

## 1 Introduction

Green infrastructure is gaining popularity as a tool that can support cities’ sustainability goals (Nelson and Bigger 2022). There is no consensus on the definition of green infrastructure but here we define green infrastructure as “vegetated infrastructure, including phytotechnology and greening practices, used to solve a variety of environmental, social, and economic problems” because we believe this definition outlines the infrastructure and context that we are discussing (Québec Vert 2024). Urban sustainability goals and urban sustainable development have been incorporated into constitutions, laws, and urban development policies globally since the 1980s (United Nations Secretary-General and World Commission on Environment and Development 1987; Ruggerio 2021). Green infrastructure projects often aim to address urban sustainable development goals but we often lack the empirical evidence to evaluate their objectives and performance (Cameron and Blanuša 2016; Jerome et al. 2019). For example, individual green roof projects within a large city may aim to provide several benefits like air pollution reduction, temperature mitigation, improved social well-being, and biodiversity support (e.g., Lampert 2019). However, while there is broad consensus in the literature on temperature mitigation (Wang et al. 2023; Zheng et al. 2023), the remains a lack of research on benefits to air pollution reduction, social well-being, and biodiversity support. Other types of green infrastructure also suffer from this issue, where the list of benefits they aim to achieve for urban residents lack supporting evidence (Cameron and Blanuša 2016; Choi et al. 2021). Although a variety of benefits are plausible, on-the ground monitoring is crucial to establish the impacts of small-scale green infrastructure. Given that green infrastructure and sustainability-focused policies are increasing as the impacts of climate change worsen, collecting evidence on how green infrastructure does and does not address policy goals is critical to informing greening projects.

When assessing the potential benefits of green infrastructure, we can build on the previously established biodiversity-ecosystem service (BES) framework to sort the benefits into two categories: “biodiversity support” and “ecosystem services” (Cardinale et al. 2012). We use these categories because green infrastructure is expected to deliver higher levels of ecosystem services than grey infrastructure (Escobedo et al. 2019) and public policies tend to focus on both ecosystem services and biodiversity support as separate entities (Newell et al. 2013). Here, we define biodiversity support as the benefits of green infrastructure that improve ecosystem functioning and environmental quality resulting in higher levels of biodiversity, essentially “nature for nature” (Kremen and Merenlender 2018). An example of biodiversity support provided by green infrastructure is increased habitat provisioning, which increases the faunal biodiversity of the ecosystem (Hanski 2011). By contrast, ecosystem services are the outcomes of green infrastructure that benefit human residents, such as temperature mitigation and increased access to nature for human residents (Millennium Ecosystem Assessment 2005), that we term as “nature for people”.

In urban areas, previous studies support the claim that green infrastructure has the potential to deliver critical biodiversity support to cities. For example, green infrastructure can increase the green space abundance, diversity, and complexity present in dense urban areas that may otherwise be hostile to flora and fauna due to the amount of built infrastructure (Pinho et al. 2021). However, the design, management, and built environment surrounding different green infrastructure types will inevitably determine the quantity and quality of biodiversity support provided (Zuniga-Teran et al. 2020; Pinho et al. 2021). For example, developing school yards as small green space pockets across a city can significantly increase connectivity, by providing habitat and supporting biodiversity (Iojă et al. 2014). However, a single school yard within a matrix of heavily built-up land will not provide the same magnitude of biodiversity support as the network. Currently, the influence of design and management on the levels of biodiversity support provided by green infrastructure is not well understood, and thus we do not know the actual benefits that many types of green infrastructure are providing.

Turning to ecosystem services, they are often a focus of city managers and planners as they increase the quality of life of urban residents in important and varied ways (Geneletti et al. 2020). Ecosystem services are diverse and can include regulating services such as air temperature regulation, cultural services such as opportunity for recreation, and provisioning services such as food production (Millennium Ecosystem Assessment 2005).One of the current gaps in ecosystem services studies is the connection between perceived or desired ecosystem services and the ecosystem services that are actually quantified by scientists. Studies are often siloed by discipline, either focusing on the psychological/sociological aspect of residents’ preferences and desires for different types of ecosystem services or the ecological aspects of measuring capacity of an ecosystem to provide a list of ecosystem services that the scientists themselves have derived (Chen et al. 2024). There is a dearth of studies where researchers quantify the ecosystem services that residents have informed them are desirable or preferred, and this, despite the fact that asking the local community for their input and separating biodiversity and ecosystem service assessments have been highlighted as important methodological approaches (van Oudenhoven et al. 2018; Schröter et al. 2021). Far more often, ecologists select ecosystem services to study based on factors that exclude resident experience or perception, such as data availability, prevalence in the literature, or human health impacts. Yet, residents often have complex relationships with nature that determine what they desire as ecosystem services and what are disservices for them (Roman et al. 2021; Drew-Smythe et al. 2023) which are also dependent on a person’s social and ecological context (Quintas-Soriano et al. 2018; Cebrián-Piqueras et al. 2020). Therefore, as cities grow and increase pressure for ecosystem services, understanding which services are desired by residents is a critical step in the implementation and management of local green infrastructure.

This brings us to a broader discussion about the rapport between humans and nature and the conceptualization of biodiversity and ecosystem services. In this paper we propose that biodiversity support can be seen as “nature for nature” and ecosystem services as “nature for people”. Both concepts are needed in sustainability policies, they can however be opposing in terms of management needs (Nassauer 1995; Ziter 2016). The opposition occurs when human preferences and norms result in green spaces that do not have capacity for biodiversity support (Nassauer 1993; Larson et al. 2020; Lis et al. 2022). For example, people may prefer manicured turf grass lawns over wildflower meadows or native grasses due to aesthetic preferences and social pressures, despite wildflower meadows and native grasses supporting more biodiversity (Nassauer et al. 2009; Simmons et al. 2011). Another example is the emphasis on social cohesion, where people desire for their yard to look like their neighbours’, regardless of ecological function or biodiversity (Nassauer et al 2009). This said, the relationship between human desires and biodiversity support is not always antagonistic. Researchers and practitioners have also found synergistic relationships between desired ecosystem services and biodiversity support (Shen et al. 2020; Karimi et al. 2021; Khosravi Mashizi and Sharafatmandrad 2021). The increased research on synergies between biodiversity support and ecosystem services provided by green infrastructure can help cities meet multiple sustainability goals.

As a result of their potential capacity for synergies between biodiversity and ecosystem services, green alleys are a type of green infrastructure North American cities are increasingly investing in (Newell et al. 2013). Alleys provide an excellent case study to assess green infrastructure. Alleys have had many uses throughout history including ice/coal delivery, garbage pickup, and socializing (Wolch et al. 2010). A key benefit of green alleys is that alleys exist within even the densest parts of our cities, reducing the need to remove built infrastructure to incorporate green infrastructure. Further, given that alleys already exist as important social and recreation spaces (Forde et al. 2024), transforming an alley into a form of green infrastructure is usually met with resident support (Seymour et al. 2010; Brazeau-Béliveau and Cloutier 2021; Forde et al. 2024). The development and governance of green alleys varies across location, from exclusively municipal management to entirely community driven (Newell et al. 2013), resulting in green alleys looking very different across and within cities (Pham et al. 2022; Forde et al. 2024). However, we have limited information on how the functioning of alleys differs in response to their variable appearances.

In Montreal, Canada, for example, green alleys are resident-led; once an alley is approved to join the program, the borough provides an initial input of funding which residents can use to “green” their alley as they wish. This format results in very variable management by residents, ranging from installation of planter boxes and trees to additions of wall murals and play equipment (Pham et al. 2022). On the other hand, municipally led green alley programs often have a specific goal, and thus more uniform designs (Newell et al. 2013). For example, the green alley program in Trois-Rivières, also in Quebec, is municipally led and focused mainly on delivering stormwater management (Ville de Trois-Rivières 2023). Given the unique form and variation in alley appearance, using established ecological methods for parks or yards and extrapolating information from other green infrastructure types is not appropriate (Dade et al. 2024). We expect that green alleys vary ecologically, but the variation in provision of biodiversity support and ecological services in green alleys is a question that, to our knowledge, has yet to be explored.

Our study considers the biodiversity support and ecosystem services provided by publicly owned green alleys in two Canadian cities, Montréal and Trois-Rivières, under two forms of governance: resident-led (with municipal support) and municipally led. We surveyed residents to understand their perceived and desired ecosystem services and measured multiple biophysical indicators in green alleys and adjacent street segments (Table 1) to assess the extent to which alley greening changes the level of biodiversity support and ecosystem services (together “ecological benefits”) provided by these spaces. We asked:

1. To what extent do green alleys provide more biodiversity support (i.e., “nature for nature”) than comparable grey infrastructure such as grey alleys and streets?
2. To what extent do green alleys provide more ecosystem services (i.e., “nature for humans”) than comparable grey infrastructure such as grey alleys and streets?

**Table 1.** Summary statistics of the measures of interests across both cities. Standard deviations are presented with means.

| **Index** | **Villeray-Saint Michel-Parc**<br>**Extension** | **Trois-Rivières** |
| --- | --- | --- |
| Number of sites (all types) | 100 | 26 |
| *Biodiversity support* |  |  |
| Average canopy cover | 33 ± 21 % | 18 ± 18 % |
| Tree species richness | 70 total species | 22 total species |
| Number of functional groups<br>(trees) | 9/9 | 5/9 |
| Average proportion of native<br>trees | 61 ± 30 % trees native | 75 ± 50 % trees native |
| Average proportion of invasive<br>trees | 14 ± 21 % trees invasive | 21 ± 42 % trees invasive |
| Average vegetative complexity | 1.40 levels / sampling point | 1.31 levels / sampling point |
| Firefly presence | 12/54 sites with fireflies | NA |
| *Ecosystem service indicators* |  |  |
| Average daytime temperature | 24.4 ± 3.6°C | 20.3 ± 4.9°C |
| Average nighttime temperature | 21.9 ± 2.5°C | 16.3 ± 4.1°C |
| Total tree abundance | 1,371 total trees | 16 total trees |
| Average # trees / site | 13 ± 13 trees | 0 ± 1 tree |
| Average diameter at breast<br>height (DBH) | 19.3 ± 17.2 cm | 21.2 ± 17.2 cm |
| Average potential maximum<br>height | 14.0 ± 1.3 m | 14.4 ± 2.8 m |
| Average proportion of<br>flowering trees | 22 ± 25 % flowering trees | 4 ± 8 % flowering trees |

We hypothesize that green alleys will provide higher levels of both biodiversity support and ecosystem services than grey alleys or streets due to active management initiatives and an input of funding. However, we predict that in areas where alley design and management is bottom-up, i.e., Montréal, the alleys will be more variable in their vegetation management and therefore have more variable levels of ecosystem services and biodiversity support than in top-down programs, i.e., Trois-Rivières.

## 2 Materials & Methods

### 2.1 Site Selection

#### 2.1.1 Montréal

Our study area was the Montréal borough of Villeray—Saint-Michel—Parc-Extension (VSMPE), composed of three distinct neighbourhoods and a total population of ∼144,000 with a population density of 8,723 residents per km^2^ (Montréal en statistiques et al. 2018). Parc-Extension is considered the most densely populated and poorest neighbourhood in Montreal.

Saint-Michel is a low-income neighbourhood whose population is composed of 50% immigrants with low education levels. Villeray has a higher proportion of francophone residents than Saint Michel, with some high-income and highly educated areas (Centraide of Greater Montreal 2020).

In total, there are 72 green alleys in VSMPE (Regroupement des éco-quartiers 2023). We surveyed 40, spread across the three neighbourhoods geographically to represent each of the neighbourhoods and span a range of canopy cover (Figure 1). We also selected 10 grey alleys that do not have a green alley designation and were similarly spread across the three neighbourhoods and a range of canopy cover to act as control sites and facilitate comparison between the two groups (Figure 1).

**Figure 1.**
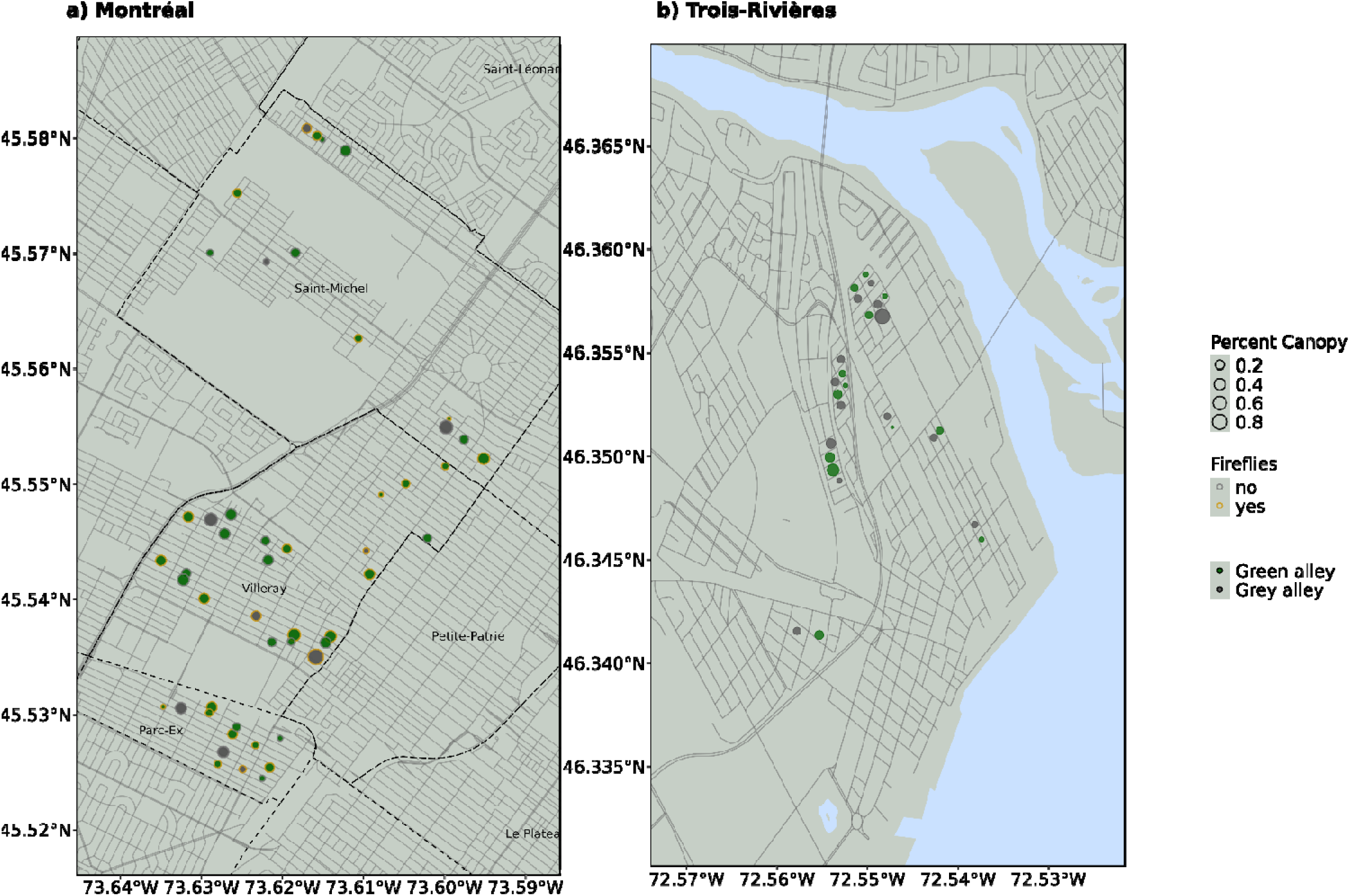
Map representing study areas in a) Montréal and b) Trois-Rivières. Each dot represents an alley included in the study, with green dots representing green alleys and grey dots representing grey alleys. A yellow outline on the dot indicates that we surveyed the alley for fireflies (Montréal only). Dot size corresponds to percent canopy cover of the alley. Light grey lines represent roads, black dashed lines represent neighbourhood boundaries. Figure created using sf, ggplot2, and osmdata packages (Padgham et al. 2023; Pebesma et al. 2024; Wickham et al. 2024).

#### 2.1.2 Trois-Rivières

Trois-Rivières (TR) is a city of ∼ 139,000 people with a population density of 155 residents per km^2^ within the Mauricie region of Quebec, approximately 139 km northeast of Montreal. TR has a mainly white, francophone population and residents with a range of income levels, with high proportions of low-income residents concentrated in the city centre (Statistics Canada and Government of Canada 2022). We sampled all 13 green alleys present in TR and 13 grey alleys adjacent to the green alleys (Figure 1).

#### 2.1.3 Street Segments

In both cities, we sampled the sidewalk and adjacent yards/boulevards parallel to the longest segment of each selected alley (hereafter “street segment”) to facilitate comparison between alleys and streets which are somewhat similar in form but have different function and management regimes. Using satellite imagery, we identified the two potential street segments parallel to the alley, then selected the one which had the most green infrastructure, i.e., segments with street trees, garden squares, and vegetated curb cutouts. If both street segments parallel to the alley had similar levels of green infrastructure, we chose via coin flip (Supplementary Materials A, Table S1).

### 2.2 Biodiversity Support

#### 2.2.1 Biodiversity Support Metric Selection

We use the framework of “nature for nature” to describe biodiversity support. We chose to focus on sampling trees and vegetative complexity, as trees play a key role in ecosystem functioning and biodiversity support (Manning et al. 2006). Additionally, trees are often important to municipal managers and residents alike (Pearce et al. 2015; Martin et al. 2025). Because alleys are small, multiuse areas we selected biodiversity support metrics that were appropriate given that context and which we could collect data on given time constraints. To calculate our first biodiversity support metric, canopy cover, we used existing, open-source data (Institut National de Santé Publique du Québec and Gouvernement du Québec 2022).

#### 2.2.2 Tree Diversity

We sampled trees by identifying and measuring the diameter at breast height (DBH) of every tree (defined as a woody plant > 2 m tall) found at the site. We then assessed tree diversity in two ways. First, we calculated the species richness of each site by counting the number of unique tree species. We recorded a total of 75 species (Supplementary Materials A, Table S2). The second diversity metric was functional group richness. Using the original dataset and methodology from Paquette et al. (2021), we assigned a functional group, i.e., a group of species that share a similar set of biological traits, to each of the tree species present in our dataset and counted the number of functional groups present at each site (Belluau et al. 2021, Supplementary Materials A, Table S2).

#### 2.2.3 Tree Species Composition

We also assessed the proportion of invasive and native trees at each study site by calculating the proportion of stems at each site that were native and invasive. We assigned a status of native or invasive to each of our study species (Philp 2024, Padvaiskas et al. Accepted, Supplementary Materials A, Table S2) according to the native ranges of the St. Lawrence Lowlands ecoregion. We categorized invasive species based on Quebec-specific sources (Plant Dynamics Laboratory and Plant Biology Research Institute 2008; Lavoie and Ayotte 2019; Lavoie et al. 2022).

#### 2.2.4 Vegetative Complexity

We measured vegetative complexity by walking linearly through each study site and taking measures every 10 m. At each stop, we counted how many of the 5 following vegetative layers were present as a proxy for complexity: ground cover, herbaceous vegetation, wall vegetation, shrubs, or canopy. Average vegetative complexity of sites was then calculated by summing the total number of layers counted and then dividing by the number of points measured.

#### 2.2.5 Firefly Survey

To assess the habitat quality of our study alleys, we selected a subset of alleys in VSMPE and performed a presence/absence survey of fireflies (Family: Lampyridae). Fireflies are known bioindicators in cities, as they require leaf litter, access to water, and low levels of light pollution (Picchi et al. 2013; Firefly Atlas et al. 2023). Due to time constraints, we were unable to sample fireflies in TR, however due to the lack of canopy and leaf litter accumulation we don’t believe they would be a strong indicator of biodiversity in the system. We selected the 28 green alleys with the highest canopy cover and most vegetative complexity for our subset, the five grey alleys with the least canopy cover and vegetative complexity as our controls, and the associated street segment for all 34 alleys surveyed. If the presence of a firefly was noted in any of the surveys, the site was assigned as having fireflies present. Additional details on methodology and data sources of the above metrics can be found in the supplementary materials (Supplementary Materials Section B).

### 2.3 Ecosystem Service Indicators

#### 2.3.1 Interviews & Ecosystem Service Indicator Selection

In the summer of 2023, we did 28 semi-structured walking interviews to assess the perceived and desired ecosystem services present in green alleys in VSMPE and TR as part of a larger multi-university project. Interviews were conducted by research professionals (EPM, NB) and graduate students (FB, PF) in urban and environmental studies. All interviewers had experience conducting interviews and interviewed individuals in the language of their choosing, either French or English. Residents were contacted via flyers posted in their neighbourhoods and Facebook posts in community groups. After agreeing to participate, residents met with two interviewers, where they followed a pre-chosen route through green alleys, other nearby greenspaces and adjacent grey alleys. The lead interviewer asked a series of interview questions designed to connect perceptions and experiences of nature to ecological benefits and conditions (Hoyle et al. 2019) while the second documented everything the interviewee mentioned using photos (for interview script see Supplementary Materials B). Although walking interviews resulted in fewer participants than other methods such as surveys, they allowed a more in depth understanding of individual’s preferences and desires (duration varying between 1 and 1.5 hours), described as they were experiencing them in the relevant context (in our case, by walking in the alleys and streets that they know well). Methods were approved by the university’s ethics committee (Permit No. 2023-5054).

Our interviewees (n = 30) covered a diverse range of ages, immigration status, genders, countries of origin, employment status, housing types, and neighbourhoods to capture the variation in resident composition of the study areas (Supplementary Materials A, Table S3).

However, one notable bias was that most of our participants (76%) were highly educated. Multiple team members (FB, PF, EPM, NB) transcribed the interviews and used thematic content analysis coding with NVivo to sort interview contents into broad categories. We then went through all broad categories and inductively coded mentions of desired and perceived ecosystem services made by interviewees. We counted both how many individuals and how many times each interviewee mentioned each ecosystem service across all interviews (Supplementary Materials A, Table S4). Examples of perceived and desired ecosystem services include:

Cultural value of trees: “On the street, the trees are enormous, and I love streets that have a lot of trees” Temperature regulation: “[About the green alley] It’s so much more pleasing to the eye and even helps oxygenate the air and cool the summer too. Yes, I think it will do some good.”
Aesthetic appreciation of flowers: “Well, you don’t often see trees like this in the city. It’s like, you stop and go wow! With the tree next to it with all the little white flowers, I don’t know what it is, but it’s beautiful.”
Using the list of perceived and desired ecosystem services mentioned by interviewees, we selected ecosystem service indicators to quantify that were highly mentioned and overlapped with the ecological data we had previously collected via fieldwork. The final list of ecosystem service indicators selected includes temperature mitigation/cooling, the presence of trees, the presence of large trees, and the presence of flowering trees.

#### 2.3.2 Cooling

To assess the capacity of alleys to provide temperature mitigation, we placed one Kestrel DROP D2 wireless temperature sensor (0.1 resolution ± 0.5 ) in every alley and recorded the temperature every 15 minutes. Due to the highly heterogeneous nature of the alleys, we opted to place the sensors in the greenest part of the alley, to assess the maximum cooling potential. The greenest section was assessed on-site, typically chosen as an area with the highest canopy cover and vegetative complexity. We deployed sensors to all study alleys in Montréal on June 27-28 and retrieved them July 25-26, 2023. We then deployed the same sensors using the same methodology in Trois-Rivières on August 6-8 and collected all sensors on October 3, 2023.

#### 2.3.3 Tree Abundance

We calculated tree abundance by summing the number of individual stems in each of the study sites. We included dead trees in this metric.

#### 2.3.4 Large Trees

We quantified two metrics to capture the presence of large trees. First, we calculated the mean observed tree DBH followed by the mean potential height for each study site. To calculate mean observed DBH, we took the average DBH of all the trees in an alley or street segment. More details on DBH measurements can be found in the supplementary materials (Supplementary Materials B).

To calculate the mean potential height of alley trees, we collected data on the maximum height for individual tree species included in our study (Supplementary Materials A, Table S2) using published data from previous studies (Hutt-Taylor and Ziter, 2022) and other sources (Missouri Botanical Garden, 2024; The Morton Arboretum, 2022; United States Department of Agriculture and Natural Resources Conservation Service, 2024; University of Florida, 2024). We then took the average maximum height of all the trees at a site.

#### 2.3.5 Flowering Trees

We calculated the proportion of flowering trees at each study site by identifying if each study species had showy flowers (Supplementary Materials A, Table S2, Hutt-Taylor and Ziter, 2022; Missouri Botanical Garden, 2024; The Morton Arboretum, 2022; United States Department of Agriculture and Natural Resources Conservation Service, 2024; University of Florida, 2024). We then calculated the proportion of stems that did have showy flowers.

### 2.4 Statistical Analyses

Our statistical approach is based on a causal inference framework (Grace and Irvine 2020; McElreath 2020), which is increasingly called for in ecology (Siegel and Dee 2024). To test the effect of alley type on our various response variables, we constructed directed acyclic graphs (DAGs) representing our assumptions and hypotheses about the causal links between variables in our system (Laubach et al. 2021; Arif and MacNeil 2023). Given the resident-led nature of VSMPE’s green alley program, based on the literature, we hypothesized several confounding relationships between alley type and socio-demographic and economic characteristics of the neighbourhoods (Brazeau-Béliveau and Cloutier 2021; Pham et al. 2022; Forde et al. 2024, Figure 2). Because Trois-Rivières is a municipally led program and green alleys were selected for implementation based exclusively on the state of the pavement, we did not hypothesize any confounding relationships and thus do not present any DAGs. We used DAGs in combination with do-calculus to identify the variables we needed to add to our model to deconfound our system and test the effect of alley type on our response variables in VSMPE (Pearl and Mackenzie 2018). This process resulted in the addition of alley type, income level, and languages spoken to our models (Figure 2). Income level was represented using median after-tax income for the dissemination area and language was represented using the proportion of the population that had knowledge of each of the official languages (i.e., English, French, both, or neither).

**Figure 2.**
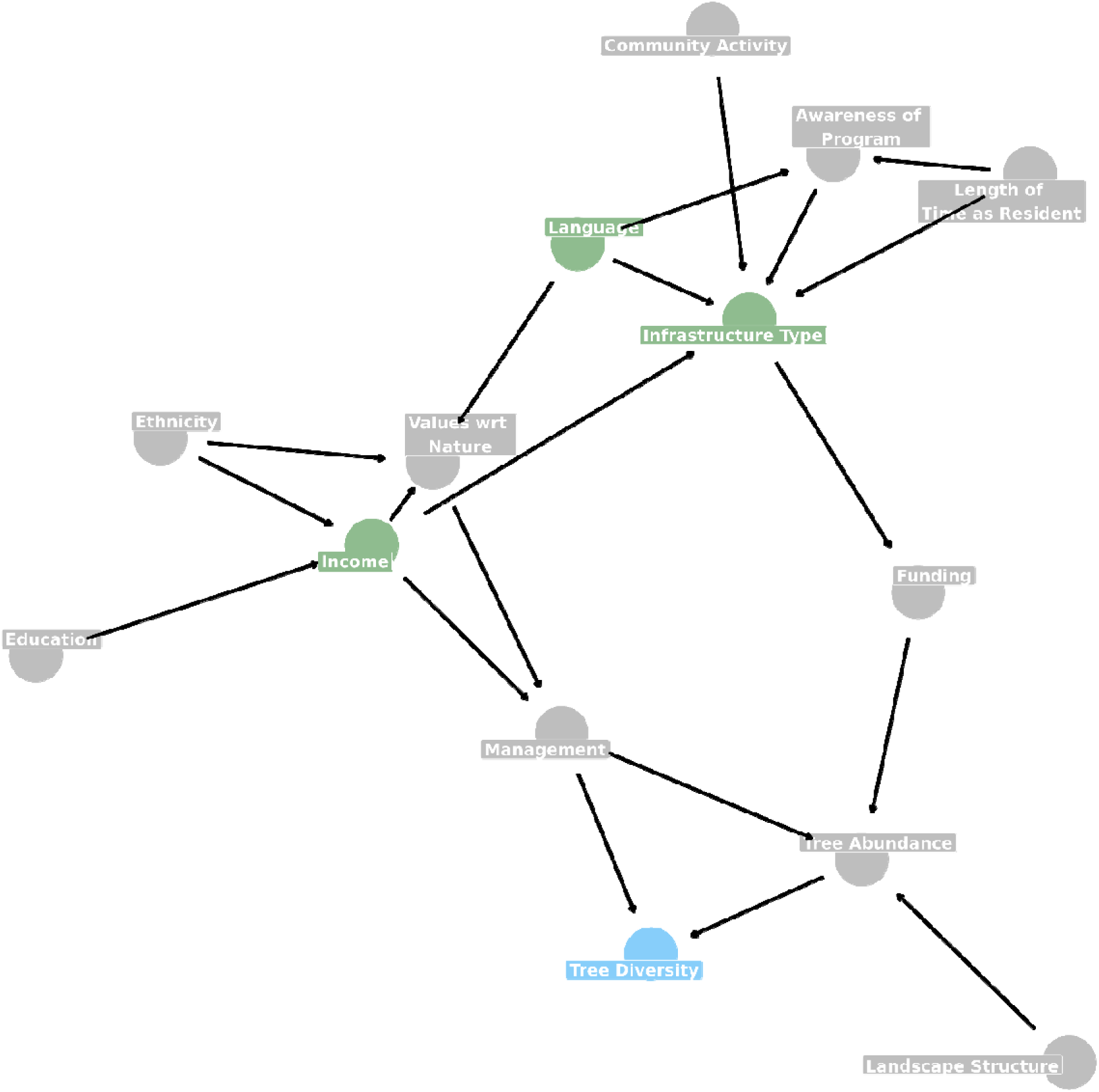
Example of a directed acyclic graph (DAG) depicting our hypothesized and assumed causal relationships between variables in our system for Villeray—Saint-Michel—Parc-Extension’s resident-led green alley program. The variable highlighted in blue is the outcome variable. Infrastructure type is the variable that we want to test the effect of and is in green. Other variables that are required in the model to deconfound our model, i.e., income and language, are highlighted in green. Figure created using ggdag and ggplot2 (Barrett 2024; Wickham et al. 2024).

To test the effect of implementing a green alley on biodiversity support and ecosystem services included in this study, we model each biodiversity support metric (n = 7) and ecosystem service indicator (n = 5) individually using generalized linear models. All biodiversity support metrics and ecosystem service indicators are modelled twice, once for each city (n = 23 models total, firefly abundance is measured only in VSMPE due to data constraints). To control for confounding of the system from the resident-led model in VSMPE (as presented in model-specific DAGs, Supplementary Materials C, Figure S1), we stratified our models by language spoken and median income and added neighbourhood as a varying intercept variable.

Temperature models had a different structure as both date and site had to be included and thus are mixed effects models in both VSMPE and TR. Temperature models included neighbourhood, date, and site ID as varying intercept random effects. Two-way interactions were used to test the relationships between time of day, infrastructure type (i.e., green alley, grey alley, or street segment), and day of year. Full model equations with priors are presented in the supplementary (Supplementary Materials D, Equations 1-23).

To calculate the effect of implementing green alleys on the level of biodiversity support and ecosystem services, we computed the global mean contrasts between infrastructure groups (i.e., green alleys, grey alleys, street segments) using the posterior distributions of each model. For VSMPE models, the adjustment variables (i.e., language spoken and median income) in the model were set to their average values when computing the contrasts. For the temperature model, we graphically represented the contrast instead of computing mean values with credible intervals, due to its complex model structure. We estimated the posterior distributions using Hamiltonian Monte Carlo as implemented in the brms 2.21.0 and cmdstanr 0.8.1 packages (Bürkner et al. 2024; Gabry et al. 2024). We assessed model convergence by inspection of trace plots, R-hat values, and effective sample sizes. Prior predictive simulations as well as model diagnostics are presented in the supplementary (Supplementary Materials E, F) and all results can be replicated using the code available at https://github.com/zule-lab/infrastructuresvertes, archived at https://zenodo.org/records/15304733.

## 3 Results

### 3.1 Summary Statistics

#### 3.1.1 Biodiversity Support

Canopy cover across all sites was on average 15% higher in VSMPE than TR (Table 1). Tree diversity and species composition were higher in VSMPE than TR, following the trends of canopy cover and tree abundance (Table 1). The proportion of native trees and invasive trees was higher in TR than VSMPE (Table 1). Vegetative complexity was slightly higher on average in VSMPE compared to TR (Table 1). Finally, in VSMPE, we recorded a total of 28 firefly sightings across 12 sites.

#### 3.1.2 Ecosystem Service Indicators

Temperature data was collected across different dates for each city, inhibiting direct comparison of raw temperature values. VSMPE had a mean daytime temperature across all sites of 24.4 ± 3.6°C and a mean nighttime temperature across all sites of 21.9 ± 2.5°C. TR, measured two months later, had a mean daytime temperature across all sites of 20.3 ± 4.9°C and a mean nighttime temperature across all sites of 16.3 ± 4.1°C. Tree abundance and size was higher overall in VSMPE when compared to TR (Table 1). Our aesthetic measurement, proportion of trees with showy flowers, had higher levels in VSMPE than TR (Table 1).

### 3.2 Modeling & Contrasts

Overall, results were variable across biodiversity support metrics and ecosystem services, as well as between cities. However, for most of the biodiversity support metrics, green alleys provide at least a slight increase in benefits when compared to grey alleys or street segments (Table 2).

**Table 2.**
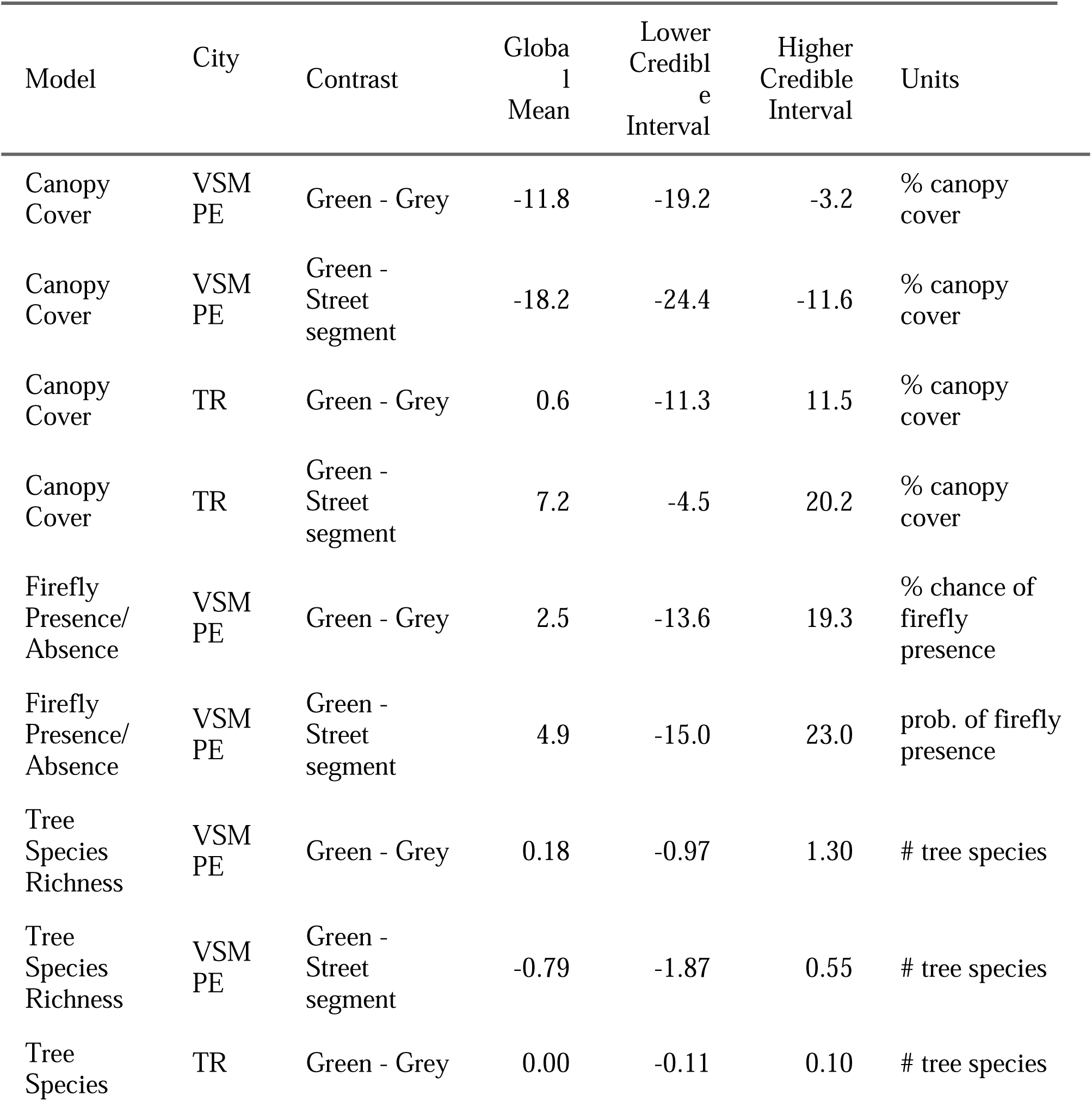
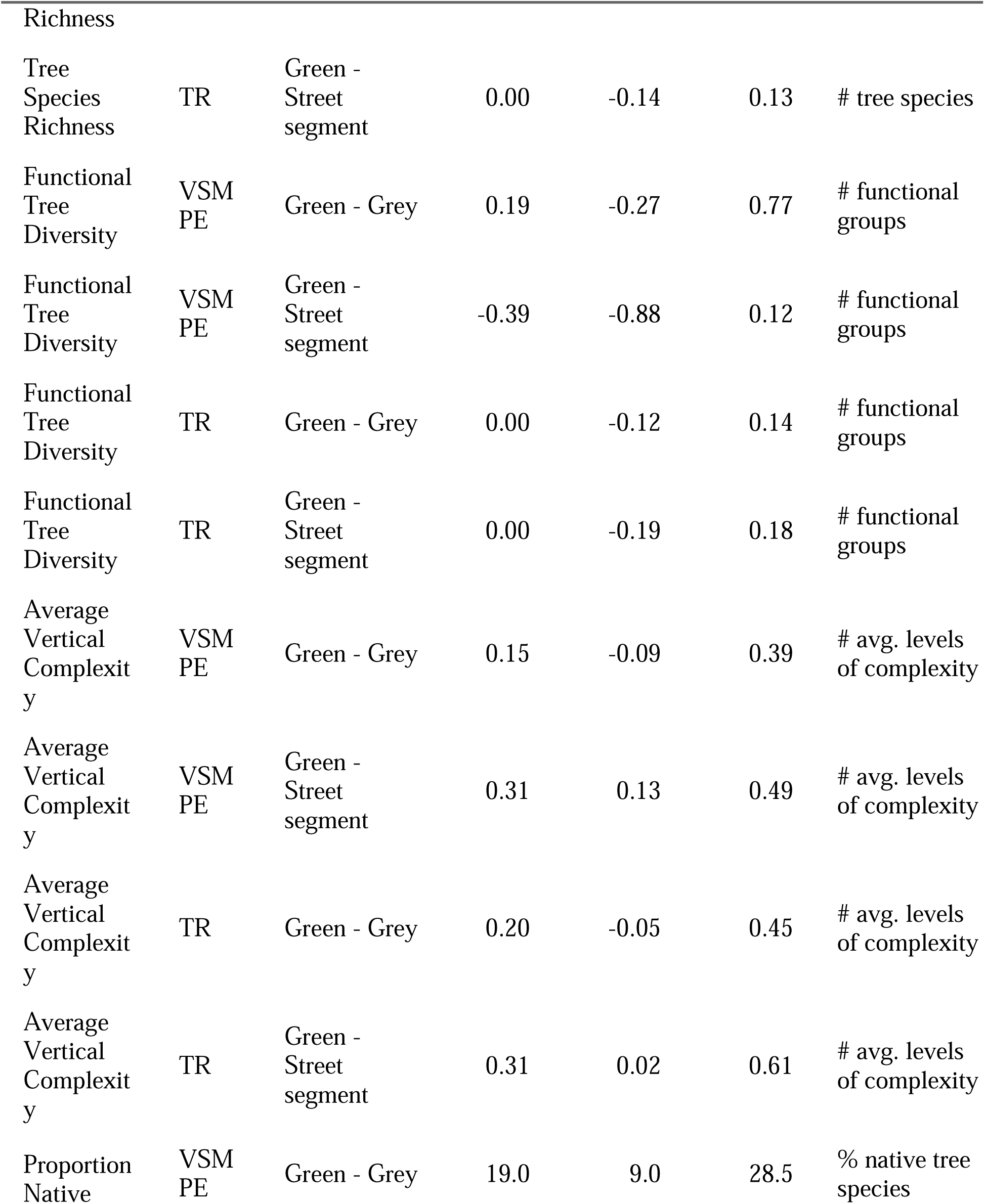
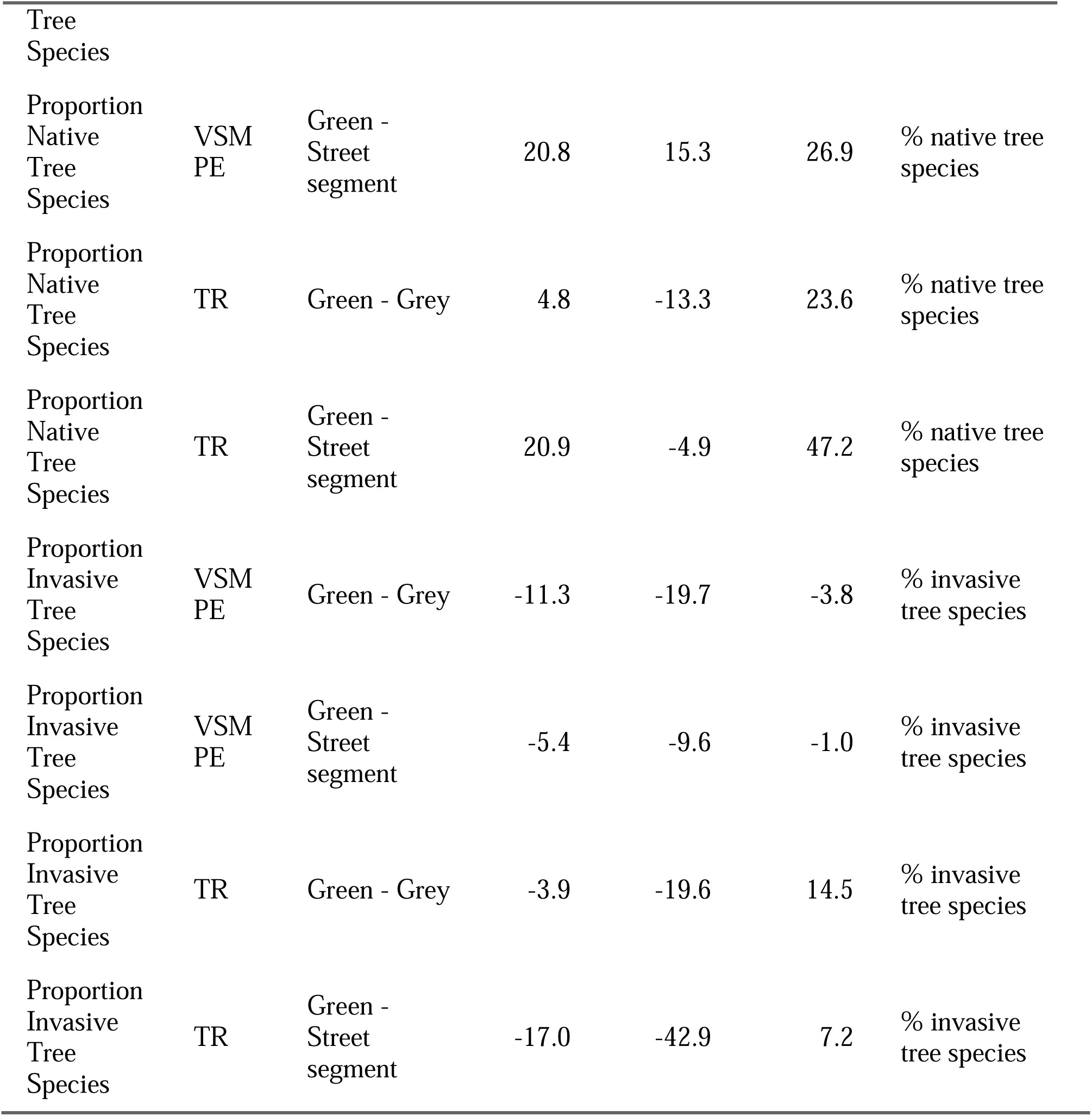
Model contrasts for biodiversity support models. Model name refers to response variable in the model, city refers to the city where the data was collected, contrast indicates the groups contrasted for each row where ‘Green’ refers to a green alley, ‘Grey’ refers to a grey alley, and ‘Street segment’ refers to street segment class, also referred to as street segment, global mean is the mean difference between the groups after a pairwise contrast was performed on the entire posterior distribution of each group, lower credible interval shows the lower 95% credible interval of the contrast, higher credible interval shows the higher 95% credible interval of the contrast, and units represents the output units of the model/contrast. For each contrast, the global mean value presents the average impact of an intervention where you switch from the state on the right (e.g., grey alley) to the state on the left (e.g., green alley) with the units provided.

Ecosystem services were very variable, and we found that their delivery in green alleys does not always align with the interests of the interviewed residents (Table 3).

**Table 3.**
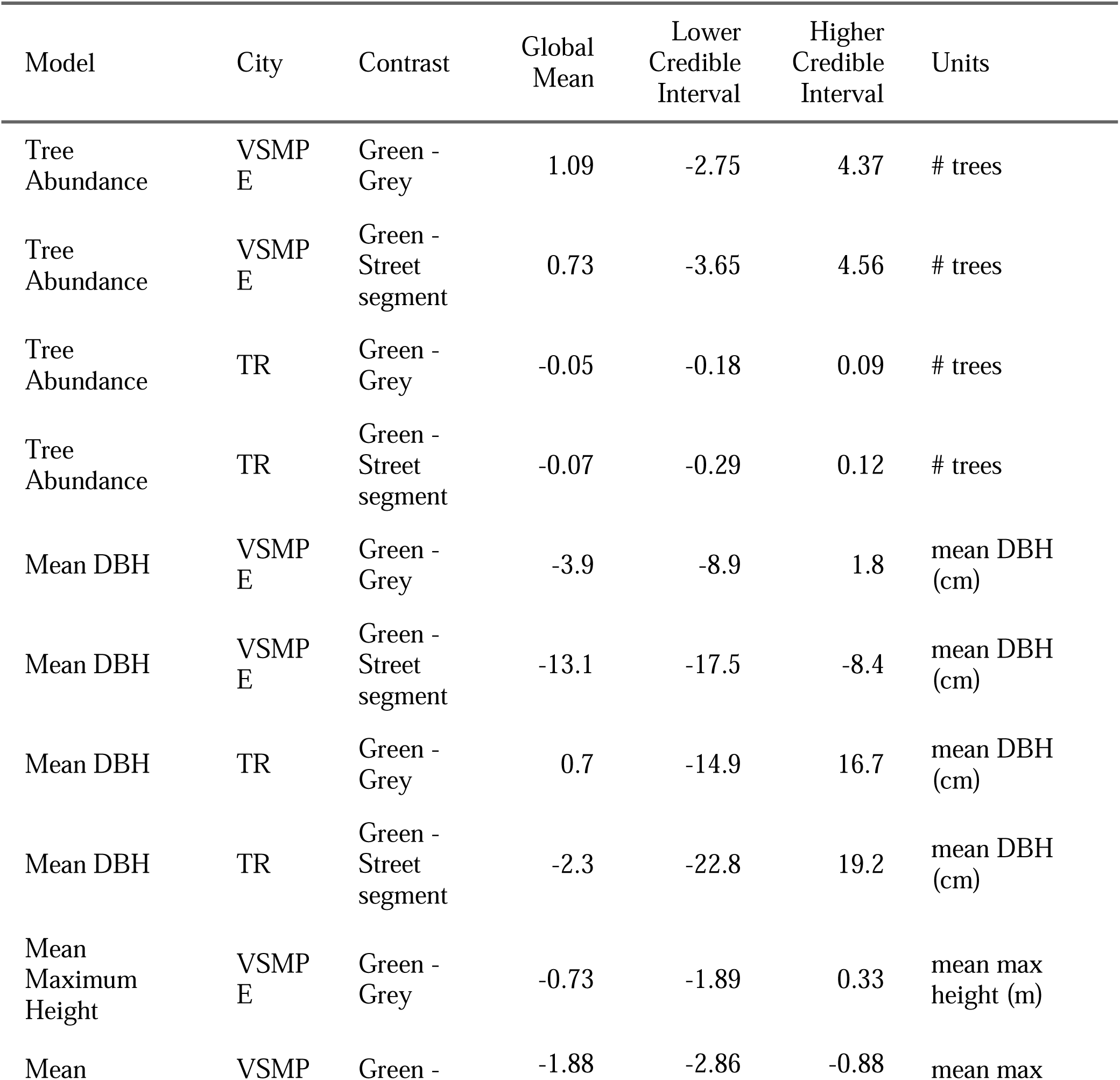
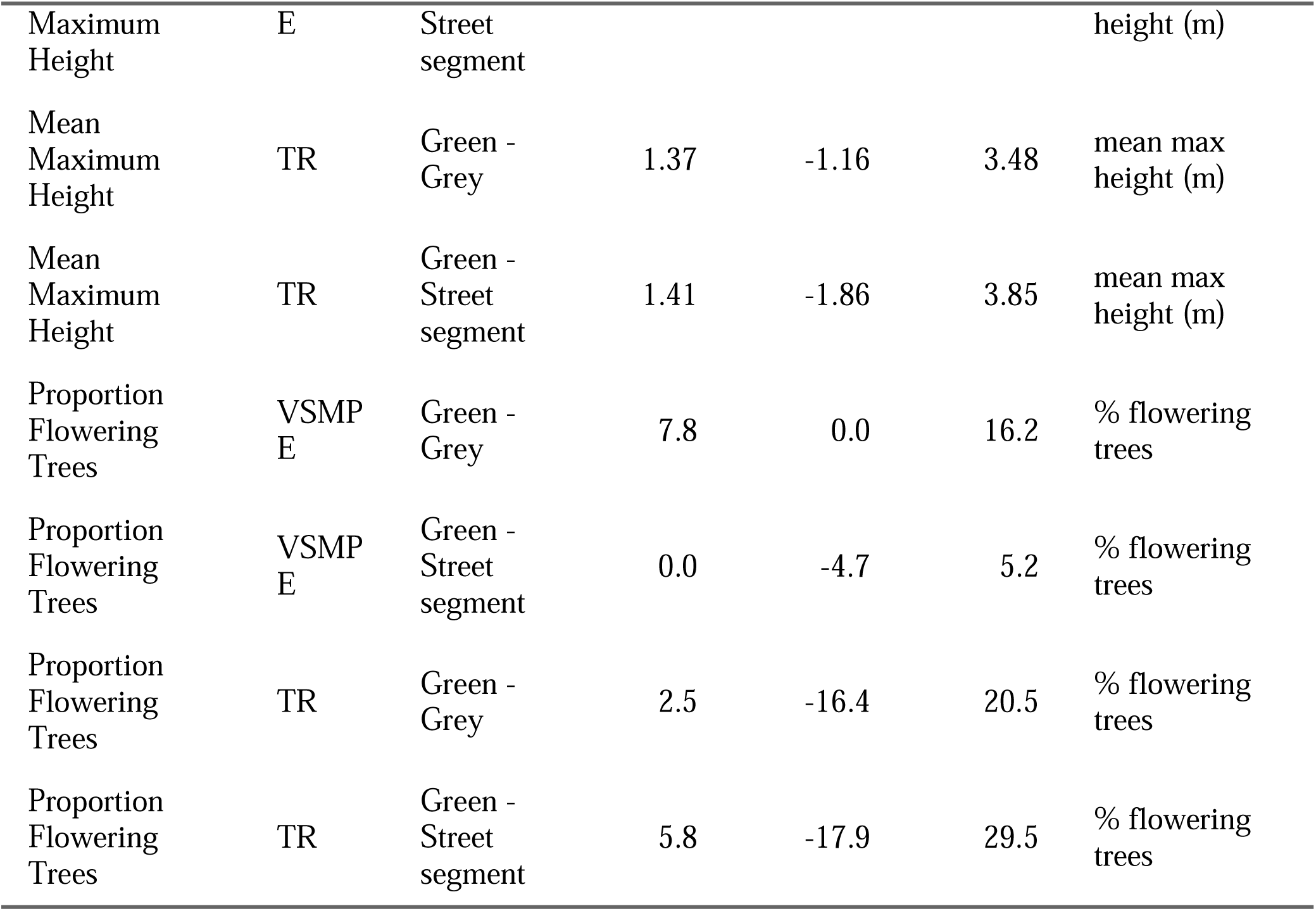
Model contrasts for ecosystem service indicator models. Model name refers to response variable in the model, city refers to the city where the data was collected, contrast indicates the groups contrasted for each row where ‘Green’ refers to a green alley, ‘Grey’ refers to a grey alley, and ‘Street segment’ refers to street segment class, also referred to as street segment, global mean is the mean difference between the groups after a pairwise contrast was performed on the entire posterior distribution of each group, lower credible interval shows the lower 95% credible interval of the contrast, higher credible interval shows the higher 95% credible interval of the contrast, and units represents the output units of the model/contrast. For each contrast, the global mean value presents the average impact of an intervention where you switch from the state on the right (e.g., grey alley) to the state on the left (e.g., green alley) with the units provided.

#### 3.2.1 Biodiversity Support

In VSMPE, where each alley is managed by the local community, we found high levels of variation in our models (Table 2). Biodiversity support was generally higher in green alleys, though by small margins for some variables (Table 2). The exceptions being canopy cover and tree diversity. Canopy cover was consistently lower in green alleys than grey alleys (global mean = - 11.8 %) and street segments (global mean = - 18.2 %). Tree species richness was similar across groups, with contrast means less than one species different. However, at the extreme ends of the 95% credible interval, green alleys could have up to 1 more species than “grey alleys” and almost 2 (1.87) less species than street segments. The same trends held for functional group richness, with differences between green alleys and other infrastructure being less than 1 functional group but general trends indicating that green alleys have slightly higher functional group richness than grey alleys and slightly less functional group richness than street segments. Firefly presence was on average 2 % more likely in green alleyways when compared to grey alleyways, and 4 % more likely in green alleyways than street segments, with high levels of variation across both groups. Vertical complexity was higher in green alleys when compared to “grey alleys” (global mean = 0.15 increase in mean levels of complexity) and even more so when compared to street segments (global mean = 0.31). The percentage of native tree species in green alleys was notably higher than in grey alleys (global mean = 19.0% increase in percentage of native species) and street segments (global mean = 20.8%). Finally, the percentage of invasive species was correspondingly lower in green alleys than in grey alleys (global mean = - 11.3 % decrease in percentage of invasive species) and street segments (global mean = - 5.4%).

In TR, we see similar trends to VSMPE with respect to higher levels of vegetative complexity and native trees, and lower proportions of invasive trees in green alleys (Table 2). However, canopy cover in green alleys is slightly higher than in grey alleys (global mean = 0.7 % increase) and street segments (global mean = 7.2 % increase). There was no difference in tree species richness or functional group richness when comparing green alleys to grey alleys or street segments. There was an increase in average vertical complexity when comparing green alleys to both grey alleys (global mean = 0.20 increase in mean levels of complexity) and street segments (global mean = 0.31). There was a slight increase in the percentage of native trees in green alleys when compared to grey alleys (global mean = 4.8 % increase) and a larger increase when compared to street segments (global mean = 20.9 %). Finally, the proportion of invasive trees decreased slightly when comparing green alleys to grey alleys (global mean = -3.9 % decrease) and decreased notably when comparing green alleys to street segments (global mean = -17.0 %).

#### 3.2.2 Ecosystem Service Indicators

The temperature models were our most complex models, and the results are presented in graphical form instead of being included in the results tables (Figure 3). When looking across the sampling period, temperature was similar between green alleys and grey alleys in both cities, with high variation across both infrastructure types (Figure 3). However, there are some key differences across infrastructure types when broken down by time of day. On average green alleys were cooler during the day than grey alleys (Figure 3). At night, there was no visible trend in VSMPE, with green alleys being both warmer and cooler than grey alleys (Figure 3). In TR, green alleys were noticeably cooler than grey alleys at night, however, they were also noticeably warmer than grey alleys during the day (Figure 3).

**Figure 3.**
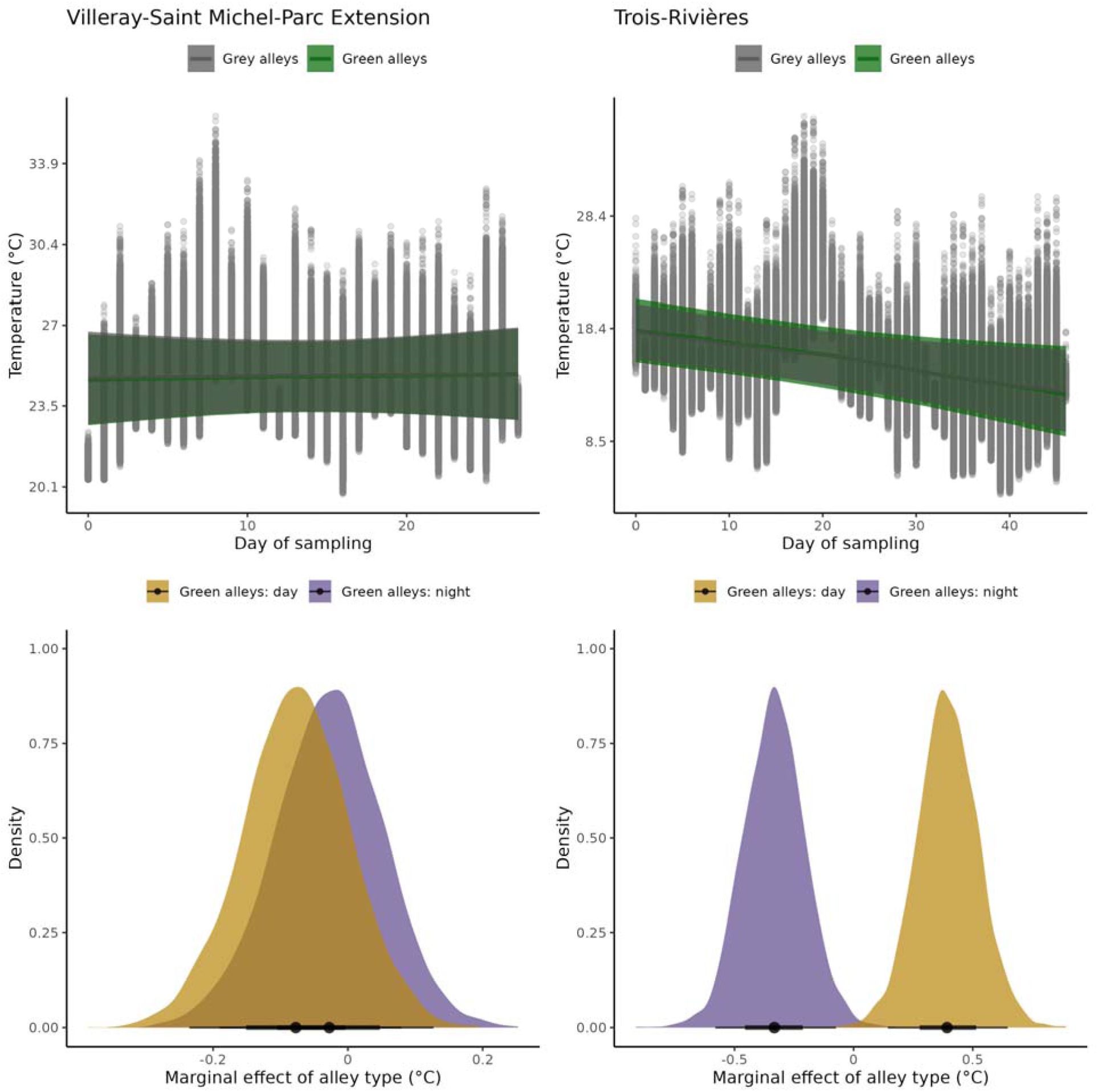
Effects of alley type on temperature. The left column presents the results for Villeray-Saint Michel-Parc Extension (VSMPE) and the right column presents the results for Trois-Rivières. The top row is displaying the model’s fit between the day of sampling on the x-axis and temperature on the y-axis. Each line indicates a different green infrastructure type, with green displaying green alleys and grey displaying grey alleys. The band surrounding the regression line is the 95% credible interval of the model. Each dot on the graph indicates an individual temperature recording from one of the sites. Temperature was measured from June 28 to July 25 in VSMPE and from August 8 to October 3 in Trois-Rivières. The bottom row presents model contrasts for each city, where pairwise comparisons were performed on the posterior distributions. For each contrast, the black dot represents the global mean value, i.e., the average impact of an intervention on temperaturwhen you switch from a grey alley to a green alley during that time of day. The black lines represent the 95% credible interval, and the coloured area is the posterior distribution of the contrast.

The rest of the results for the ecosystem service indicator models are presented in Table 3. These results are more complex and varied than the biodiversity support models. In VSMPE, tree abundance was higher in green alleys than grey alleys (global mean = 1 more tree in green alleys but up to 4 at the high end of the 95% credible interval) and slightly higher than street segments (global mean = 0.7). The average tree size tended to be smaller in green alleys than grey alleys (global mean = -3.9 cm) and much smaller when compared to street segments (global mean = - 13.1 cm). Similarly, the mean maximum (or potential) height of trees in green alleys was smaller than that in grey alleys (global mean = -0.73 m) and street segments (global mean = -1.88 m).

Finally, the proportion of trees with showy flowers was slightly higher in green alleys when compared to grey alleys (global mean = 7.8 % increase) and was the same as what was found in street segments.

In TR, trends are slightly different (Table 3). Tree abundance was at a similar level although slightly lower in green alleys when compared to grey alleys (global mean = -0.05 trees) and street segments (global mean = -0.07 trees). However, the average tree size was higher in green alleys compared to grey alleys (global mean = 0.66 cm larger). When compared to street segments, average tree size was lower in green alleys (global mean = -2.27 cm smaller), similar to VSMPE. The average maximum height of trees in green alleys was on average higher than in grey alleys (global mean = 1.37 m higher) and street segments (global mean = 1.41 m higher). Finally, the proportion of flowering trees was similar across all infrastructure types, but green alleys had on average slightly higher proportions than both grey alleys (global mean = 2.5 % increase) and street segments (global mean = 5.8 % increase).

## 4 Discussion

Our hypothesis that green alleys would have higher levels of both biodiversity support and ecosystem services when compared to grey alleys and street segments was only partially supported. Green alleys did provide either similar or higher levels of biodiversity support, i.e., nature for nature, when compared to grey alleys and street segments, except for canopy cover in VSMPE (Montreal). However, ecosystem service indicators, i.e., nature for people, did not follow the same trend. In general, the ecosystem services desired by local residents were of similar levels or lower in green alleys compared to grey alleys and street segments, except for tree abundance in Montreal which was higher in green alleys. We also predicted that due to the resident-led program design of green alleys in Montreal, the level of biodiversity support and ecosystem services would be more variable than in Trois-Rivières. For most of the 11 indices, this was not true. Both Montreal and TR had variable levels of biodiversity support and ecosystem service indicators. However, for both temperature regulation and tree abundance, Montreal was more variable than TR, as predicted.

We found that Montreal’s green alleys have lower levels of canopy cover than observed in grey alleys and street segments. Initially, this finding may be counterintuitive. Since the green alley program in Montreal prioritizes and funds planting trees (Ville de Montréal 2024), and the presence of large trees was listed as an important ecosystem service indicator by residents (Supplementary Materials A, Table S4), it is reasonable to expect that canopy cover would be higher in green alleys than grey alleys. However, significantly increasing canopy cover in an alley is difficult without large scale intervention. Alleys are narrow spaces, averaging approximately 4 m wide, and usually serve a variety of purposes. They are spaces for play and neighbourhood gatherings while also still sometimes used for vehicle access for backyard parking spots and parking for maintenance vehicles and construction equipment (Wolch et al. 2010; Newell et al. 2013; Pham et al. 2022). Due to the design of alleyways, tree planting typically requires the removal of pavement, for which is resource and effort-intensive and can be restricted by city bylaws, participant energy, and desire to get along with other residents (Brazeau-Béliveau and Cloutier 2021). Additionally, tree planting and demineralization can be controversial as it can hinder or block car circulation. Often, the small-scale greening interventions that residents make in green alleys include planter boxes and narrow bands of vegetation along the side of the alley, which do not provide the space required for trees with large canopies (Pham et al. 2022). By contrast, streets have many of the same constraints as alleys but often receive large scale public interventions that increase canopy cover such as tree pits (Grey et al. 2018). Consequently, most of the tree cover in green alleys, and residential areas in general, is provided by trees on private land (Pataki et al. 2013). Under the given conditions, green alleys having lower levels of canopy can potentially be explained by resident motivation. Residents with less access to private green space and associated canopy cover may be more motivated to participate in a green alley program. By contrast, in Trois-Rivieres the municipality chooses the green alley locations and thus the same mechanism does not apply. Future studies that can assess differences in resident perception, motivation, and ecological benefits before and after the development of a green alley instead of comparing different alleys that have already been built may provide an opportunity to gain a more detailed insight into the unexpected levels of canopy cover. Given that canopy and tree cover was mentioned by interviewees in our study as a desirable aspect of green infrastructure, understanding why it may be lower than expected in green alleys and incorporating solutions such as providing expert support to residents designing green alleys and/or using funds specifically designed for greening purposes to increase canopy cover in green alleys moving forward should be a priority (Brazeau-Béliveau and Cloutier 2021; Ville de Québec 2025).

An important implication of canopy cover, and vegetation in general, is cooling (Ziter et al. 2019). Cooling and shade were mentioned by residents frequently as valuable ecosystem services (Table S4). Southern Quebec reaches high temperatures in the summer, with the urban heat island exacerbating the summer heat levels contributing to heat-related illness and mortality (Boudreault et al. 2024). Given that green alleys are integrated into the urban matrix and have the potential to provide cooling near residences, they can play an important role in combating heat-stress and other heat-related illnesses. Our temperature analysis shows different cooling trends across the two cities. In Montreal, green alleys provide a minimal cooling effect of 0.07°C on average during the day and a negligible cooling effect at night at 0.02°C on average, with some alleys showing the potential to cool more (up to 0.3°C). In contrast, green alleys in Trois-Rivières are providing a stronger cooling effect at night than those in Montreal, with a nighttime cooling average of 0.3°C. During the day in TR, green alleys are actually warmer than grey alleys by an average of 0.4°C. Notably, we measured temperature in these cities at different times, with our measurements in TR occurring in late summer, which may have decreased our ability to capture a stronger cooling effect. The opposing day-night cooling patterns between the two cities highlight the differing management strategies across the two cities. Alleys in Montreal have on average 13 trees and are mostly paved, whereas alleys in Trois-Rivières typically have no trees but have very little impervious surface with the entire alley covered in permeable pavement planted with turfgrass (Supplementary Materials C Figure S2). Open, pervious surfaces such as those found in green alleys in TR have been shown to cool quickly at night, because there is no canopy to trap the heat and no impervious surface to radiate heat (Wujeska-Klause and Pfautsch 2020). By contrast, Montreal’s green alleys have impervious surfaces with higher levels of trees and canopy cover, resulting in slightly lower daytime temperatures but little to no effect at night.

Night is a critical time to provide cooling in cities because it is when the human body recovers from heat stress, and the urban heat island effect can be up to three times stronger at night (Sarofim et al. 2016; Iungman et al. 2023). The differences in impervious surfaces and tree cover across Montreal and TR can be attributed to different goals. TR’s green alley program was focused on improving stormwater management, whereas Montreal has tree planting listed as one of the goals of the green alley program (Ville de Trois-Rivières 2023; Ville de Montréal 2024). Given that green alleys serve many purposes and have different goals, not just cooling, managing the ideal green alley may incorporate a mixed approach of tree planting to increase canopy in tandem with replacing impervious surfaces and maintaining open spaces to meet the needs and wants of residents. Additionally, assessing cooling in such a small and varied area is a challenge, and green alleys may have more of an effect on cooling if they are treated as a network, using a mixed approach to promote and assess cooling alongside other benefits at a neighbourhood scale.

Green alleys in both cities had notably higher levels of native species and lower levels of invasive species compared to both grey alleys and sidewalks, which is an important form of biodiversity support. Invasive and non-native species are a problem in cities, contributing to the decline of native species, overall ecosystem homogenization, and sometimes escaping into habitat outside the city limits (Hardberger et al. 2025). Native plant species have often been considered less desirable by residents as they don’t always have the same aesthetic features of non-natives (Hardberger et al. 2025), but our results show that green alleys may present an opportunity for native plant establishment. The dominance of native trees in green alleys also likely provides more resources for pollinators and birds (Garland and Wells 2020), which were two additional ecosystem services desired by residents. Understanding the mechanism behind the establishment of higher levels of native plants in green alleys could help managers and residents continue this trend in other urban green spaces such as private yards and parks to support other forms of biodiversity such as native insects and birds.

A multifaceted approach to green infrastructure management is necessary if we want to provide both biodiversity support and ecosystem services. Our results show that although green alleys generally increase biodiversity support, they do not increase capacity for the ecosystem services desired or perceived by residents. However, there are many potential synergies between these two categories that can be managed for. For example, appreciating the odour and presence of flowers and flowering trees was an ecosystem service mentioned by residents as desirable (Supplementary Materials A, Table S4). Planting native flowering trees can increase tree abundance, provide resources for pollinators and birds, decrease the proportion of invasive species, while increasing the biodiversity of the area (Garland and Wells 2020). The presence of flowering trees was one of the only ecosystem service indicators that was consistently provided at a higher level by green alleys. This is a success for both biodiversity support and ecosystem services which can be highlighted and encouraged in the future.

We also found many synergies between the ecosystem services that residents desire, and attributes commonly considered part of a healthy functioning ecosystem. For example, the stated desire for large trees and habitat provision by residents are both features that support ecological functioning and biodiversity. Notably, most of our interview participants were highly educated, which may indicate that they have more knowledge about ecosystem functioning and biodiversity, potentially resulting in a list of ecosystem services that are unusually synergistic with support for biodiversity (Wojewódzka-Wiewiórska et al. 2022). Repeating more studies that include people with a range of education levels to see if this pattern holds would be beneficial.

Also, many of the ecosystem services that people are seeking, and which would improve quality of life for residents, may require more funding, continued management, and assistance than is currently being provided. For example, providing ongoing, expert support that supports the vision of the residents can facilitate green alleys having a more inclusive and equitable approach, as well as a socially and ecologically successful output (Brazeau-Béliveau and Cloutier 2021).

Notably, many of the stated goals or desires of residents for green alleys can be achieved through pathways that are not tree related. This study focused on the biodiversity support and ecosystem services provided by trees, but there were ecosystem services listed by residents as important – such as overall “greenness”, food provisioning, and habitat provisioning for fauna (Table S4)– that would be better captured by assessing non-tree vegetation. We focused on trees as the city of Montreal includes tree planting as a desired outcome of green alleys (Ville de Montréal 2024), trees perform a disproportionate role in ecosystem services such as cooling (Schwaab et al. 2021), and the presence of trees was highly desired by interviewees (Supplementary Materials A, Table S4). This was a limitation of our research in Trois-Rivieres especially, as the green alleys prioritized permeable pavement and had very few trees. There is a need for future research that focuses on the biodiversity support and ecosystem services provided by other types of vegetation in green alleys, such as herbaceous plants, and faunal surveys for pollinators, birds, and other taxa of interest.

Our study advances our knowledge on monitoring a poorly understood type of green infrastructure, green alleys. Although there have been some surveys of the biodiversity support and ecosystem services provided by green alleys (Coseo and Larsen 2015; Dade et al. 2024), they are nowhere near as comprehensively studied as other types of green infrastructure such as urban parks or green roofs. Green alleys are difficult to study, as they have a very unique form compared to other types of green infrastructure typically surveyed by ecologists, and vegetation and management can be highly variable even within the same alley (Dade et al. 2024). In addition to the highly variable structure within and across alleys, the goals of green alleys can be very different from one to the next, especially if they are managed by neighbourhood residents. The high levels of social and ecological variability can make it difficult to determine what variables are relevant or meaningful to measure. We believe that using interdisciplinary approaches, such as combining resident interviews and knowledge with ecological fieldwork as we did for this study, can help provide a framework for social-ecological work that results in meaningful indicators and biodiversity studies (van Oudenhoven et al. 2018; Schröter et al. 2021). Using a combination of our knowledge as ecologists, social scientists, and local experts, i.e., residents, we were able to conduct surveys that were relevant for both human and non-human residents of alleys. Using this work, we can continue to build and expand our surveys so that we have a more complete picture of what benefits green alleys are providing in our cities and build better green alleys that serves both other forms of nature and people in the future.

## 5 Conclusion

Green alleys are an increasingly popular type of green infrastructure in cities, with potential to provide benefits for nature and people. We found that green alleys are indeed capable of providing both biodiversity support and ecosystem services, although they do not always do so and currently provide relatively more biodiversity support. There were differences in biodiversity support and ecosystem service provision by green alleys between the two cities, which have different management regimes. The community-based management of green alleys in Montreal resulted in higher variation in temperature and tree abundance. The municipal-based “one size fits all” approach of Trois-Rivières resulted in less variation and more nighttime cooling, but also a lower abundance of trees, which were listed as an important provider of ecosystem services by residents. Designing green alleys to meet the needs of both human and non-human residents requires asking people what they need, allowing communities to lead the design and implementation of these green infrastructure projects, and providing technical and expert support to allow communities to reach their goals.

## Acknowledgements

Thank you to Niraj Dayanandan, Adela Bartova, and Dr. Emma Despland for help with fieldwork and methodological advice. Thank you to Dr. Danielle Dagenais, Paul Emile Tchinda, Patrick Boivin, Eva Doan-Lavoie, Khac Minh Tran, Kelly Vu, and Lisa Abou Rjeily for help with project conceptualization and organization. Thank you to members of the Ziter Urban Landscape Ecology lab for helpful revisions on earlier versions of this manuscript. Thank you to Dr. Andrew MacDonald for statistical support. Finally, thank you to alley residents and interviewees for helping us understand your perspectives on green alleys.

