## Supplementary Materials for "Green alleys in Quebec provide variable biodiversity support and ecosystem services"

**References** **96**

#### A) Tables

Table S1. Site characteristics. Infrastructure ID is the unique ID assigned to each individual site, where CON indicates a grey alley, RV indicates a green alley, and SS indicates a street segment. Parallel street names one of the parallel streets to each alley, and for street segments it indicates the street segment that was sampled for each alley. City is where the sampling occurred, VSMPE = Villeray-Saint Michel-Parc Extension and TR = Trois-Rivières. Infrastructure type indicates the site type, with three potential options, green alley, grey alley, and street segment. Percent canopy indicates the percent canopy of the site in decimal form. Firefly presence is a binomial variable, where 1 indicates that fireflies were found, 0 indicates no fireflies were found, and a blank indicates that the site was not sampled for fireflies. Number of points is the number of points where vegetative complexity was sampled (i.e., number of 10 m intervals).

| Infrastructure ID | Parallel Street | City | Infrastructure Type | Percent Canopy | Firefly Presence | Number of Points |
|---|---|---|---|---|---|---|
| CON-SS-TR-10 | Ste Angèle | TR | street segment | 0.39 | | 17 |

| Infrastructure ID | Parallel Street | City | Infrastructure Type | Percent Canopy | Firefly Presence | Number of Points |
|---|---|---|---|---|---|---|
| CON-SS-VSMPE-1 | d'Outremont | VSMPE | street segment | 0.06 | 0 | 15 |
| CON-SS-VSMPE-2 | d'Outremont | VSMPE | street segment | 0.35 | | 31 |
| CON-SS-VSMPE-3 | de l'Epée | VSMPE | street segment | 0.53 | | 25 |
| CON-SS-VSMPE-4 | 51e | VSMPE | street segment | 0.12 | 0 | 9 |
| CON-SS-VSMPE-5 | 9e | VSMPE | street segment | 0.41 | | 52 |
| CON-SS-VSMPE-6 | 2e | VSMPE | street segment | 0.61 | | 21 |
| CON-SS-VSMPE-7 | Casgrain | VSMPE | street segment | 0.54 | 0 | 17 |
| CON-SS-VSMPE-8 | Chambord | VSMPE | street segment | 0.65 | 1 | 15 |
| CON-SS-VSMPE-9 | de Chateaubriand | VSMPE | street segment | 0.61 | | 17 |
| CON-TR-10 | Ste Angèle | TR | grey alley | 0.12 | | 21 |
| CON-VSMPE-1 | d'Outremont | VSMPE | grey alley | 0.10 | 0 | 20 |
| CON-VSMPE-2 | d'Outremont | VSMPE | grey alley | 0.39 | | 37 |
| CON-VSMPE-3 | de l'Epée | VSMPE | grey alley | 0.38 | | 26 |
| CON-VSMPE-5 | 9e | VSMPE | grey alley | 0.08 | | 47 |
| CON-VSMPE-6 | 2e | VSMPE | grey alley | 0.60 | | 24 |

| Infrastructure ID | Parallel Street | City | Infrastructure Type | Percent Canopy | Firefly Presence | Number of Points |
|---|---|---|---|---|---|---|
| CON-VSMPE-7 | Casgrain | VSMPE | grey alley | 0.90 | 0 | 21 |
| CON-VSMPE-9 | de Chateaubriand | VSMPE | grey alley | 0.54 | | 17 |
| RV-SS-TR-11 | Gingras | TR | green alley | 0.37 | | 6 |
| RV-TR-10 | St Paul | TR | green alley | 0.13 | | 21 |
| RV-VSMPE-10 | Durocher | VSMPE | green alley | 0.25 | 0 | 28 |
| RV-VSMPE-11 | 2e | VSMPE | green alley | 0.07 | | 24 |
| RV-VSMPE-12 | Louis-Hébert | VSMPE | green alley | 0.19 | | 28 |
| RV-VSMPE-13 | 13e | VSMPE | green alley | 0.21 | 0 | 23 |
| RV-VSMPE-14 | 1re | VSMPE | green alley | 0.17 | | 6 |
| RV-VSMPE-15 | 6e | VSMPE | green alley | 0.21 | | 25 |
| RV-VSMPE-16 | 9e | VSMPE | green alley | 0.31 | 0 | 27 |
| RV-VSMPE-17 | 2e | VSMPE | green alley | 0.15 | | 26 |

| Infrastructure ID | Parallel Street | City | Infrastructure Type | Percent Canopy | Firefly Presence | Number of Points |
|---|---|---|---|---|---|---|
| RV-VSMPE-18 | 47e | VSMPE | green alley | 0.38 | | 9 |
| RV-VSMPE-19 | Bressani | VSMPE | green alley | 0.31 | | 35 |
| RV-VSMPE-2 | Durocher | VSMPE | green alley | 0.04 | 1 | 26 |
| RV-VSMPE-20 | 48e | VSMPE | green alley | 0.11 | 0 | 9 |
| RV-VSMPE-21 | 44e | VSMPE | green alley | 0.30 | 0 | 11 |
| RV-VSMPE-22 | des Belges | VSMPE | green alley | 0.40 | 0 | 18 |
| RV-VSMPE-23 | Drolet | VSMPE | green alley | 0.24 | 0 | 25 |
| RV-VSMPE-24 | Papineau | VSMPE | green alley | 0.24 | | 31 |
| RV-VSMPE-25 | St Andre | VSMPE | green alley | 0.30 | 0 | 17 |
| RV-VSMPE-26 | Boyer | VSMPE | green alley | 0.50 | 0 | 25 |
| RV-VSMPE-28 | Drolet | VSMPE | green alley | 0.11 | | 30 |

| Infrastructure ID | Parallel Street | City | Infrastructure Type | Percent Canopy | Firefly Presence | Number of Points |
|---|---|---|---|---|---|---|
| RV-VSMPE-29 | de Lorimier | VSMPE | green alley | 0.40 | 1 | 26 |
| RV-VSMPE-3 | de l'Epée | VSMPE | green alley | 0.32 | 1 | 16 |
| RV-VSMPE-30 | St Andre | VSMPE | green alley | 0.04 | 0 | 13 |
| RV-VSMPE-31 | de Chateaubriand | VSMPE | green alley | 0.04 | 0 | 10 |
| RV-VSMPE-32 | de Normanville | VSMPE | green alley | 0.13 | 0 | 25 |
| RV-VSMPE-33 | Drolet | VSMPE | green alley | 0.10 | | 24 |
| RV-VSMPE-34 | Casgrain | VSMPE | green alley | 0.06 | | 26 |
| RV-VSMPE-36 | de Gaspé | VSMPE | green alley | 0.19 | 0 | 25 |
| RV-VSMPE-37 | Casgrain | VSMPE | green alley | 0.35 | | 7 |
| RV-VSMPE-38 | Chabot | VSMPE | green alley | 0.35 | | 25 |
| RV-VSMPE-39 | Henri-Julien | VSMPE | green alley | 0.31 | 0 | 21 |

| Infrastructure ID | Parallel Street | City | Infrastructure Type | Percent Canopy | Firefly Presence | Number of Points |
|---|---|---|---|---|---|---|
| RV-VSMPE-4 | Stuart | VSMPE | green alley | 0.35 | | 34 |
| RV-VSMPE-40 | des Belges | VSMPE | green alley | 0.12 | 0 | 25 |
| RV-VSMPE-6 | de l'Epée | VSMPE | green alley | 0.10 | 1 | 33 |
| RV-VSMPE-7 | d'Outremont | VSMPE | green alley | 0.18 | | 29 |
| RV-VSMPE-8 | Durocher | VSMPE | green alley | 0.10 | 0 | 32 |
| RV-VSMPE-9 | Bloomfield | VSMPE | green alley | 0.06 | 1 | 35 |
| SS-VSMPE-1 | de l'Epée | VSMPE | street segment | 0.70 | | 24 |
| SS-VSMPE-10 | Durocher | VSMPE | street segment | 0.19 | 0 | 23 |
| SS-VSMPE-11 | 2e | VSMPE | street segment | 0.42 | | 20 |
| SS-VSMPE-12 | Louis-Hébert | VSMPE | street segment | 0.59 | | 20 |
| SS-VSMPE-13 | 13e | VSMPE | street segment | 0.16 | 0 | 14 |
| SS-VSMPE-14 | 1re | VSMPE | street segment | 0.62 | | 6 |

| Infrastructure ID | Parallel Street | City | Infrastructure Type | Percent Canopy | Firefly Presence | Number of Points |
|---|---|---|---|---|---|---|
| SS-VSMPE-15 | 6e | VSMPE | street segment | 0.49 | | 26 |
| SS-VSMPE-16 | 9e | VSMPE | street segment | 0.10 | 0 | 27 |
| SS-VSMPE-18 | 47e | VSMPE | street segment | 0.35 | | 17 |
| SS-VSMPE-19 | Bressani | VSMPE | street segment | 0.39 | | 33 |
| SS-VSMPE-2 | Durocher | VSMPE | street segment | 0.16 | 0 | 26 |
| SS-VSMPE-20 | 48e | VSMPE | street segment | 0.01 | 1 | 9 |
| SS-VSMPE-22 | des Belges | VSMPE | street segment | 0.66 | 0 | 18 |
| SS-VSMPE-23 | Drolet | VSMPE | street segment | 0.58 | 0 | 25 |
| SS-VSMPE-24 | Papineau | VSMPE | street segment | 0.53 | | 26 |
| SS-VSMPE-25 | St Andre | VSMPE | street segment | 0.48 | 0 | 18 |
| SS-VSMPE-26 | Boyer | VSMPE | street segment | 0.59 | 0 | 21 |

| Infrastructure ID | Parallel Street | City | Infrastructure Type | Percent Canopy | Firefly Presence | Number of Points |
|---|---|---|---|---|---|---|
| SS-VSMPE-27 | Saint Dominique | VSMPE | street segment | 0.36 | | 23 |
| SS-VSMPE-28 | Drolet | VSMPE | street segment | 0.55 | | 27 |
| SS-VSMPE-29 | de Lorimier | VSMPE | street segment | 0.22 | 0 | 20 |
| SS-VSMPE-3 | de l'Epée | VSMPE | street segment | 0.30 | 0 | 14 |
| SS-VSMPE-30 | St Andre | VSMPE | street segment | 0.66 | 1 | 18 |
| SS-VSMPE-31 | de Chateaubriand | VSMPE | street segment | 0.62 | 0 | 7 |
| SS-VSMPE-32 | de Normanville | VSMPE | street segment | 0.86 | 0 | 16 |
| SS-VSMPE-33 | Drolet | VSMPE | street segment | 0.57 | | 22 |
| SS-VSMPE-34 | Casgrain | VSMPE | street segment | 0.63 | | 25 |
| SS-VSMPE-35 | de Gaspé | VSMPE | street segment | 0.43 | 1 | 8 |
| SS-VSMPE-36 | de Gaspé | VSMPE | street segment | 0.74 | 0 | 25 |

| Infrastructure ID | Parallel Street | City | Infrastructure Type | Percent Canopy | Firefly Presence | Number of Points |
|---|---|---|---|---|---|---|
| SS-VSMPE-37 | Casgrain | VSMPE | street segment | 0.66 | | 7 |
| SS-VSMPE-38 | Chabot | VSMPE | street segment | 0.56 | | 26 |
| SS-VSMPE-39 | Henri-Julien | VSMPE | street segment | 0.51 | 0 | 21 |
| SS-VSMPE-4 | Stuart | VSMPE | street segment | 0.31 | | 31 |
| SS-VSMPE-40 | des Belges | VSMPE | street segment | 0.55 | 0 | 22 |
| SS-VSMPE-5 | Hutchison | VSMPE | street segment | 0.19 | | 31 |
| SS-VSMPE-6 | de l'Epée | VSMPE | street segment | 0.31 | 0 | 30 |
| SS-VSMPE-7 | d'Outremont | VSMPE | street segment | 0.03 | | 26 |
| SS-VSMPE-8 | Durocher | VSMPE | street segment | 0.19 | 0 | 31 |
| SS-VSMPE-9 | Bloomfield | VSMPE | street segment | 0.33 | 0 | 28 |
| CON-TR-1 | Saint-Louis | TR | grey alley | 0.06 | | 7 |
| CON-TR-11 | Ste Angèle | TR | grey alley | 0.10 | | 9 |
| CON-TR-12 | Williams | TR | grey alley | 0.16 | | 8 |

| Infrastructure ID | Parallel Street | City | Infrastructure Type | Percent Canopy | Firefly Presence | Number of Points |
|---|---|---|---|---|---|---|
| CON-TR-13 | Amherst | TR | grey alley | 0.18 | | 4 |
| CON-TR-2 | Saint-Louis | TR | grey alley | 0.30 | | 7 |
| CON-TR-3 | Dumoulin | TR | grey alley | 0.15 | | 26 |
| CON-TR-4 | Gingras | TR | grey alley | 0.85 | | 10 |
| CON-TR-5 | Cloutier | TR | grey alley | 0.11 | | 16 |
| CON-TR-6 | Jutras | TR | grey alley | 0.08 | | 14 |
| CON-TR-7 | Jutras | TR | grey alley | 0.17 | | 14 |
| CON-TR-8 | Honoré Mercier | TR | grey alley | 0.16 | | 13 |
| CON-TR-9 | Montcalm | TR | grey alley | 0.13 | | 12 |
| CON-VSMPE-10 | Henri-Julien | VSMPE | grey alley | 0.26 | 1 | 25 |
| CON-VSMPE-4 | 51e | VSMPE | grey alley | 0.22 | 0 | 9 |
| CON-VSMPE-8 | Chambord | VSMPE | grey alley | 0.07 | 0 | 15 |
| RV-TR-1 | Avenue 4 | TR | green alley | 0.50 | | 18 |
| RV-TR-11 | Gingras | TR | green alley | 0.04 | | 6 |
| RV-TR-12 | Williams | TR | green alley | 0.13 | | 11 |
| RV-TR-13 | Wolfe | TR | green alley | 0.11 | | 9 |
| RV-TR-2 | Avenue 5 | TR | green alley | 0.29 | | 18 |

| Infrastructure ID | Parallel Street | City | Infrastructure Type | Percent Canopy | Firefly Presence | Number of Points |
|---|---|---|---|---|---|---|
| RV-TR-3 | Amherst | TR | green alley | 0.04 | | 6 |
| RV-TR-4 | Honoré Mercier | TR | green alley | 0.21 | | 10 |
| RV-TR-5 | Brébeuf | TR | green alley | 0.24 | | 17 |
| RV-TR-6 | Godbout | TR | green alley | 0.06 | | 11 |
| RV-TR-7 | Godbout | TR | green alley | 0.15 | | 12 |
| RV-TR-8 | Mgr Cooke | TR | green alley | 0.03 | | 16 |
| RV-TR-9 | Ste Cécile | TR | green alley | 0.05 | | 10 |
| RV-VSMPE-1 | de l'Epée | VSMPE | green alley | 0.21 | | 32 |
| RV-VSMPE-27 | Saint Dominique | VSMPE | green alley | 0.43 | | 15 |
| RV-VSMPE-35 | de Gaspé | VSMPE | green alley | 0.17 | 1 | 7 |
| RV-VSMPE-5 | Hutchison | VSMPE | green alley | 0.22 | | 41 |
| CON-SS-TR-1 | Saint-Louis | TR | street segment | 0.46 | | 7 |
| CON-SS-TR-11 | Ste Angèle | TR | street segment | 0.10 | | 9 |
| CON-SS-TR-12 | Williams | TR | street segment | 0.03 | | 9 |

| Infrastructure ID | Parallel Street | City | Infrastructure Type | Percent Canopy | Firefly Presence | Number of Points |
|---|---|---|---|---|---|---|
| CON-SS-TR-13 | Amherst | TR | street segment | 0.01 | | 8 |
| CON-SS-TR-2 | Saint-Louis | TR | street segment | 0.29 | | 6 |
| CON-SS-TR-3 | Dumoulin | TR | street segment | 0.10 | | 17 |
| CON-SS-TR-4 | Gingras | TR | street segment | 0.00 | | 11 |
| CON-SS-TR-5 | Cloutier | TR | street segment | 0.00 | | 12 |
| CON-SS-TR-6 | Jutras | TR | street segment | 0.03 | | 14 |
| CON-SS-TR-7 | Jutras | TR | street segment | 0.00 | | 18 |
| CON-SS-TR-8 | Honoré Mercier | TR | street segment | 0.08 | | 12 |
| CON-SS-TR-9 | Montcalm | TR | street segment | 0.00 | | 15 |
| CON-SS-VSMPE-10 | Henri-Julien | VSMPE | street segment | 0.50 | 1 | 25 |
| RV-SS-TR-1 | Avenue 4 | TR | green alley | 0.42 | | 14 |
| RV-SS-TR-10 | St Paul | TR | green alley | 0.32 | | 14 |
| RV-SS-TR-12 | Williams | TR | green alley | 0.00 | | 9 |
| RV-SS-TR-13 | Wolfe | TR | green alley | 0.20 | | 9 |
| RV-SS-TR-2 | Avenue 5 | TR | green alley | 0.34 | | 14 |

| Infrastructure ID | Parallel Street | City | Infrastructure Type | Percent Canopy | Firefly Presence | Number of Points |
|---|---|---|---|---|---|---|
| RV-SS-TR-3 | Amherst | TR | green alley | 0.63 | | 6 |
| RV-SS-TR-4 | Honoré Mercier | TR | green alley | 0.00 | | 12 |
| RV-SS-TR-5 | Brébeuf | TR | green alley | 0.33 | | 16 |
| RV-SS-TR-6 | Godbout | TR | green alley | 0.06 | | 12 |
| RV-SS-TR-7 | Godbout | TR | green alley | 0.02 | | 13 |
| RV-SS-TR-8 | Mgr Cooke | TR | green alley | 0.46 | | 12 |
| RV-SS-TR-9 | Ste Cécile | TR | green alley | 0.00 | | 11 |
| SS-VSMPE-17 | 2e | VSMPE | street segment | 0.01 | | 24 |
| SS-VSMPE-21 | 44e | VSMPE | street segment | 0.00 | 0 | 9 |

Table S2. Tree species characteristics for species found in this study. Species represents binomial nomenclature for the species, max height represents the maximum height (in metres) for the species, flowers indicate if the species has showy or non-showy flowers when in bloom, native is a binomial category that indicates if the species is native to the St Lawrence Lowlands ecoregion (1 = yes, 0 = no), invasive is a binomial category that indicates if the species is invasive to the region (1 = yes, 0 = no), functional group represents the functional group assigned by Paquette et al. for that species (Paquette et al., 2021, Praise for diversity: A functional approach to reduce risks in urban forests). Blank cells indicate that we could not find the information for that species.

| Species | Max Height (m) | Flowers | Native_SLL | Invasive | Functional Group |
|---|---|---|---|---|---|
| Abies concolor | 15.2 | not showy | 0 | 0 | 1A |
| Acer freemanii | 15.0 | not showy | 1 | 0 | 2A |
| Acer miyabei | 13.5 | not showy | 0 | 0 | |
| Acer negundo | 12.0 | not showy | 0 | 0 | 2B |
| Acer nigrum | 22.9 | not showy | 0 | 0 | 2A |
| Acer platanoides | 13.5 | not showy | 0 | 1 | 2B |
| Acer platanoides 'Columnare' | 18.2 | not showy | 0 | 1 | 2B |
| Acer rubrum | 15.0 | showy | 1 | 0 | 2A |
| Acer saccharinum | 18.0 | not showy | 1 | 0 | 2A |
| Acer saccharum | 20.5 | not showy | 1 | 0 | 2A |
| Acer tataricum | 5.5 | not showy | 0 | 1 | |
| Aesculus glabra | 9.0 | showy | 0 | 0 | 4B |
| Aesculus hippocastanum | 19.0 | showy | 0 | 0 | 4B |

| Species | Max Height (m) | Flowers | Native_SLL | Invasive | Functional Group |
|---|---|---|---|---|---|
| Amelanchier arborea | 6.5 | showy | 1 | 0 | 2B |
| Catalpa speciosa | 15.0 | showy | 0 | 0 | 2B |
| Celtis occidentalis | 15.0 | not showy | 1 | 0 | 2A |
| Cercidiphyllum japonicum | 15.0 | not showy | 0 | 0 | 5 |
| Dead sp. | | | | | |
| Fagus grandifolia | 18.0 | not showy | 1 | 0 | 2A |
| Fraxinus americana | 19.5 | not showy | 1 | 0 | 2A |
| Fraxinus pennsylvanica | 16.5 | not showy | 1 | 0 | 2A |
| Ginkgo biloba | 19.5 | not showy | 0 | 0 | 1B |
| Gleditsia triacanthos | 15.0 | not showy | 0 | 0 | 2A |
| Gymnocladus dioicus | 20.5 | not showy | 0 | 0 | 4A |
| Juglans cinerea | 15.0 | not showy | 1 | 0 | 4B |
| Larix decidua | 22.0 | not showy | 0 | 0 | 1B |

| Species | Max Height (m) | Flowers | Native_SLL | Invasive | Functional Group |
|---|---|---|---|---|---|
| Larix sp. | 22.0 | not showy | 0 | 0 | 1B |
| Lonicera sp. | 3.4 | showy | 0 | 0 | |
| Maackia amurensis | 7.5 | showy | 0 | 0 | 3B |
| Magnolia sp. | 7.6 | showy | 0 | 0 | 2B |
| Malus baccata | 8.5 | showy | 0 | 0 | 3A |
| Morus alba | 12.0 | not showy | 0 | 0 | 3A |
| Ostrya virginiana | 10.0 | showy | 1 | 0 | 2A |
| Phellodendron amurense | 11.5 | not showy | 0 | 0 | 2B |
| Picea abies | 15.0 | not showy | 0 | 0 | 1A |
| Picea glauca | 15.0 | not showy | 1 | 0 | 1A |
| Picea pungens | 13.5 | not showy | 0 | 0 | 1A |
| Pinus resinosa | 19.5 | not showy | 1 | 0 | 1A |
| Pinus strobus | 19.5 | not showy | 1 | 0 | 1A |

| Species | Max Height (m) | Flowers | Native_SLL | Invasive | Functional Group |
|---|---|---|---|---|---|
| Populus canadensis | 40.0 | not showy | 0 | 0 | 5 |
| Populus deltoides | 26.5 | not showy | 1 | 0 | 5 |
| Populus grandidentata | 21.3 | not showy | 1 | 0 | 5 |
| Prunus cerasus | 9.0 | showy | 0 | 0 | 3 |
| Prunus pensylvanica | 10.7 | showy | 1 | 0 | 2B |
| Prunus virginiana | 8.5 | showy | 1 | 0 | 2B |
| Pyrus domestica | 8.3 | showy | 0 | 0 | 3A |
| Quercus alba | 19.5 | not showy | 1 | 0 | 4A |
| Quercus bicolor | 16.5 | not showy | 1 | 0 | 4A |
| Quercus macrocarpa | 22.5 | not showy | 1 | 0 | 4A |
| Quercus robur ’Fastiagata’ | 16.8 | not showy | 0 | 0 | 4A |
| Quercus rubra | 20.5 | not showy | 1 | 0 | 4A |

| Species | Max Height (m) | Flowers | Native_SLL | Invasive | Functional Group |
|---|---|---|---|---|---|
| Rhamnus catharticus | 6.5 | not showy | 0 | 1 | 2A |
| Rhus typhina | 6.5 | not showy | 1 | 0 | 2B |
| Robinia pseudoacacia | 12.0 | showy | 0 | 0 | 3B |
| Rosa woodsii | 2.0 | showy | 0 | 0 | |
| Salix discolor | 2.9 | showy | 1 | 0 | |
| Saraca indica | 10.0 | showy | 0 | 0 | |
| Sorbus alnifolia | 15.2 | showy | 0 | 0 | 3A |
| Sorbus aucuparia | 9.0 | showy | 0 | 0 | 3A |
| Syringa reticulata | 7.5 | showy | 0 | 0 | 3A |
| Syringa vulgaris | 3.5 | showy | 0 | 0 | 3A |
| Thuja occidentalis | 15.0 | not showy | 1 | 0 | 1A |
| Tilia americana | 21.0 | showy | 1 | 0 | 2B |
| Tilia cordata | 19.5 | showy | 0 | 0 | 2B |

| Species | Max Height (m) | Flowers | Native_SLL | Invasive | Functional Group |
|---|---|---|---|---|---|
| Ulmus | 24.1 | not showy | | | |
| Ulmus 'Homestead' | 18.2 | not showy | 0 | 0 | |
| Ulmus 'Morton Glossy' | 18.0 | not showy | 0 | 0 | 3A |
| Ulmus 'New Horizon' | 18.0 | not showy | 0 | 0 | 3A |
| Ulmus americana | 21.0 | not showy | 1 | 0 | 2B |
| Ulmus crassifolia | 21.3 | not showy | 0 | 0 | |
| Ulmus davidiana 'Discovery' | 24.3 | not showy | 0 | 0 | 3A |
| Ulmus davidiana 'Prospector' | 24.3 | not showy | 0 | 0 | 3A |
| Ulmus frontier | 12.2 | not showy | 0 | 0 | |
| Ulmus hollandica | 18.2 | not showy | 0 | 0 | |
| Ulmus parvifolia | 13.7 | not showy | 0 | 0 | 3A |

| Species | Max Height (m) | Flowers | Native_SLL | Invasive | Functional Group |
|---|---|---|---|---|---|
| Ulmus pumila | 18.0 | not showy | 0 | 1 | 3A |
| Unknown sp. | | | | | |

Table S3. Data displaying the composition of interviewees across different categories. Count indicates the number of interviewees who belonged to that category, and Percent indicates the percentage of interviewees who belonged to that category.

| Variable | Levels | Count | Percent |
|---|---|---|---|
| Age | 20-29 | 7 | 2 |
| Age | 30-39 | 6 | 1 |
| Age | 40-49 | 3 | 1 |
| Age | 50-59 | 7 | 2 |
| Age | 60-69 | 5 | 1 |
| Age | 70-79 | 0 | 0 |
| Age | 80+ | 1 | 0 |
| Gender | Man | 10 | 2 |
| Gender | Woman | 19 | 5 |
| Gender | Other | 1 | 0 |
| Country of Origin | Canada/QC | 16 | 4 |
| Country of Origin | France | 4 | 1 |
| Country of Origin | China | 1 | 0 |

| Variable | Levels | Count | Percent |
|---|---|---|---|
| Country of Origin | Ukraine | 1 | 0 |
| Country of Origin | Ghana | 1 | 0 |
| Country of Origin | Morocco | 1 | 0 |
| Country of Origin | Switzerland | 2 | 0 |
| Country of Origin | Ireland | 1 | 0 |
| Country of Origin | Colombia | 1 | 0 |
| Country of Origin | Iran | 1 | 0 |
| Years in QC | Née au Québec | 16 | 4 |
| Years in QC | Immi 0-4 ans | 4 | 1 |
| Years in QC | Immi 5-9 ans | 3 | 1 |
| Years in QC | Immi 10-14 ans | 1 | 0 |
| Years in QC | Immi 15-19 ans | 0 | 0 |
| Years in QC | Immi 20 ans + | 6 | 1 |
| Household Size | 1 person | 14 | 3 |
| Household Size | 2 people | 10 | 2 |

| Variable | Levels | Count | Percent |
|---|---|---|---|
| Household Size | 3 people | 5 | 1 |
| Household Size | 4 people | 1 | 0 |
| Number of Kids | 0 kids | 18 | 4 |
| Number of Kids | 1 kid | 9 | 2 |
| Number of Kids | 2 kids | 2 | 0 |
| Number of Kids | 3 kids | 1 | 0 |
| Education Level | Universitaire | 23 | 6 |
| Education Level | Collégial | 5 | 1 |
| Education Level | Secondaire | 2 | 0 |
| Employment Status | Student | 7 | 2 |
| Employment Status | Retired | 4 | 1 |
| Employment Status | Full-time | 16 | 4 |
| Employment Status | Unemployed | 1 | 0 |
| Employment Status | Self-employed | 2 | 0 |
| Annual Salary | -19 999$ | 4 | 1 |

| Variable | Levels | Count | Percent |
|---|---|---|---|
| Annual Salary | 20 000$-29 999$ | 4 | 1 |
| Annual Salary | 30 000$-39 999$ | 2 | 0 |
| Annual Salary | 40 000$-49 999$ | 0 | 0 |
| Annual Salary | 50 000$-59 999$ | 5 | 1 |
| Annual Salary | 60 000$-69 999$ | 1 | 0 |
| Annual Salary | 70 000$-79 999$ | 2 | 0 |
| Annual Salary | 80 000$-89 999$ | 0 | 0 |
| Annual Salary | 90 000$-99 999$ | 0 | 0 |
| Annual Salary | 100 000$ + | 9 | 2 |
| Car | Yes | 14 | 3 |
| Car | No | 16 | 4 |
| Housing Type | Apartment | 18 | 4 |
| Housing Type | Single family home | 8 | 2 |

| Variable | Levels | Count | Percent |
|---|---|---|---|
| Housing Type | Condo | 3 | 1 |
| Housing Type | Coop | 1 | 0 |
| Alley Knowledge | Know the alleys | 9 | 2 |
| Alley Knowledge | Don’t know the alleys | 20 | 5 |
| Alley Knowledge | Know one alley | 1 | 0 |
| City | Montréal | 16 | 4 |
| City | Trois-Rivières | 14 | 3 |
| Montreal Neighbourhood | Villeray | 12 | 3 |
| Montreal Neighbourhood | Parc-Extension | 3 | 1 |
| Montreal Neighbourhood | Saint-Michel | 1 | 0 |
| TR Neighbourhood | Sainte-Cécile | 4 | 1 |
| TR Neighbourhood | Saint-Patrick | 5 | 1 |
| TR Neighbourhood | Vieux Trois-Rivières | 2 | 0 |

| Variable | Levels | Count | Percent |
|---|---|---|---|
| TR Neighbourhood | Saint-Sacrement | 3 | 1 |

Table S4. Data showing the number of mentions that each ecosystem service received in the interviews. Ecosystem service category categorizes each ecosystem service using the four categories proposed in the Millenium Ecosystem Assessment (2005). Total mentions refers to the total number of times an ecosystem service was mentioned, regardless of if the same person brought it up multiple times. Number of people mentioned refers to the total number of people who mentioned that ecosystem service, without incorporating how many times they mentioned it. A darker colour depicts a higher number of mentions.

| Ecosystem Service | Ecosystem Service Category | Total Mentions | Number of People Mentioned |
|---|---|---|---|
| Cultural and aesthetic values of greening | Cultural | 66 | 26 |
| Temperature regulation | Regulating | 59 | 25 |
| Cultural value of trees | Cultural | 58 | 20 |
| Aesthetic value of flowers | Cultural | 52 | 17 |
| Supporting wildlife | Supporting | 48 | 18 |

| Ecosystem Service | Ecosystem Service Category | Total Mentions | Number of People Mentioned |
|---|---|---|---|
| Provisioning of food from plants | Provisionir | 40 | 17 |
| Cultural value of vegetation management | Cultural | 38 | 15 |
| Presence of biodiversity | Supportinç | 33 | 14 |
| Cultural value of large trees | Cultural | 31 | 20 |
| Cultural value of wild plants | Cultural | 27 | 13 |
| Aesthetic value of vines | Cultural | 23 | 12 |
| Cultural experience of odours | Cultural | 19 | 12 |

| Ecosystem Service | Ecosystem Service Category | Total Mentions | Number of People Mentioned |
|---|---|---:|---:|
| Cultural value of vegetation maintenance | Cultural | 18 | 14 |
| Aesthetic value of colours | Cultural | 13 | 8 |
| Stormwater management | Regulating | 10 | 7 |
| Presence of native species | Supporting | 8 | 4 |
| Permeable surface management | Regulating | 3 | 2 |
| Air quality regulation | Regulating | 2 | 2 |
| Production of allergens | Provisionin | 1 | 1 |

### B) Methodological Details

#### Canopy Cover

We calculated percent canopy cover of all green alleys, grey alleys, and street segments using publicly available LiDAR data collected in 2020 at a spatial resolution of 1 m (Institut National de Santé Publique du Québec and Gouvernement du Québec, 2022). For street segments, we calculated the canopy cover of sidewalks using the same dataset. Sidewalk boundaries were extracted from an open dataset for VSMPE (Ville de Montréal, 2024) and hand-drawn for TR using QGIS 3.16.8 (QGIS Development Team, 2020).

#### Tree Diversity

Tree diversity was assessed in two ways. First, we calculated the species richness of each alley and street segment by counting the number of unique tree species. We identified the species of each individual in the field (Farrar, 1995; Little, 1980). If a tree species could not be identified in the field, it was marked as unknown (n = 1/1,469). If a tree was dead, it was still recorded but was removed from the diversity calculations (n = 7/1,469). We recorded a total of 75 species (Table S2).

The second diversity metric was functional diversity. Using the original dataset and methodology from Paquette et al. (2021), we assigned a functional group to each of the tree species present in our dataset (Belluau et al., 2021, Table S2). The database published online did not have a functional group assignment for 15 of our species (20%, predominantly cultivars), thus those species were removed from the functional diversity calculations. If a cultivar or hybrid was not present in the database, but both parent species were and they shared the same functional group, we assigned the functional group of the parent species to the cultivar. We then summed how many functional groups each alley or street segment contained.

#### Proportion of Invasives/Natives

To assess the proportion of invasive and native species at each study site, we assigned a status of native or invasive to each of our study species (Padvaiskas et al., subm, Philp, 2024, Table S2). A species' native range was evaluated using the St. Lawrence Lowlands ecoregion. The St. Lawrence Lowlands ecoregion surrounds the St. Lawrence River ranging from Quebec City south through Montreal and just beyond Ottawa (Minister of Industry, 2010). If the native range of a species fell within a region we deemed it native to that region (Canadian Wildlife Federation, 2024; Dirr and Warren, 2019; Fryer, 2018; Government of Canada, 2013; The Morton Arboretum, 2022; Tree Canada, 2024; United States Department of Agriculture and Natural Resources Conservation Service, 2024; Woody Invasives of the Great Lakes Collaborative, 2019).

We categorized invasive species based on sources within the Eastern Temperate Forest ecoregion (The Morton Arboretum, 2022; Woody Invasives of the Great Lakes Collaborative, 2019). This is a larger ecoregion which runs west to east from Ontario to Nova Scotia and north to south from Southern Canada to Florida, USA (bplant.org, 2024). Any cultivar of a parent species that was considered invasive was also recorded as invasive. Hybrids with an invasive parent were noted as invasive only if the invasive traits persisted in the hybrid (Dirr and Warren, 2019). Once each species was assigned a status, we calculated the proportion of stems at each site that were native and invasive.

#### Vegetative Complexity

We measured vegetative complexity by walking through each study site and stopping every 10 m. At each stop, we assessed which of the 5 following vegetative layers were present: ground cover, herbaceous vegetation, wall vegetation, shrubs, or canopy. If the presence of one of the layers was less than 1% of the area, it was ignored (e.g., one dandelion does not indicate the presence of an herbaceous vegetation layer). We define wall vegetation as vines and crawling vegetation that live on fences or walls. Average vegetative complexity of alleys and street segments was calculated by summing the total number of layers counted and then dividing by the number of points measured.

##### Firefly Presence

To assess the habitat quality of our study alleys, we selected a subset of alleys in VSMPE and performed a presence/absence survey of fireflies (Family: Lampyridae). Fireflies are known bioindicators in cities, as they require leaf litter, access to water, and low levels of light pollution (Firefly Atlas et al., 2023; Picchi et al., 2013). We selected the 28 green alleys with the highest canopy cover and most vegetative complexity for our subset, the five grey alleys with the least canopy cover and vegetative complexity as our controls, and the associated street segment for all 34 alleys surveyed. We sampled in groups of three between dusk and early nightfall, between 9:00 and 11:30 pm. Each person surveyed a section of the alley for five minutes and watched for the presence of fireflies. When a firefly was seen, the sampler recorded its presence. After 5 minutes, each person rotated spots and repeated the survey in the next segment. We did this three times for each alley. We did not sample in the rain, when firefly flight does not occur (Firefly Atlas et al., 2023; Picchi et al., 2013). If the presence of a firefly was noted in any of the surveys, the alley was assigned as having fireflies present.

##### DBH

We assessed the presence of large trees in two ways, we calculated the mean observed tree DBH and the mean potential height for each study site. To calculate mean observed DBH, we evaluated tree size by identifying and measuring the diameter at breast height (DBH) of every tree (defined as a woody plant $> 2$ m tall) found in the alley. If a tree had a split stem with the split above or near DBH, we measured the DBH at the narrowest part of the stem below the fork. If the split occurred below DBH but above 6" from the ground, we measured each stem individually and calculated DBH by taking the square root of the sum of all squared stem DBHs. If the split occurred below 6” from the ground, we counted the stems as separate individuals (Magarik et al., 2020). We then took the average DBH of all the trees in an alley or street segment.

#### Interview Questions

**Sociodemographic data:**

- Age
- Gender
- Ethnocultural group
- Time since arriving in Quebec
- Household composition
- Number of kids / their ages
- Level of education
- Employment status
- Annual revenue bracket
- Housing type
- Car ownership

**The neighbourhood**

1. What represents the neighbourhood for you? What characterizes it? What do you value in the neighbourhood? (or what would you be ready to invest in?)
2. What are the boundaries of your neighbourhood?
3. What is here in terms of public green space management?
4. According to you, what are the objectives and intended benefits of these developments for the riverside community? Are they being achieved?
5. What are the downsides of the developments for the riverside community?
6. What do you like the most or the least about these green spaces?

7. How do these spaces improve the neighbourhood (social, road and urban safety, mobility, health)
8. According to you, what does the city do to ensure the management, quality of development, and the management of these spaces? Are they doing enough?

**The neighbourhood, green spaces, and changes**

1. What has changed in your neighbourhood in the last years (green infrastructure, management of the neighbourhood, demographics, social composition, etc.)?
2. When do you remember the changes starting?
3. Who makes the changes and for what reasons?
4. Has there been citizen mobilization?
5. Who has changed their activities since the changes?
6. How have the neighbourhood relationships changed (residential, institutional)?
7. Do you make any links between the management of green spaces and the changes in the neighbourhood?

**The walk starts**

1. Discuss what attracts your attention here?
2. Discuss how you feel here?
3. Discuss what activities you do here?
4. Discuss the interventions/management/equipment (anthropogenic elements)
5. Discuss the richness, diversity, cohabitation, and conditions of the space?

**Before concluding**

1. Which green spaces do you like the most and the least? Why?
2. Which green spaces should be improved? Why?

3. When comparing the spaces we visited, does your perception of the spaces change?
4. After the visit, what do you think is different (or similar) between green spaces and non-green spaces? What do the green spaces you visit have in common?
5. How do these spaces relate to other green spaces in the city/nature in the city? What do these green spaces have in common with other green spaces in the city?
6. Do you have other comments to share with us?

#### C) Figures

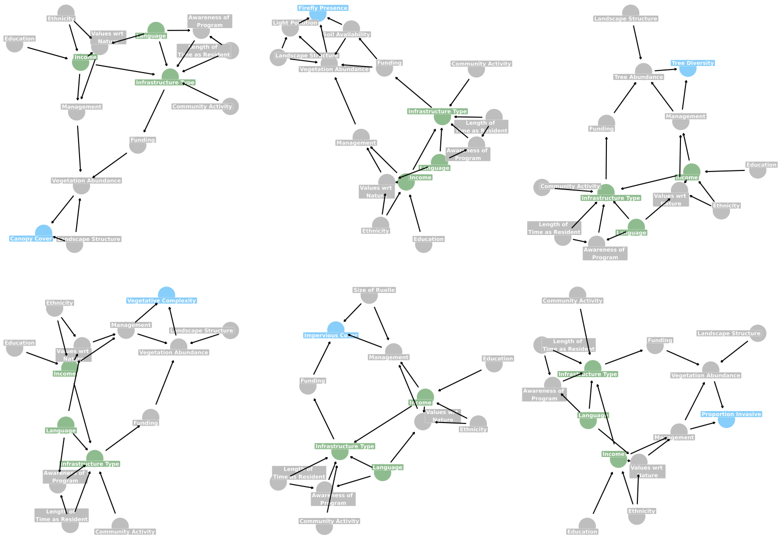

Figure S1a. DAGs that indicate the system assumptions for biodiversity support indices measured in this study. Blue indicates the response variable of interest, green indicates the variable of interest and all variables adjusted for.

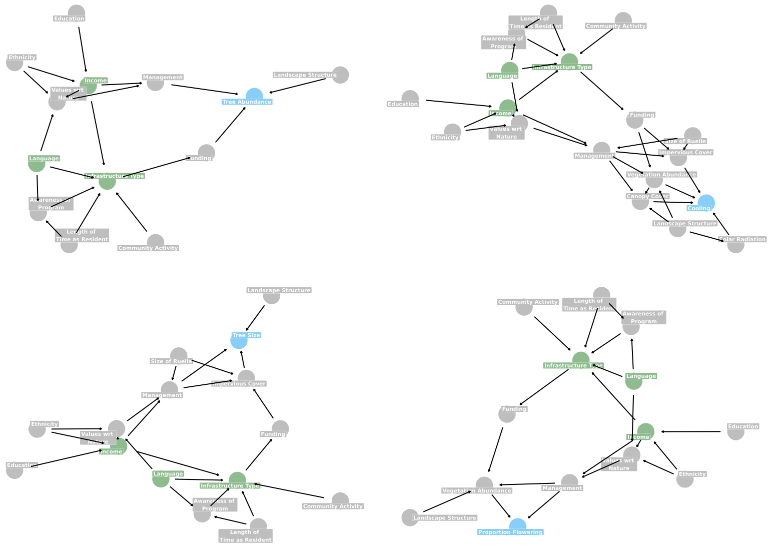

Figure S1b. DAGs that indicate the system assumptions for ecosystem service indices measured in this study. Blue indicates the response variable of interest, green indicates the variable of interest and all variables adjusted for.

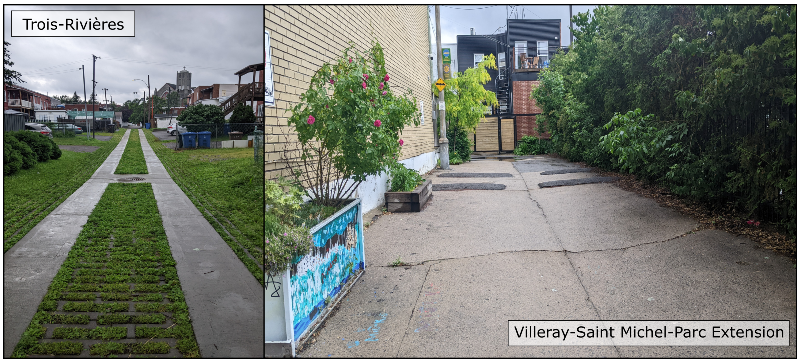

Figure S2. Examples of typical green alleys in Trois-Rivières and Villeray-Saint Michel-Parc Extension. Photos were taken by Isabella C Richmond in the summer of 2023.

### D) Equations

Below are the mathstats formulas for each of the models included in the paper.

**Canopy Cover**

1. VSMPE

   $Canopy_i \sim Normal(\mu_i, \sigma)\mu_i = \alpha_{Qsocio[i]} + \beta_{type[i]} + \beta_{medinc} medinc_i + \beta_{french} french_i + \beta_{english} english_i + \beta_{none}$[illegible]

   Terms:

- `Canopy` is the centered and scaled percent canopy cover of each site
- `type` is the type of infrastructure (i.e., green alley, grey alley, street segment)
- `medinc` is the centered and scaled median income of the dissemination area surrounding the site
- `french` is the centered and scaled percent of residents in the dissemination area surrounding the site that speak french
- `english` is the centered and scaled percent of residents in the dissemination area surrounding the site that speak english
- `none` is the centered and scaled percent of residents in the dissemination area surrounding the site that do not speak french or english

2. TR

$Canopy \sim Normal(\mu_i, \sigma)\mu_i = \alpha + \beta_1 green_{[i]} + \beta_2 street_{[i]} \alpha \sim Normal(0, 0.7) \beta_j \sim Normal(0, 0.7) \sigma \sim Exponential($[illegible]

Terms:

- `Canopy` is the centered and scaled percent canopy cover of each site

- `green` is the factor level from `type` that indicates green alleys. Grey alleys are the default level, absorbed into the intercept.

- `street` is the factor level from `type` that indicates street segments. Grey alleys are the default level, absorbed into the intercept.

##### Species Richness

3. VSMPE

$$SpeciesRichness_i \sim Gamma{-}Poisson(\lambda_i, \phi) log(\lambda_i) = \alpha_{Qsocio[i]} + \beta_{type[i]} + \beta_{medinc} medinc_i + \beta_{french} french_i + \beta_{engl}$$

Terms:

- `Species Richness` is number of unique tree species found at each site
- `Q socio` is the neighbourhood of the site (i.e., Villeray, Saint-Michel, or Parc Extension)
- `type` is the type of infrastructure (i.e., green alley, grey alley, or street segment)
- `medinc` is the centered and scaled median income of the dissemination area surrounding the site
- `french` is the centered and scaled percent of residents in the dissemination area surrounding the site that speak french
- `english` is the centered and scaled percent of residents in the dissemination area surrounding the site that speak english
- `none` is the centered and scaled percent of residents in the dissemination area surrounding the site that do not speak french or english

4. TR

$$SpeciesRichness_i \sim Poisson(\lambda\_i) log(\lambda\_i) = \alpha + \beta_1 green{*}[i] + \beta_2 street_{[i]} \alpha \sim Normal(0, 0.5) \beta_j \sim Normal(0, 0.2)$$

Terms:

- `Canopy` is the centered and scaled percent canopy cover of each site

- `green` is the factor level from `type` that indicates green alleys. Grey alleys are the default level, absorbed into the intercept.
- `street` is the factor level from `type` that indicates street segments. Grey alleys are the default level, absorbed into the intercept.

###### Functional Diversity

5. VSMPE

$$FunctionalGroups_i \sim Binomial(9, p_i) logit(p_i) = \alpha_{Qsocio[i]} + \beta_{type[i]} + \beta_{medinc} medinc_i + \beta_{french} french_i + \beta_{english} eng$$ [illegible]

Terms:

- `Functional Groups` is number of unique functional groups found at each site
- `Q socio` is the neighbourhood of the site (i.e., Villeray, Saint-Michel, or Parc Extension)
- `type` is the type of infrastructure (i.e., green alley, grey alley, or street segment)
- `medinc` is the centered and scaled median income of the dissemination area surrounding the site
- `french` is the centered and scaled percent of residents in the dissemination area surrounding the site that speak french
- `english` is the centered and scaled percent of residents in the dissemination area surrounding the site that speak english
- `none` is the centered and scaled percent of residents in the dissemination area surrounding the site that do not speak french or english

6. TR

$$FunctionalGroups_i \sim Binomial(9, p_i) logit(p_i) = \alpha + \beta_1 green_{[i]} + *\beta_2 street_{[i]} \alpha \sim Normal(0, 0.5) \beta_j \sim Normal(0, 0.$$

Terms:

- `Functional Groups` is number of unique functional groups found at each site
- `green` is the factor level from `type` that indicates green alleys. Grey alleys are the default level, absorbed into the intercept.
- `street` is the factor level from `type` that indicates street segments. Grey alleys are the default level, absorbed into the intercept.

##### Proportion of Native Species

7. VSMPE

$$ProportionNative_i \sim Binomial(N_i, p_i) logit(p_i) = \alpha_{Qsocio[i]} + \beta_{type[i]} + \beta_{medinc} medinc_i + \beta_{french} french_i + \beta_{english} en$$

Terms:

- `N` is the total number of trees found at each site
- `Proportion Native` is number of native trees found at each site
- `Q socio` is the neighbourhood of the site (i.e., Villeray, Saint-Michel, or Parc Extension)
- `type` is the type of infrastructure (i.e., green alley, grey alley, or street segment)
- `medinc` is the centered and scaled median income of the dissemination area surrounding the site
- `french` is the centered and scaled percent of residents in the dissemination area surrounding the site that speak french

- `english` is the centered and scaled percent of residents in the dissemination area surrounding the site that speak english

- `none` is the centered and scaled percent of residents in the dissemination area surrounding the site that do not speak french or english

8. TR

   $ProportionNative_i \sim Binomial(N_i, p_i) logit(p_i) = \alpha + \beta_1 green_{[i]} + \beta_2 street_{[i]} \alpha \sim Normal(0, 1.5) \beta_j \sim Norma$

- `N` is the total number of trees found at each site
- `Proportion Native` is number of native trees found at each site
- `green` is the factor level from `type` that indicates green alleys. Grey alleys are the default level, absorbed into the intercept.
- `street` is the factor level from `type` that indicates street segments. Grey alleys are the default level, absorbed into the intercept.

##### Proportion of Invasive Species

9. VSMPE

$$ProportionInvasive_i \sim Binomial(N_i, p_i) logit(p_i) = \alpha_{Qsocio[i]} + \beta_{type[i]} + \beta_{medinc} medinc_i + \beta french french_i + \beta_{englis}$$

Terms:

- `N` is the total number of trees found at each site

- `Proportion Invasive` is number of invasive trees found at each site

- `Q socio` is the neighbourhood of the site (i.e., Villeray, Saint-Michel, or Parc Extension)

- `type` is the type of infrastructure (i.e., green alley, grey alley, or street segment)

- `medinc` is the centered and scaled median income of the dissemination area surrounding the site - `french` is the centered and scaled percent of residents in the dissemination area surrounding the site that speak french
- `english` is the centered and scaled percent of residents in the dissemination area surrounding the site that speak english
- `none` is the centered and scaled percent of residents in the dissemination area surrounding the site that do not speak french or english

10. TR

$$ProportionInvasive_i \sim Binomial(N_i, p_i) logit(p_i) = \alpha + \beta_1 green_{[i]} + \beta_2 street_{[i]} \alpha \sim Normal(0, 1.5) \beta_j \sim Normal(0,$$

Terms:

- `N` is the total number of trees found at each site
- `Proportion Invasive` is number of invasive trees found at each site
- `green` is the factor level from `type` that indicates green alleys. Grey alleys are the default level, absorbed into the intercept.
- `street` is the factor level from `type` that indicates street segments. Grey alleys are the default level, absorbed into the intercept.

###### Vegetative Complexity

11. VSMPE

$$Complexity_i \sim Normal(\mu_i, \sigma) \mu_i = \alpha_{Qsocio[i]} + \beta_{type[i]} + \beta_{medinc} medinc_i + \beta_{french} french_i + \beta_{english} english_i + \beta_{none} n$$

Terms:

- `Complexity` is the centered and scaled average vegetative complexity of each site
- `type` is the type of infrastructure (i.e., green alley, grey alley, street segment)
- `medinc` is the centered and scaled median income of the dissemination area surrounding the site
- `french` is the centered and scaled percent of residents in the dissemination area surrounding the site that speak french
- `english` is the centered and scaled percent of residents in the dissemination area surrounding the site that speak english
- `none` is the centered and scaled percent of residents in the dissemination area surrounding the site that do not speak french or english

12. TR

$$Complexity_i \sim Normal(\mu_i, \sigma) \mu_i = \alpha + \beta_1 green_{[i]} + \beta_2 street_{[i]} \alpha \sim Normal(0, 0.5) \beta_j \sim Normal(0, 0.5)$$

Terms:

- `Complexity` is the centered and scaled average vegetative complexity of each site
- `green` is the factor level from `type` that indicates green alleys. Grey alleys are the default level, absorbed into the intercept.
- `street` is the factor level from `type` that indicates street segments. Grey alleys are the default level, absorbed into the intercept.

##### Firefly Presence

13. VSMPE

$$FireflyPresence_i \sim Bernoulli(p_i)logit(p_i) = \alpha_{Qsocio[i]} + \beta_{type[i]} + \beta_{medinc}medinc_i + \beta_{french}french_i + \beta_{english}englis$$ [illegible]

Terms:

- `Firefly Presence` is binary success of observing a firefly at a site (1 = presence, 0 = absence)
- `Q socio` is the neighbourhood of the site (i.e., Villeray, Saint-Michel, or Parc Extension)
- `type` is the type of infrastructure (i.e., green alley, grey alley, or street segment)
- `medinc` is the centered and scaled median income of the dissemination area surrounding the site
- `french` is the centered and scaled percent of residents in the dissemination area surrounding the site that speak french
- `english` is the centered and scaled percent of residents in the dissemination area surrounding the site that speak english
- `none` is the centered and scaled percent of residents in the dissemination area surrounding the site that do not speak french or english

##### Temperature

14. VSMPE

$$Temperature_i \sim Normal(\mu_i, \sigma)\mu_i = \alpha_{date[i]} + \alpha_{Qsocio[i]} + \alpha_{ID[i]} + \gamma type[i] + \gamma_{tod[i]} + \gamma_{doy[i]} + \beta_{type}tod + \beta_{tod}doy + \beta_{doy}ty$$ [illegible]

Terms:

- $\alpha$ and $\gamma$ both represent intercepts, $\alpha$ is used for random effects and $\gamma$ for fixed interaction effecst
- `Temperature` is the centered and scaled temperature measurement in degrees Celsius
- `date` is the calendar date represented as a character with a different level for each day
- `Q socio` is the neighbourhood of the site (i.e., Villeray, Saint-Michel, or Parc Extension)
- `ID` is the individual identifier for each infrastructure (e.g., VSMPE-RV-1)
- `type` is the type of infrastructure (i.e., green alley, grey alley, or street segment)
- `tod` is the time of day of each measurement, split into two categories; day (after sunrise/before sunset) and night (after sunset/before sunrise)
- `doy` is the numeric and continuous value representing day of year, which has been subtracted by the first day of sampling so that it starts at 0
- `medinc` is the centered and scaled median income of the dissemination area surrounding the site
- `french` is the centered and scaled percent of residents in the dissemination area surrounding the site that speak french
- `english` is the centered and scaled percent of residents in the dissemination area surrounding the site that speak english
- `none` is the centered and scaled percent of residents in the dissemination area surrounding the site that do not speak french or english

15. TR

$$Temperature_i \sim Normal(\mu_i, \sigma)\mu_i = \alpha_{date[i]}+\alpha_{ID[i]}+\gamma_{type[i]}+\gamma_{tod[i]}+\gamma_{doy[i]}+\beta_{type}tod+\beta_{tod}doy+\beta_{doy}type+\beta_{type[i]}+ [illegible]$$

Terms:

- $\alpha$ and $\gamma$ both represent intercepts, $\alpha$ is used for random effects and $\gamma$ for fixed interaction effects
- `Temperature` is the centered and scaled temperature measurement in degrees Celsius
- `date` is the calendar date represented as a character with a different level for each day
- `ID` is the individual identifier for each infrastructure (e.g., VSMPE-RV-1 )
- `type` is the type of infrastructure (i.e., green alley, grey alley, or street segment)
- `tod` is the time of day of each measurement, split into two categories; day (after sunrise/before sunset) and night (after sunset/before sunrise)
- `doy` is the numeric and continuous value representing day of year, which has been subtracted by the first day of sampling so that it starts at 0

##### Tree Abundance

16. VSMPE

$$TreeAbundance_i \sim Gamma{-}Poisson(\lambda_i, \phi)log(\lambda_i) = \alpha_{Qsocio[i]} + \beta_{type[i]} + \beta_{medinc}medinc_i + \beta_{french}french_i + \beta_{engli}$$

Terms:

- `Tree Abundance` is number of individual trees found at a site
- `Q socio` is the neighbourhood of the site (i.e., Villeray, Saint-Michel, or Parc Extension)
- `type` is the type of infrastructure (i.e., green alley, grey alley, or street segment)
- `medinc` is the centered and scaled median income of the dissemination area surrounding the site

- `french` is the centered and scaled percent of residents in the dissemination area surrounding the site that speak french
- `english` is the centered and scaled percent of residents in the dissemination area surrounding the site that speak english
- `none` is the centered and scaled percent of residents in the dissemination area surrounding the site that do not speak french or english

17. TR

$$TreeAbundance_i \sim Poisson(\lambda\_i) log(\lambda\_i) = \alpha + \beta_1 green_{[i]} + \beta_2 street_{[i]} \alpha \sim Normal(0, 0.5) \beta_j \sim Normal(0, 0.2)$$

Terms:

- `Tree Abundance` is number of individual trees found at a site
- `green` is the factor level from `type` that indicates green alleys. Grey alleys are the default level, absorbed into the intercept.
- `street` is the factor level from `type` that indicates street segments. Grey alleys are the default level, absorbed into the intercept.

##### Tree Size (Diameter at Breast Height)

18. VSMPE

$$DBH_i \sim Normal(\mu_i, \sigma) \mu_i = \alpha_{Qsocio[i]} + \beta_{type[i]} + \beta_{medinc} medinc_i + \beta_{french} french_i + \beta_{english} english_i + \beta_{none} none_i \beta_j$$

Terms:

- `DBH` is the centered and scaled mean diameter at breast height (DBH) of trees at each site

- `type` is the type of infrastructure (i.e., green alley, grey alley, street segment)
- `medinc` is the centered and scaled median income of the dissemination area surrounding the site
- `french` is the centered and scaled percent of residents in the dissemination area surrounding the site that speak french
- `english` is the centered and scaled percent of residents in the dissemination area surrounding the site that speak english
- `none` is the centered and scaled percent of residents in the dissemination area surrounding the site that do not speak french or english

19. TR

$$DBH_i \sim Normal(\mu_i, \sigma)\mu_i = \alpha + \beta_1 green_{[i]} + \beta_2 street_{[i]} \alpha \sim Normal(0, 0.7) \beta_j \sim Normal(0, 0.7) \sigma \sim Exponent$$ [illegible]

Terms:

- `DBH` is the centered and scaled mean diameter at breast height (DBH) of trees at each site
- `green` is the factor level from `type` that indicates green alleys. Grey alleys are the default level, absorbed into the intercept.
- `street` is the factor level from `type` that indicates street segments. Grey alleys are the default level, absorbed into the intercept.

##### Tree Size (Potential Maximum Height)

20. VSMPE

$$MaxHeight_i \sim Normal(\mu_i, \sigma)\mu_i = \alpha_{Qsocio[i]} + \beta_{type[i]} + \beta_{medinc} medinc_i + \beta_{french} french_i + \beta_{english} english_i + \beta_{none} n$$ [illegible]

Terms:

- `Max Height` is the centered and scaled mean maximum (potential) height of trees at each site

- `type` is the type of infrastructure (i.e., green alley, grey alley, street segment)
- `medinc` is the centered and scaled median income of the dissemination area surrounding the site
- `french` is the centered and scaled percent of residents in the dissemination area surrounding the site that speak french
- `english` is the centered and scaled percent of residents in the dissemination area surrounding the site that speak english
- `none` is the centered and scaled percent of residents in the dissemination area surrounding the site that do not speak french or english

21. TR

$$MaxHeight_i \sim Normal(\mu_i, \sigma) \mu_i = \alpha + \beta_1 green_{[i]} + \beta_2 street_{[i]} \alpha \sim Normal(0, 0.7) \beta_j \sim Normal(0, 0.7) \sigma \sim Exponen$$ [illegible]

Terms:

- `Max Height` is the centered and scaled mean maximum (potential) height of trees at each site
- `green` is the factor level from `type` that indicates green alleys. Grey alleys are the default level, absorbed into the intercept.
- `street` is the factor level from `type` that indicates street segments. Grey alleys are the default level, absorbed into the intercept.

###### Proportion of Trees with Showy Flowers

22. VSMPE

$$ProportionShowy_i \sim Binomial(N_i, p_i) logit(p_i) = \alpha_{Qsocio[i]} + \beta_{type[i]} + \beta_{medinc} medinc_i + \beta_{french} french_i + \beta_{english} en$$

Terms:

- `N` is the total number of trees found at each site
- `Proportion Showy` is number of trees with showy flowers found at each site
- `Q socio` is the neighbourhood of the site (i.e., Villeray, Saint-Michel, or Parc Extension)
- `type` is the type of infrastructure (i.e., green alley, grey alley, or street segment)
- `medinc` is the centered and scaled median income of the dissemination area surrounding the site
- `french` is the centered and scaled percent of residents in the dissemination area surrounding the site that speak french
- `english` is the centered and scaled percent of residents in the dissemination area surrounding the site that speak english
- `none` is the centered and scaled percent of residents in the dissemination area surrounding the site that do not speak french or english

23. TR

$$ProportionShowy_i \sim Binomial(N_i, p_i) logit(p_i) = \alpha + \beta_1 green_{[i]} + \beta_2 street_{[i]} \alpha \sim Normal(0, 0.5) \beta_j \sim Norma$$

Terms:

- `N` is the total number of trees found at each site
- `Proportion Showy` is number of trees with showy flowers found at each site
- `green` is the factor level from `type` that indicates green alleys. Grey alleys are the default level, absorbed into the intercept.
- `street` is the factor level from `type` that indicates street segments. Grey alleys are the default level, absorbed into the intercept.

#### E) Prior Predictive Checks

Prior predictive checks are used to ensure that the values selected for priors for our models allow a biologically reasonable range of values. For numeric predictor variables, we simulate predictive draws for prior only models and visualize the slope/intercept of the values. We then do a "posterior predictive check" but with the prior only model, to see if the data is captured in the priors. Note that all data is scaled and centered in these data.

For Trois-Rivieres, there is only a categorical predictor variable. Therefore, only the posterior predictive check is presented.

##### Villeray-Saint Michel-Parc Extension

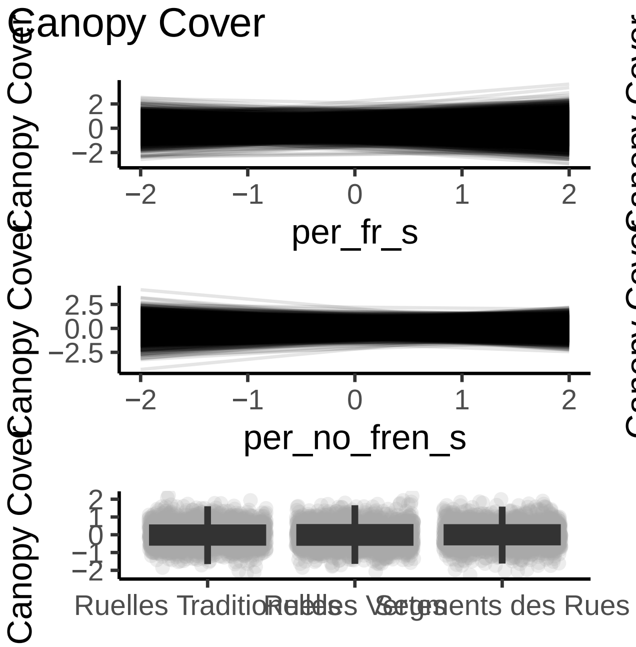

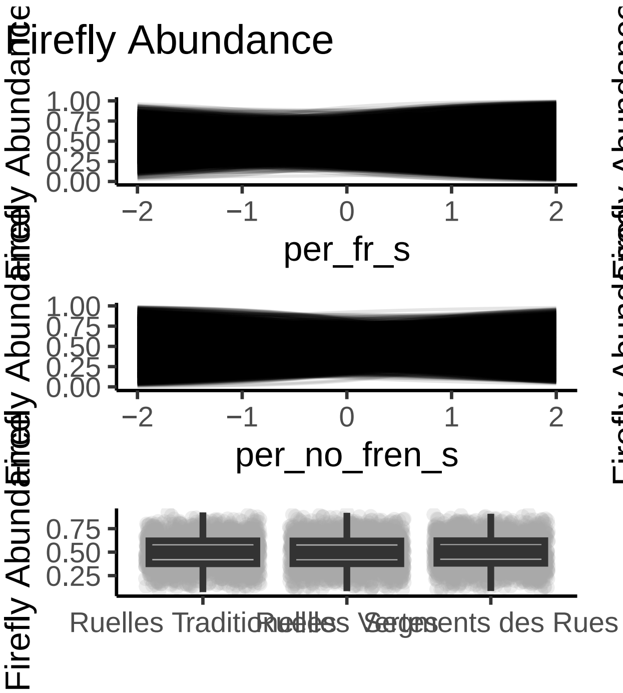
Firefly Abundance
Firefly Abundance
1.00
0.75
0.50
0.25
0.00
−2
−1
0
1
2
per_fr_s
per_en_s
per_no_fren_s
medinc_s
Ruelles Traditionnelles
Ruelles Vertes
Segments des Rues

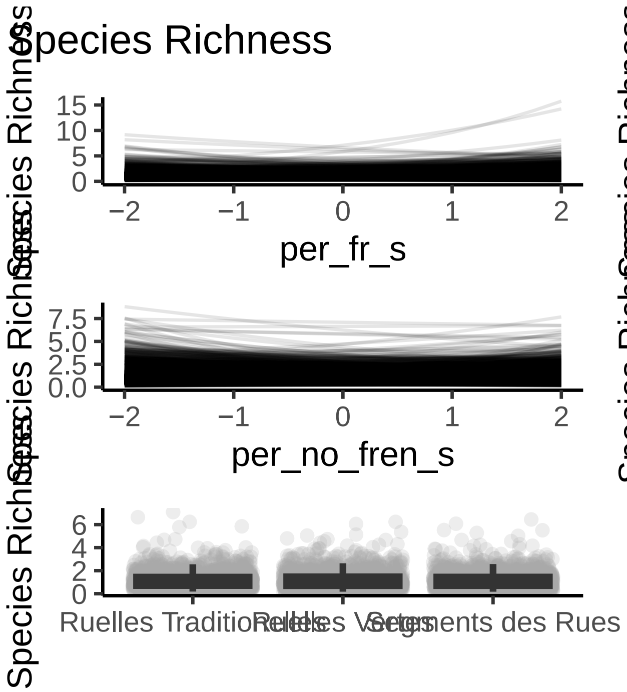
Species Richness
Species Richness
per_fr_s
per_en_s
per_no_fren_s
medinc_s
Ruelles Traditionnelles
Ruelles Vertes
Segments des Rues

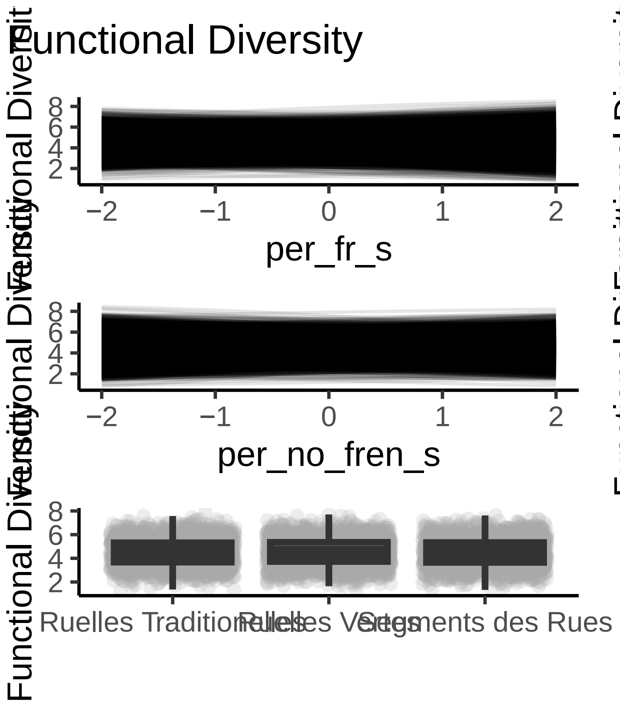
Functional Diversity
Functional Diversity
per_fr_s
per_en_s
per_no_fren_s
medinc_s
Ruelles Traditionnelles
Ruelles Vertes
Segments des Rues

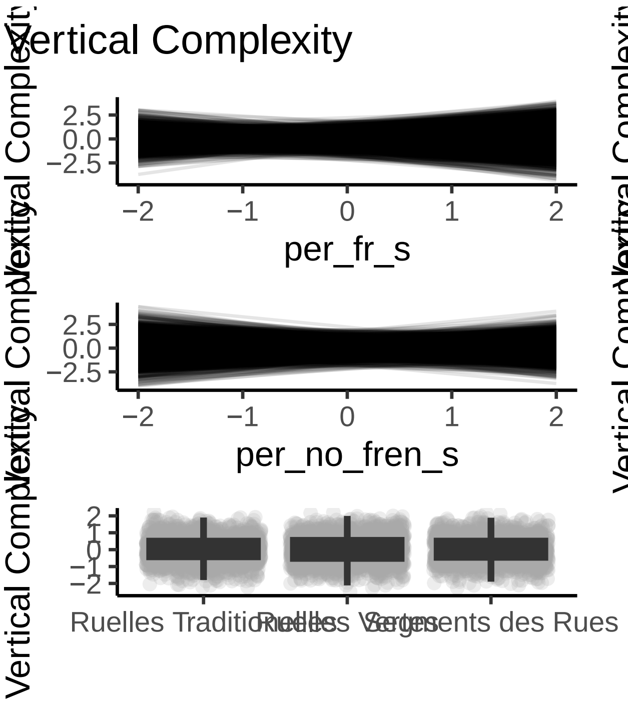
Vertical Complexity
Vertical Complexity
2.5
0.0
−2.5
−2
−1
0
1
2
per_fr_s
per_en_s
per_no_fren_s
medinc_s
Ruelles Traditionnelles
Ruelles Vertes
Segments des Rues

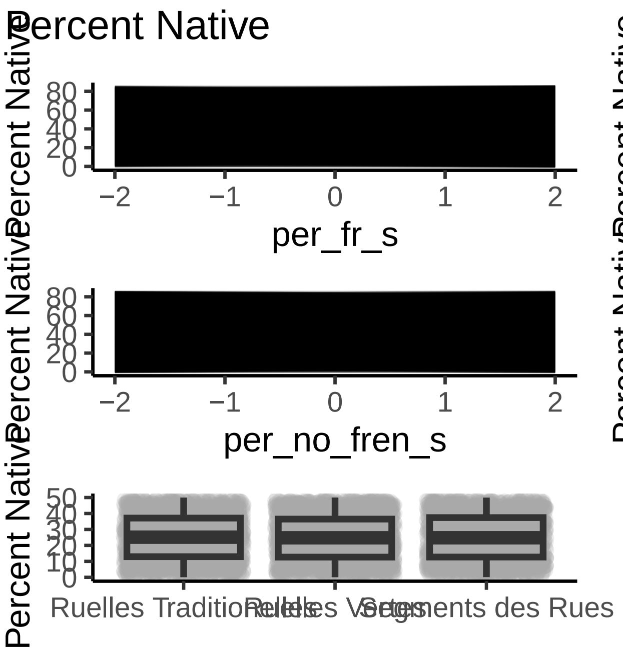
Percent Native
Percent Native
per_fr_s
Percent Native
per_en_s
Percent Native
per_no_fren_s
Percent Native
medinc_s
Percent Native
Ruelles Traditionnelles
Ruelles Vertes
Segments des Rues

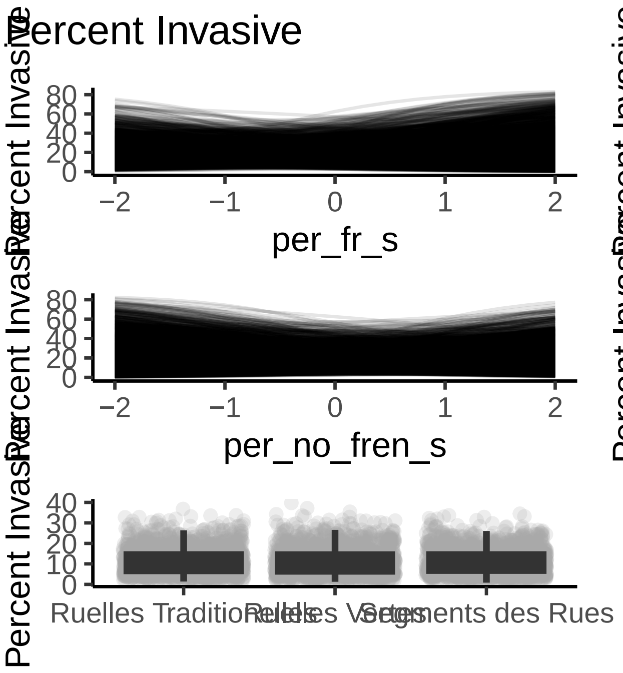

Percent Invasive
Percent Invasive
0
20
40
60
80
−2
−1
0
1
2
per_fr_s
Percent Invasive
per_en_s
Percent Invasive
per_no_fren_s
Percent Invasive
medinc_s
Percent Invasive
0
10
20
30
40
Ruelles Traditionnelles
Ruelles Vertes
Segments des Rues

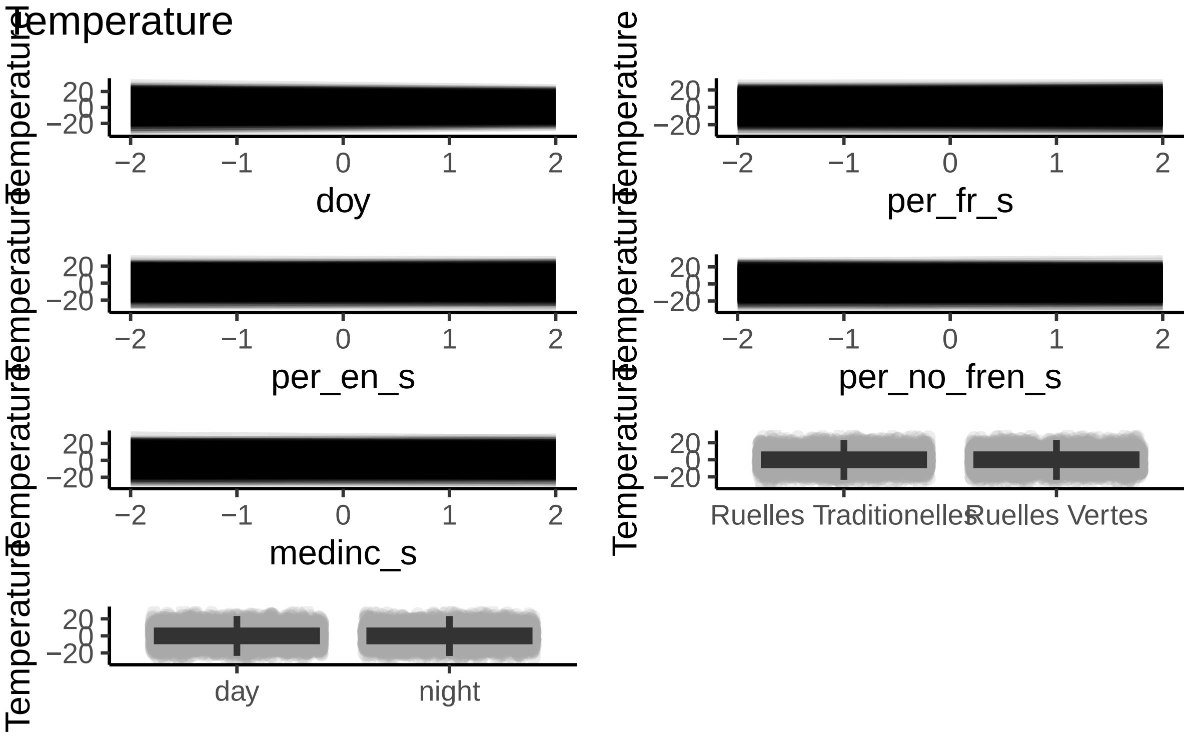
Temperature
Temperature
20
0
−20
−2
−1
0
1
2
doy
Temperature
per_fr_s
per_en_s
per_no_fren_s
medinc_s
Ruelles Traditionelles
Ruelles Vertes
day
night

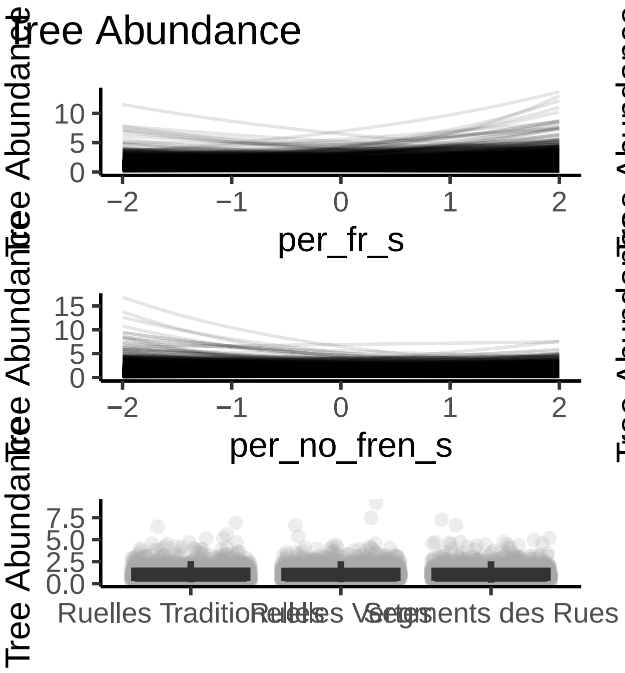
Tree Abundance
Tree Abundance
10
5
0
−2
−1
0
1
2
per_fr_s
Tree Abundance
9
6
3
0
per_en_s
Tree Abundance
15
10
5
0
per_no_fren_s
Tree Abundance
medinc_s
Tree Abundance
7.5
5.0
2.5
0.0
Ruelles Traditionnelles
Ruelles Vertes
Segments des Rues

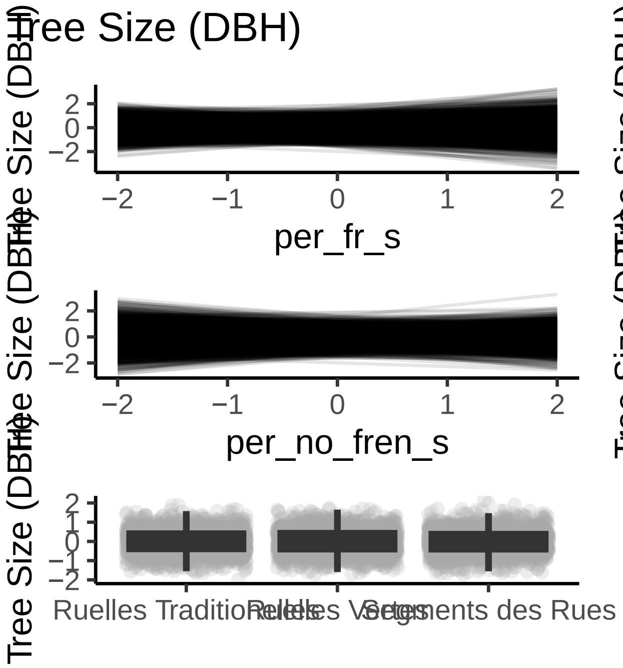
Tree Size (DBH)
Tree Size (DBH)
per_fr_s
Tree Size (DBH)
per_en_s
Tree Size (DBH)
per_no_fren_s
Tree Size (DBH)
medinc_s
Tree Size (DBH)
Ruelles Traditionelles
Ruelles Vertes
Segments des Rues

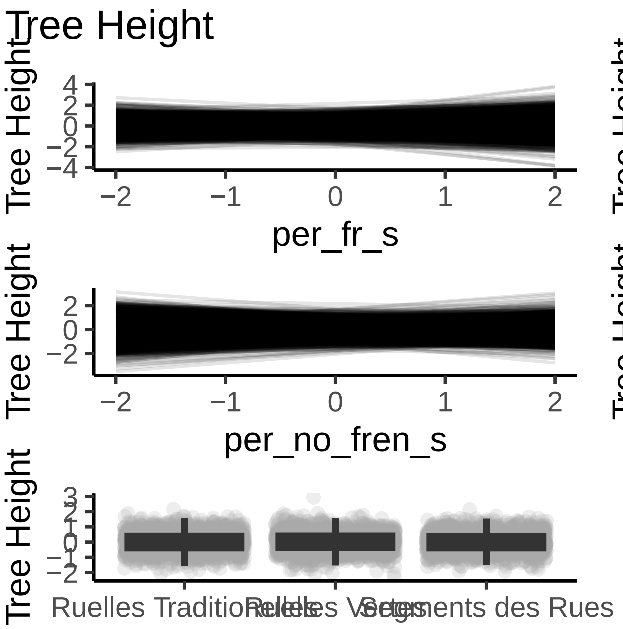
Tree Height
Tree Height
per_fr_s
Tree Height
per_en_s
Tree Height
per_no_fren_s
Tree Height
medinc_s
Tree Height
Ruelles Traditionnelles
Ruelles Vertes
Segments des Rues

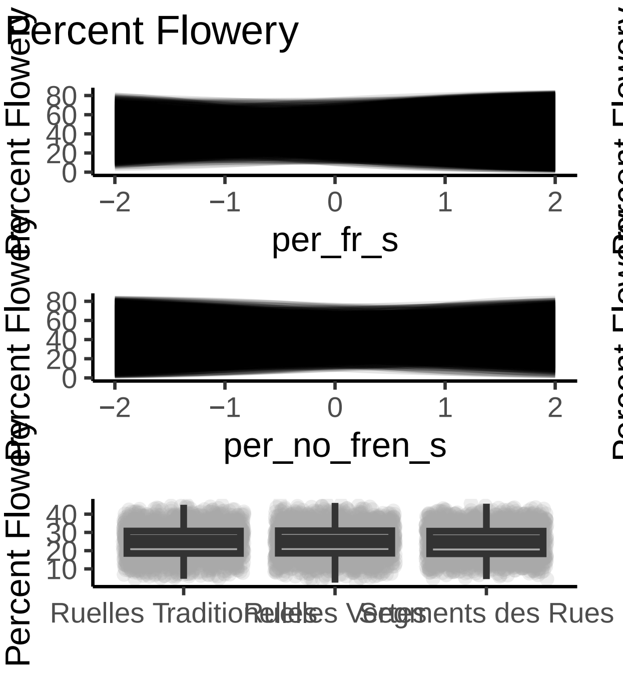
Percent Flowery
Percent Flowery
per_fr_s
per_en_s
per_no_fren_s
medinc_s
Ruelles Traditionelles
Ruelles Vertes
Segments des Rues

**Trois-Rivieres**

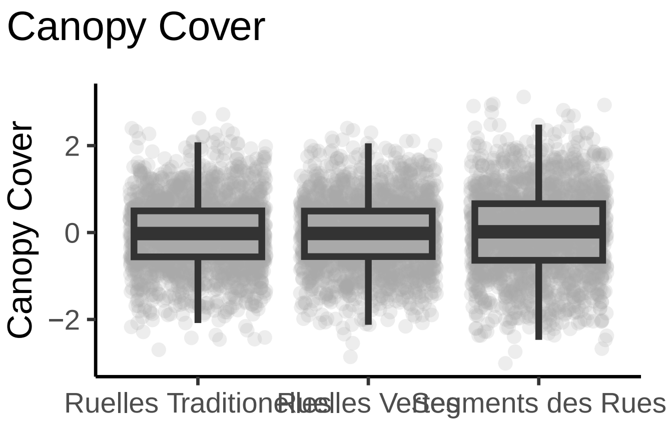

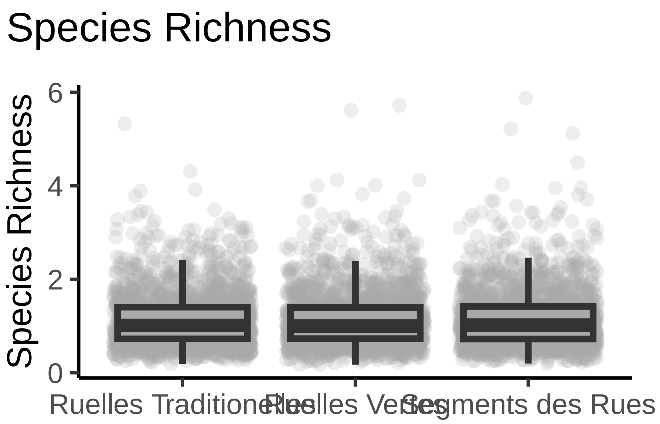

Species Richness
Species Richness
6
4
2
0
Ruelles Traditionelles
Ruelles Vertes
Segments des Rues

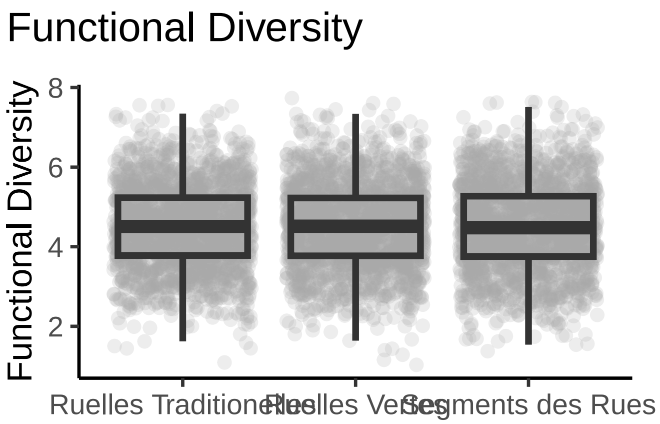
Functional Diversity
Functional Diversity
8
6
4
2
Ruelles Traditionelles
Ruelles Vertes
Segments des Rues

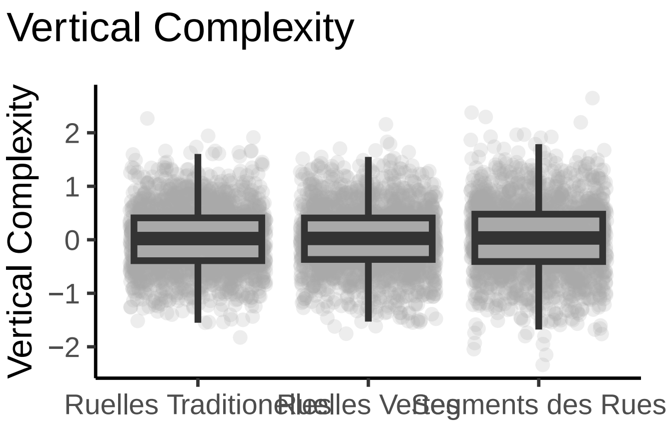
Vertical Complexity
Vertical Complexity
2
1
0
−1
−2
Segments des Rues

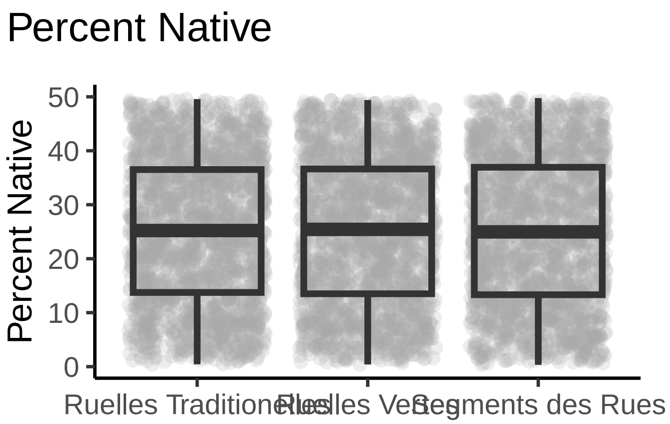
Percent Native
Percent Native
50
40
30
20
10
0
Segments des Rues

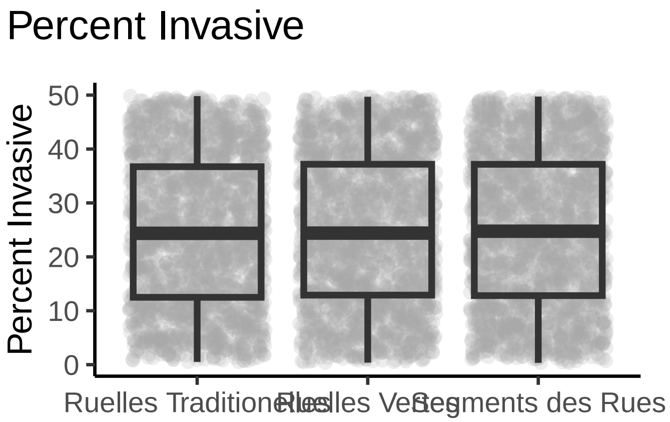
Percent Invasive
Percent Invasive
50
40
30
20
10
0
Ruelles Traditionelles
Ruelles Vertes
Segments des Rues

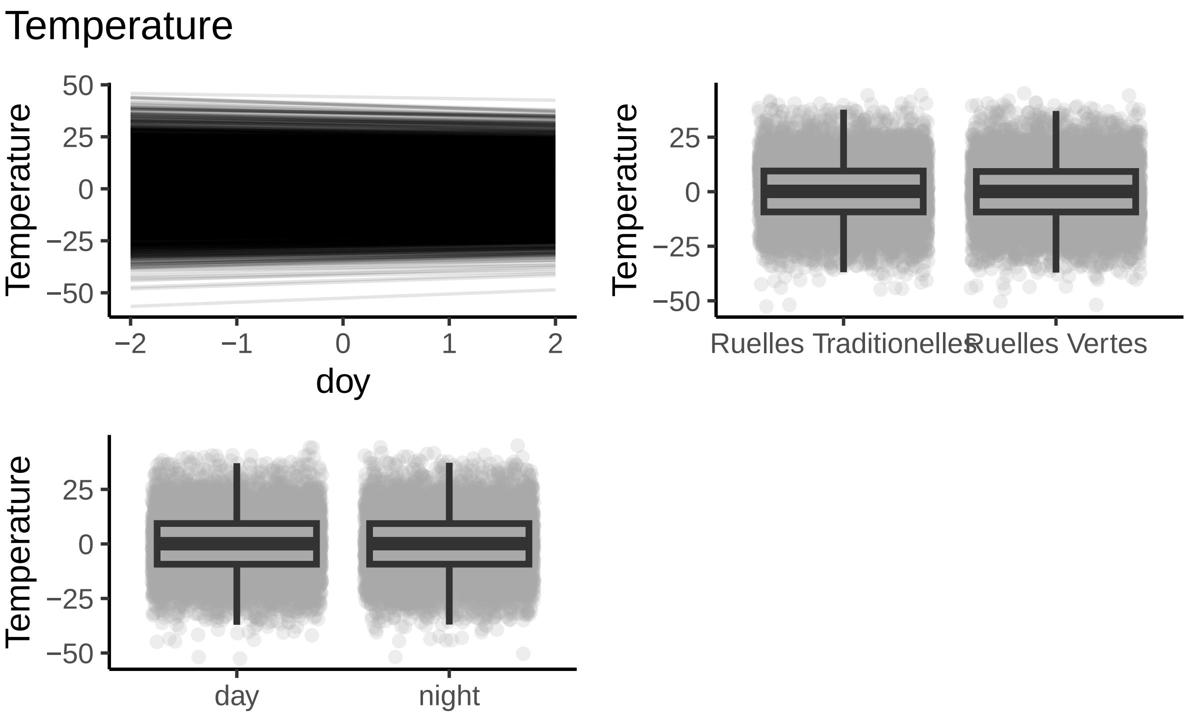
Temperature
Temperature
50
25
0
−25
−50
−2
−1
0
1
2
doy
Temperature
25
0
−25
−50
Ruelles Traditionelles
Ruelles Vertes
Temperature
25
0
−25
−50
day
night

Tree Abundance

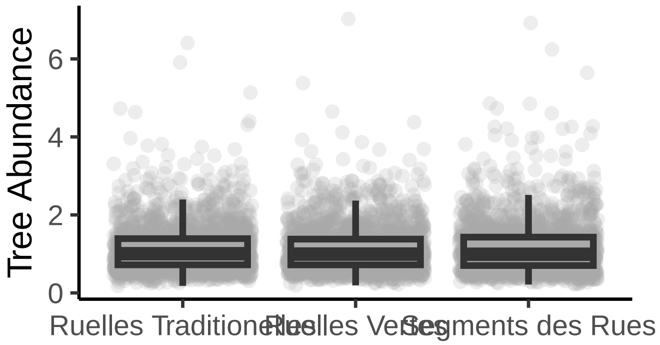

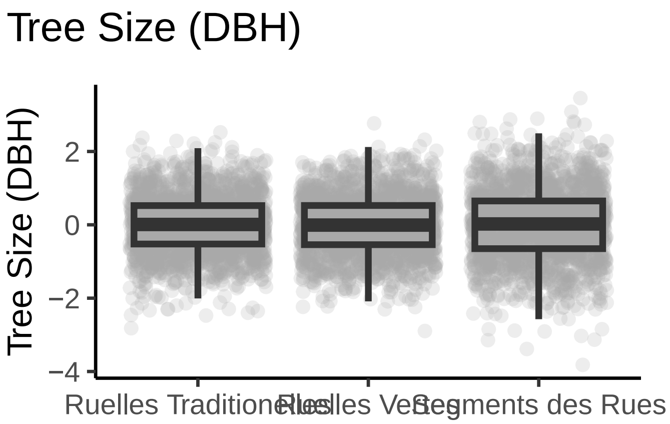
Tree Size (DBH)
Tree Size (DBH)
2
0
−2
−4
Segments des Rues

Tree Height

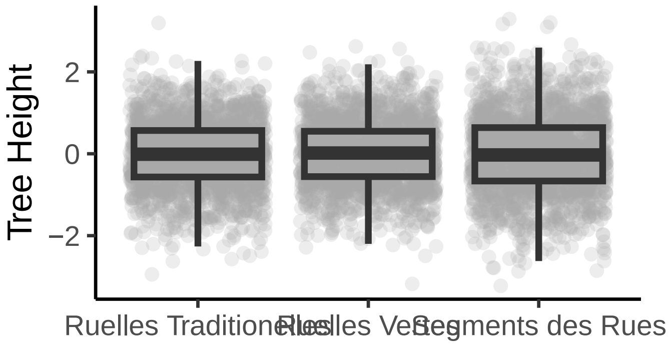

Tree Height
2
0
−2
Ruelles Traditionelles
Ruelles Vertes
Segments des Rues

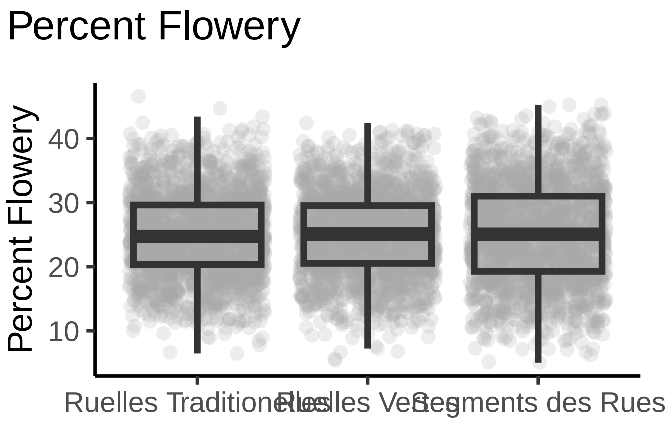

#### F) Model Diagnostics

These model diagnostic plots assess whether the chains of our models are converged and well mixed, and if the model is well specified and has an adequate fit.

The first plot of the series shows trace plots for each of our parameters, where we want to see stationary and well-mixed chains. The second plot shows an autocorrelation plot by chain and parameter. We want our autocorrelation to quickly drop to zero with increasing lag. Thirdly, the Rhat plot monitors whether a chain has converged to the equilibrium distribution, if all chains are at equilibrium Rhat will be one. If chains have not converged, Rhat will be greater than 1. The fourth plot is the ratio between effective sample size (Neff) and total sample size (N).Because the draws within a Markov chain are not independent if there is autocorrelation, the effective sample size, neff, is usually smaller than the total sample size, N. The larger the ratio, the better. Finally, we have the posterior predictive check where we want the black line to be within/close to the blue lines, to indicate that our model is adequately generative.

### Villeray-Saint Michel-Parc Extension

Canopy Cover

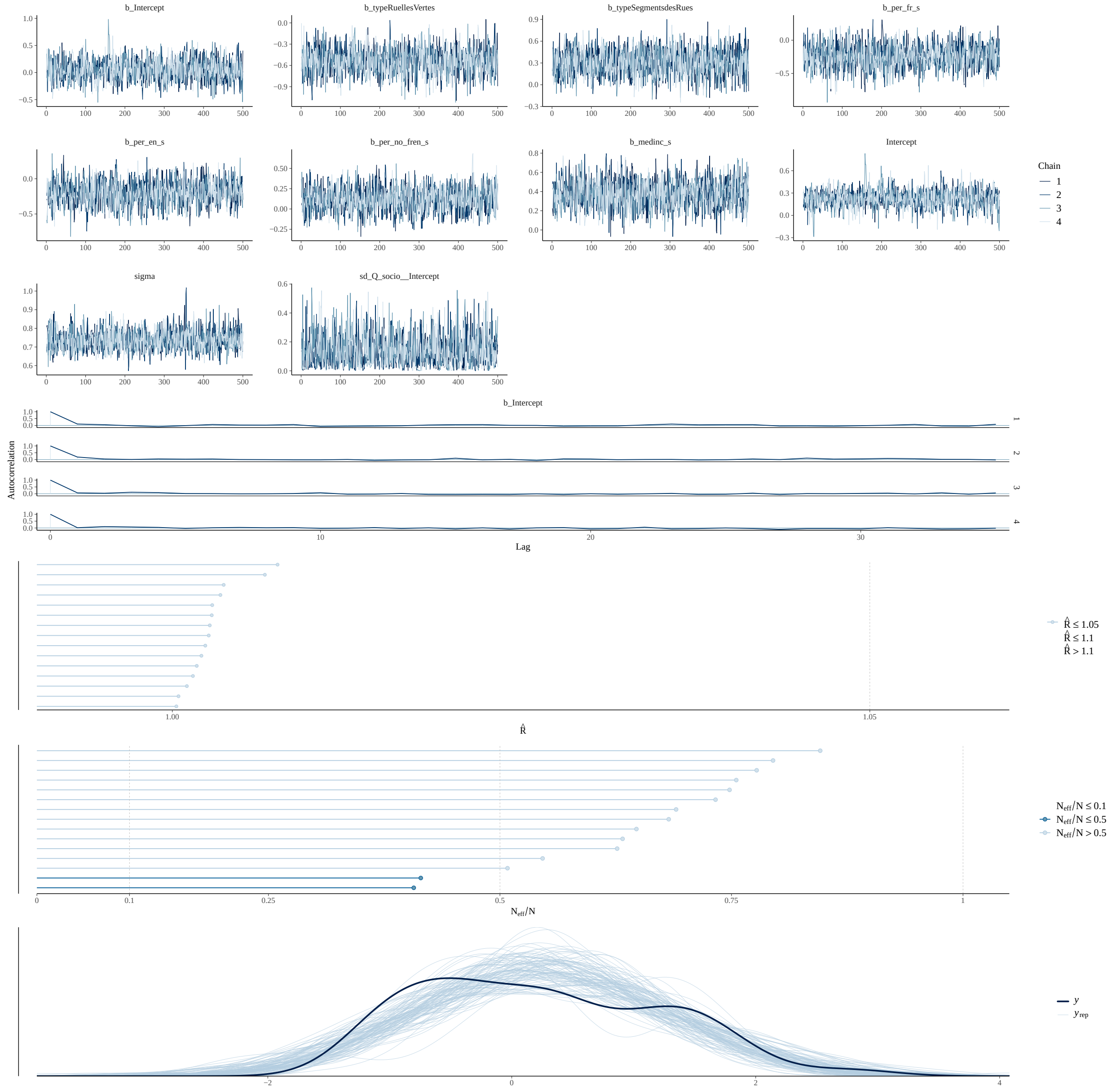

Firefly Abundance

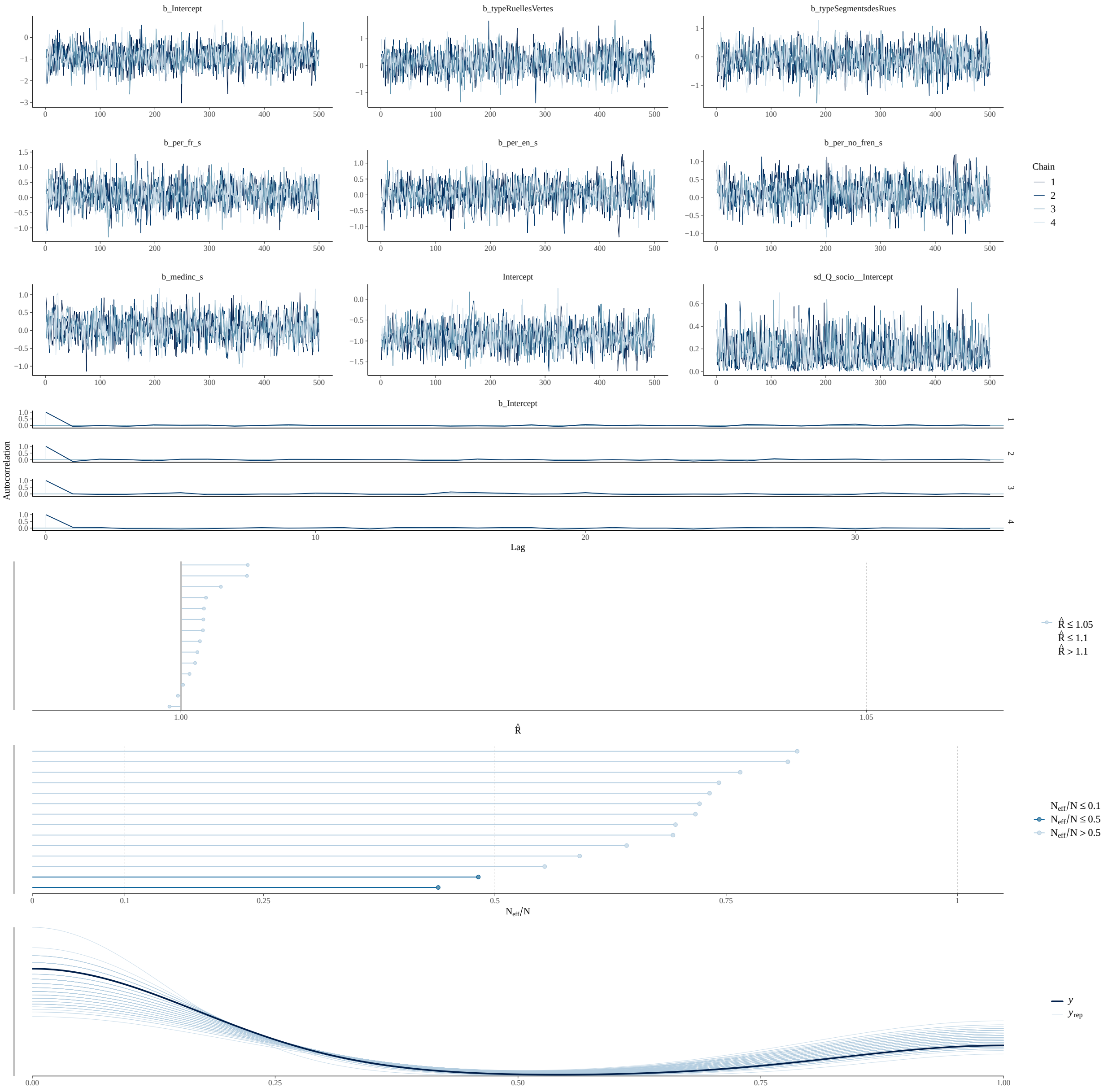

Species Richness

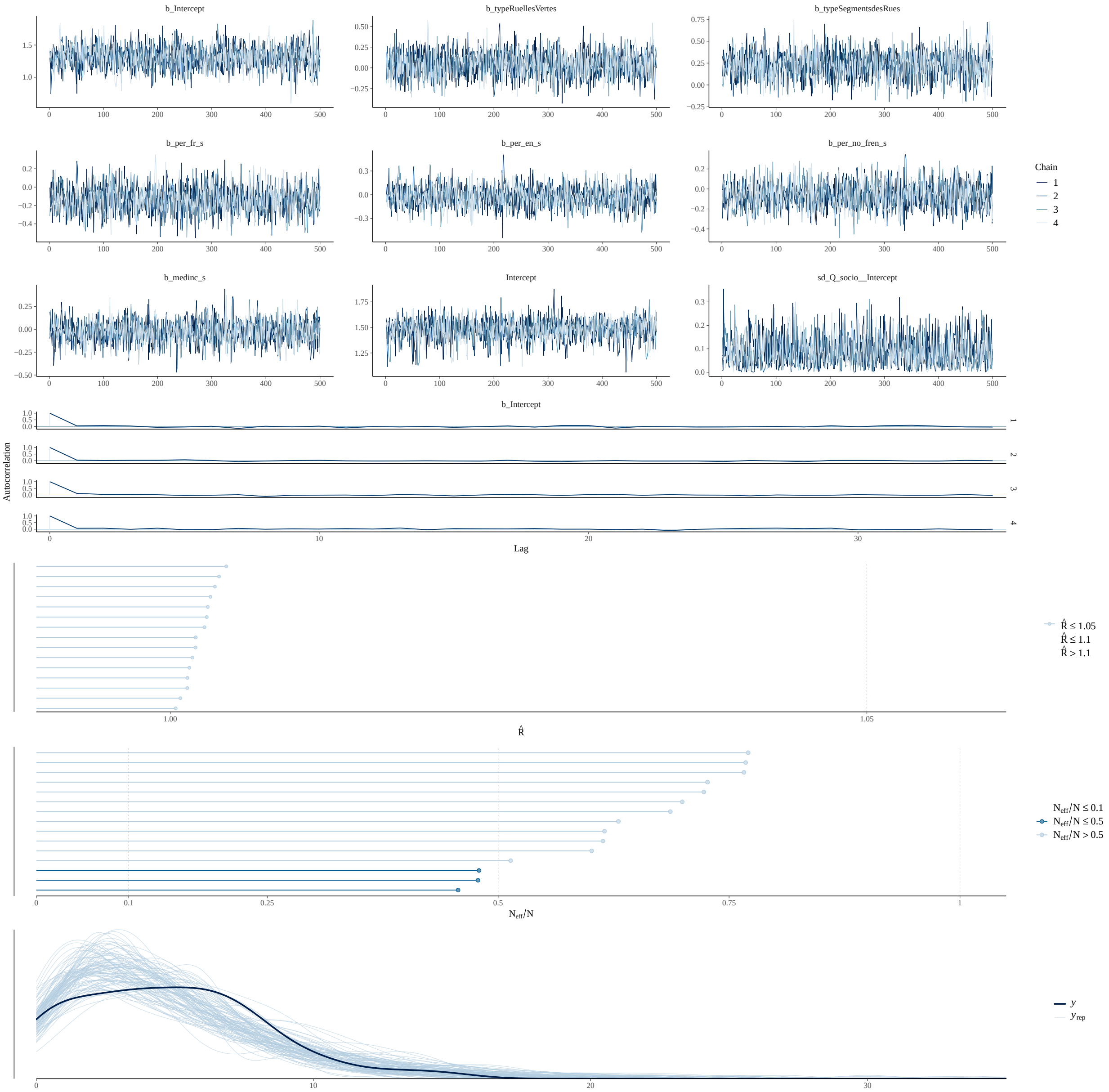

#### Functional Diversity

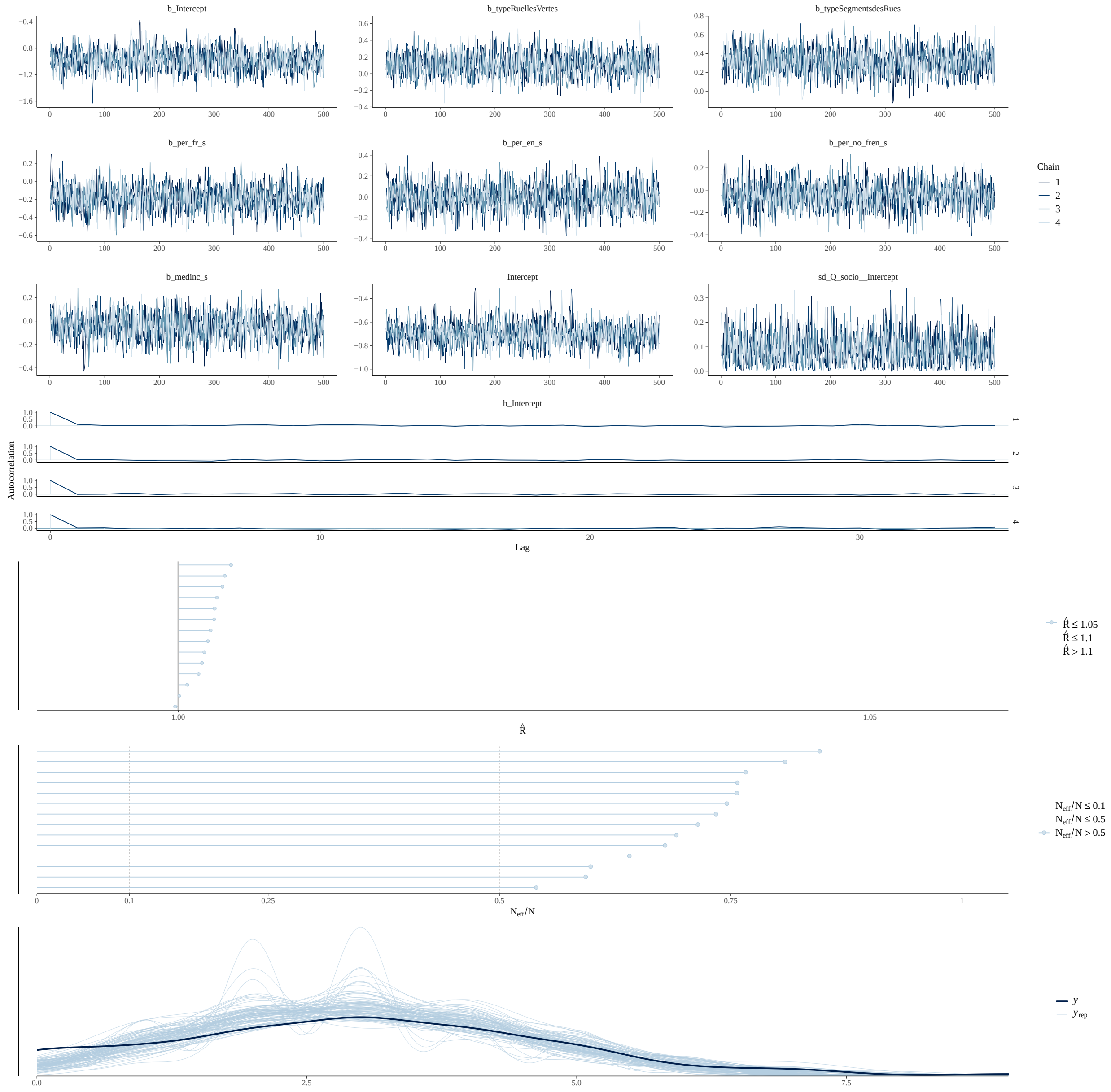

Vertical Complexity

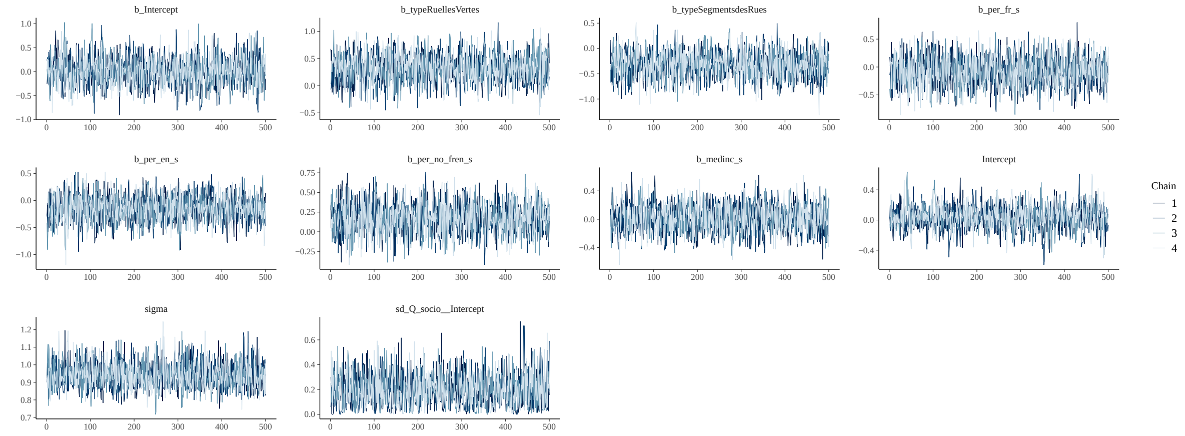

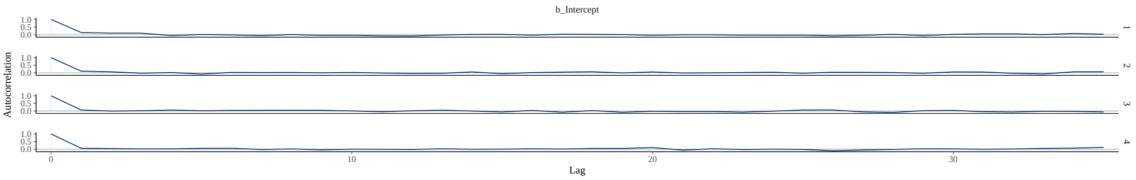

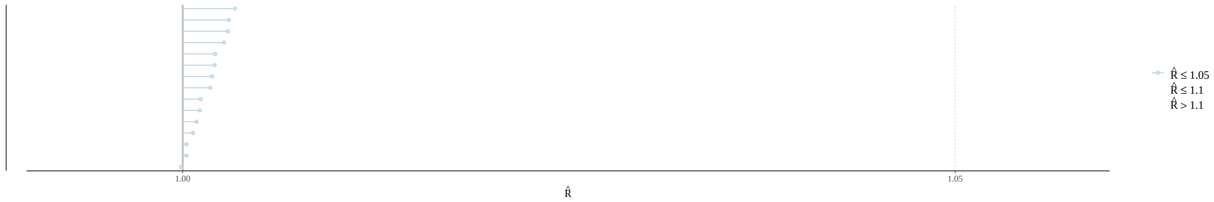

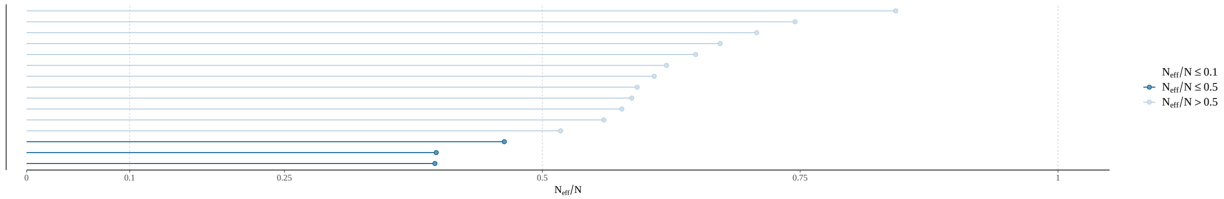

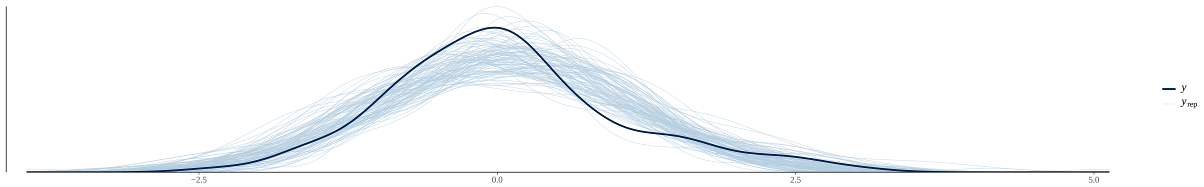

Percent Native

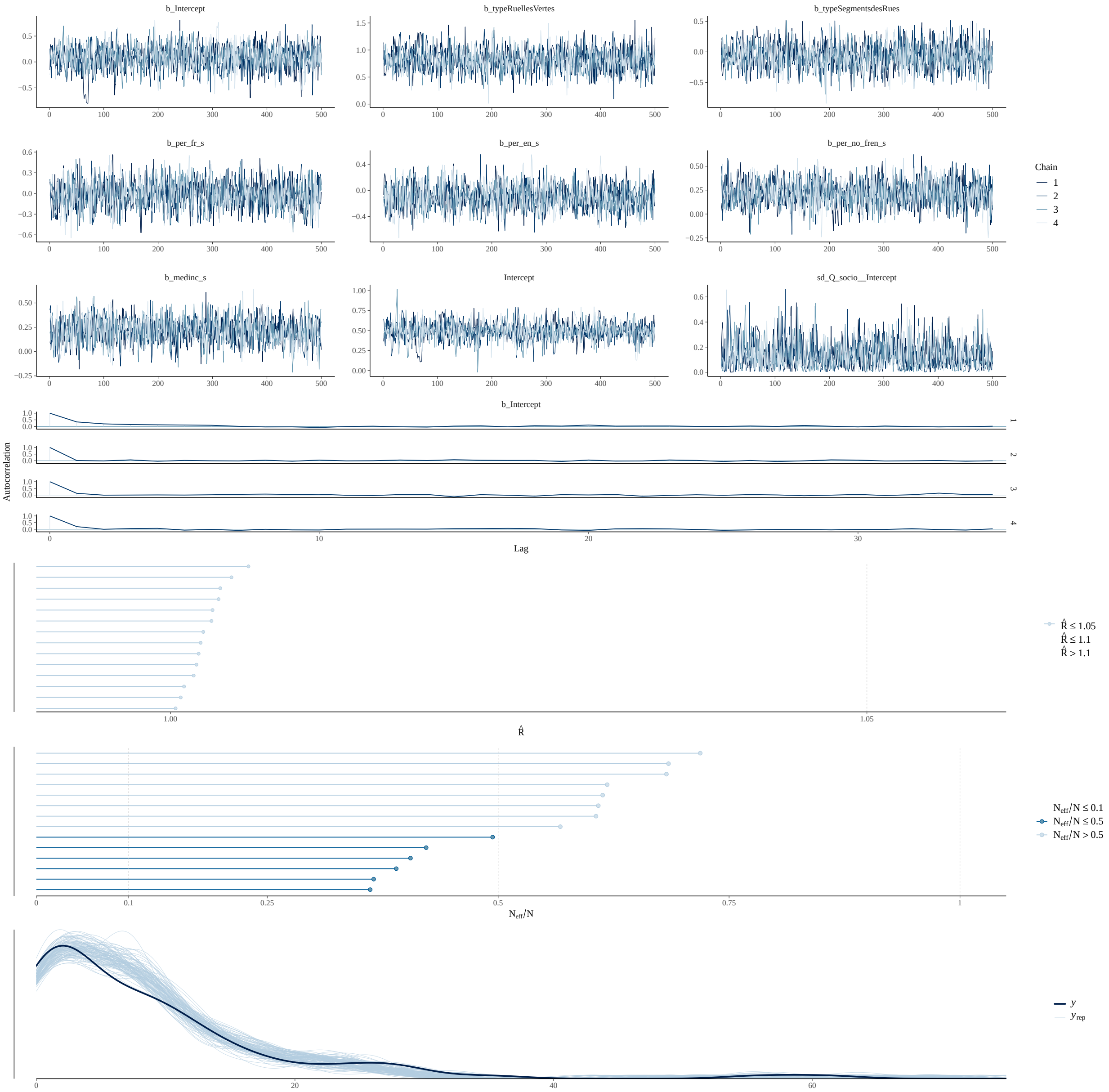

Percent Invasive

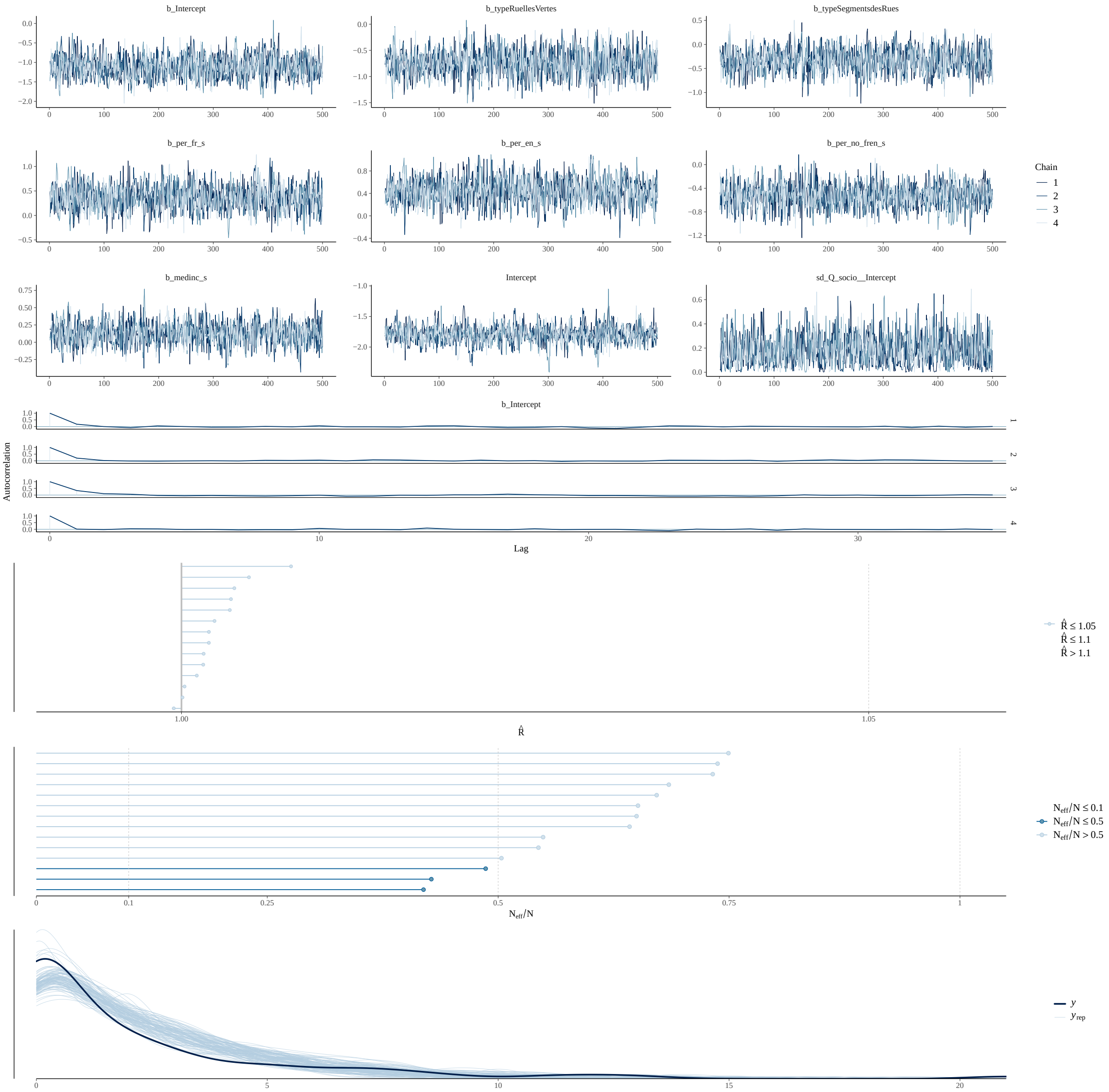

Temperature

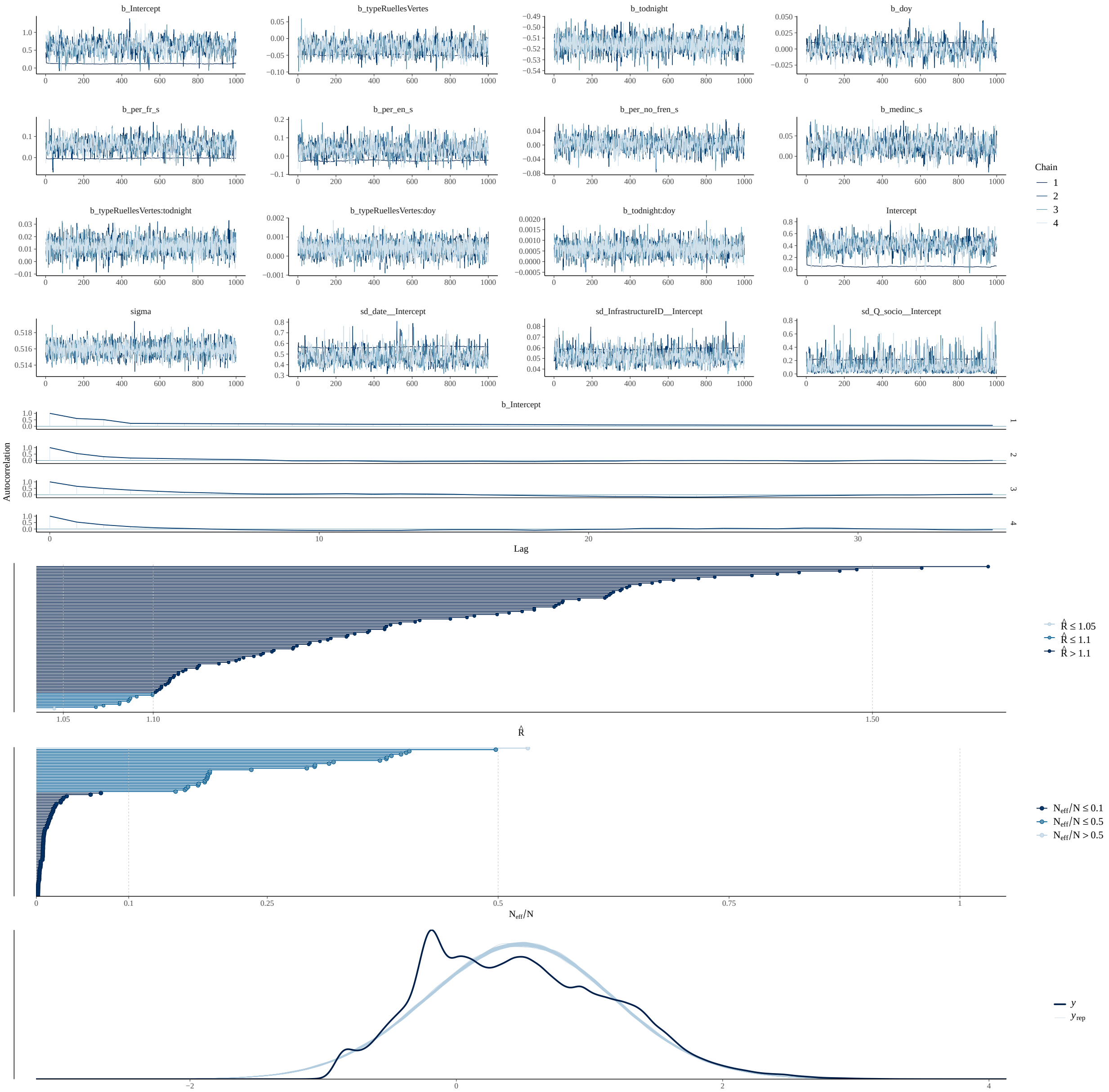

Tree Abundance

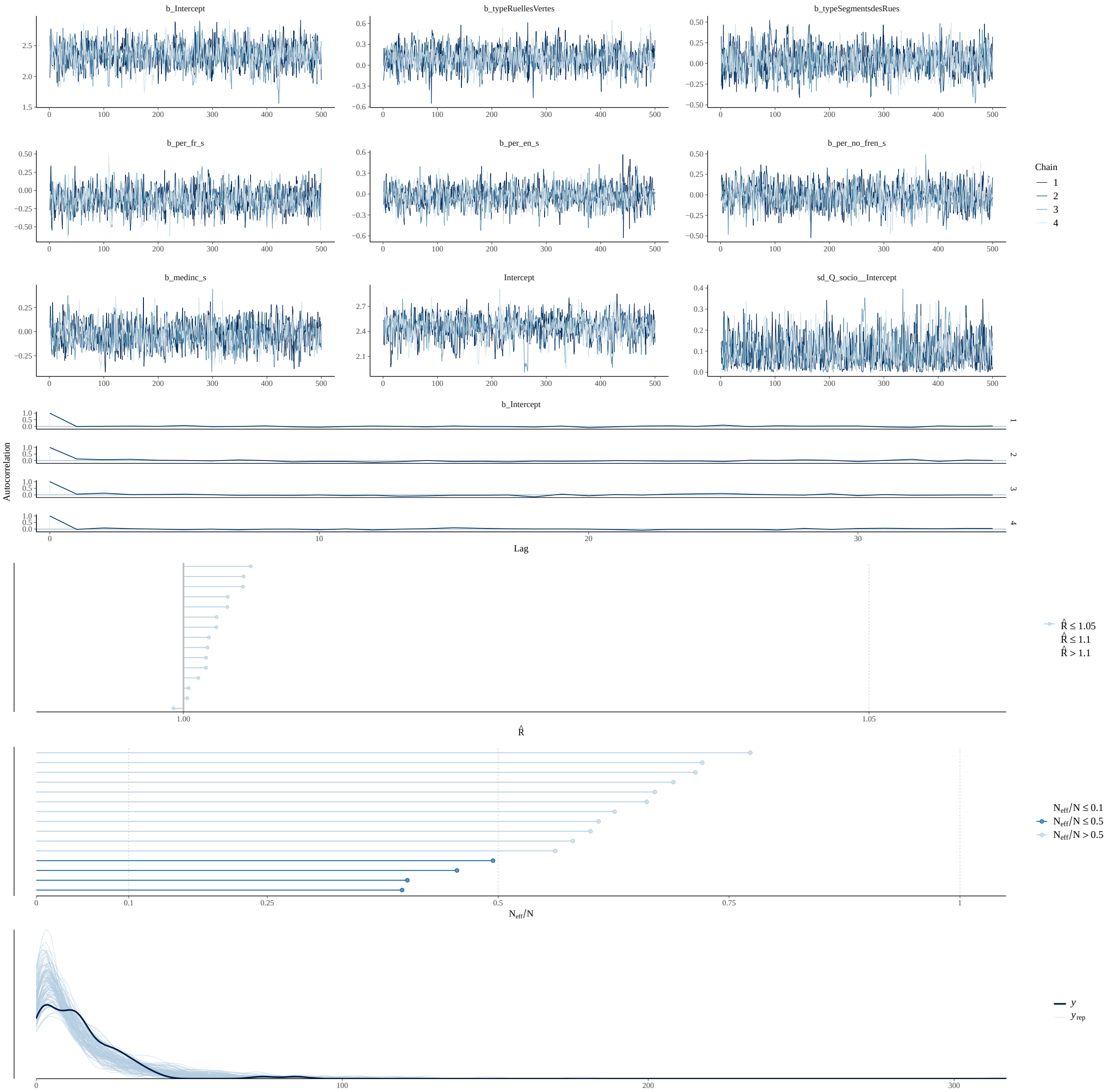

Tree Size (DBH)

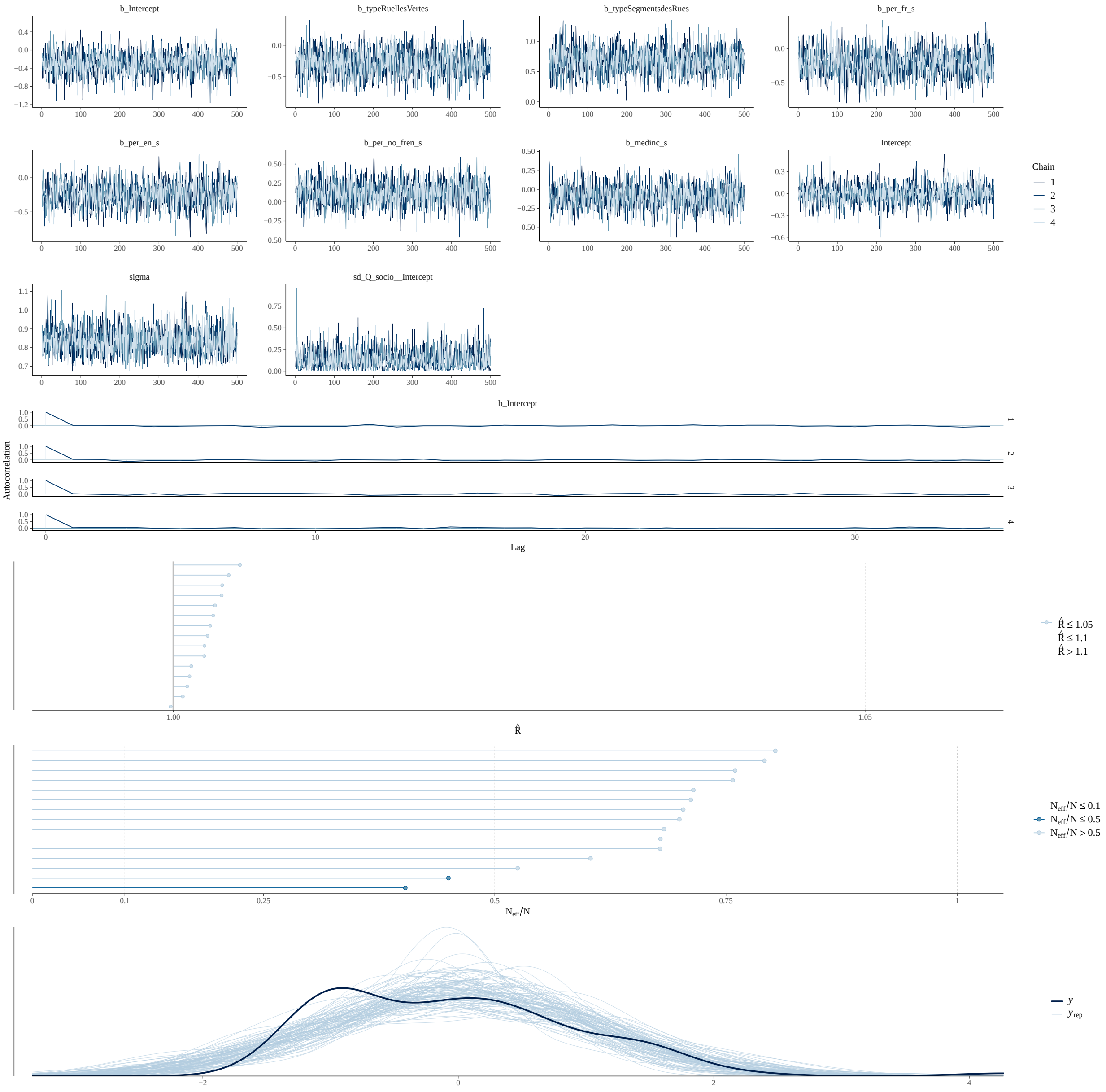

Tree Height

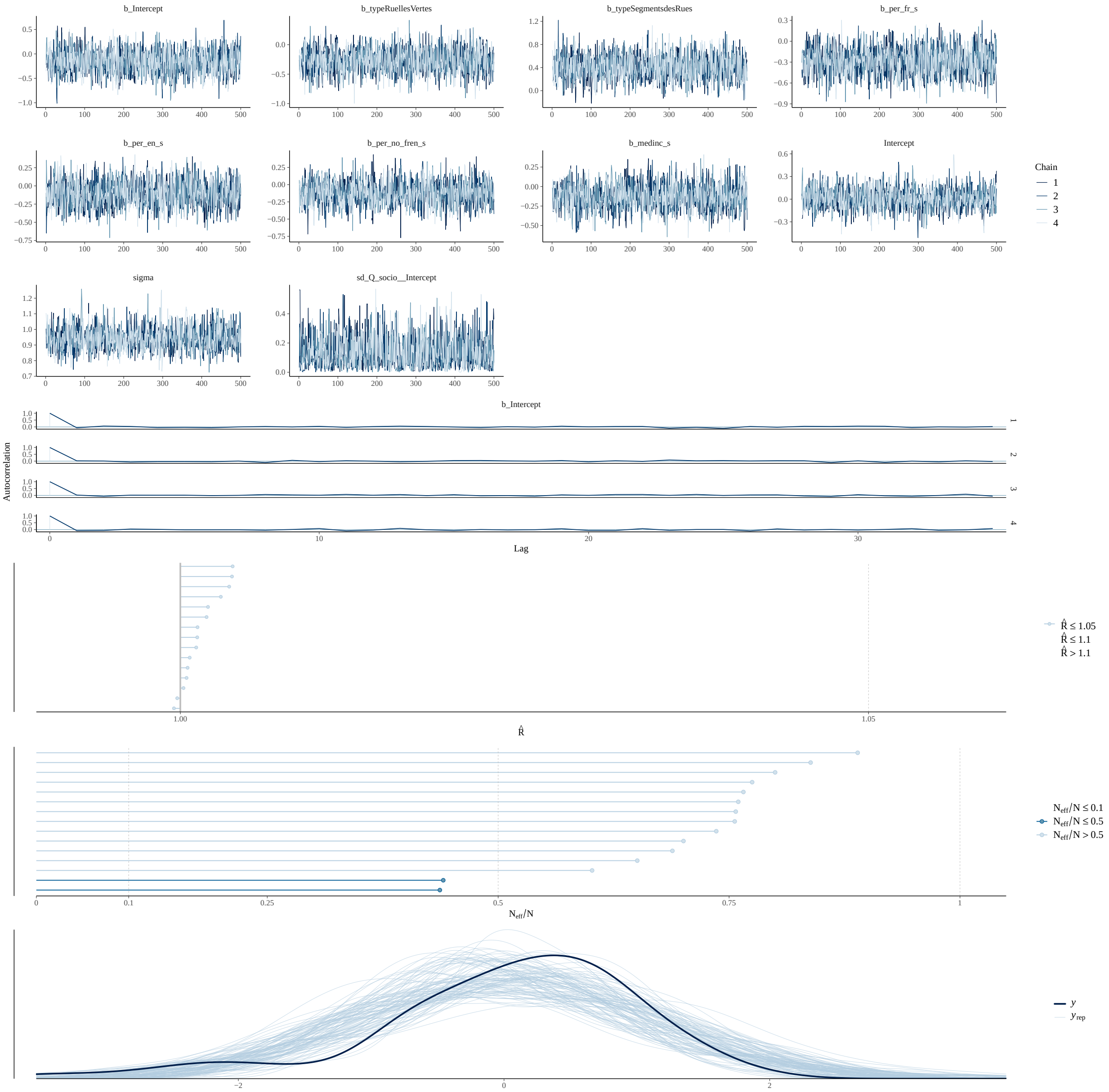
b_Intercept
b_typeRuellesVertes
b_typeSegmentsdesRues
b_per_fr_s
b_per_en_s
b_per_no_fren_s
b_medinc_s
Intercept
sigma
sd_Q_socio__Intercept
Chain
1
2
3
4
Autocorrelation
Lag
R̂ ≤ 1.05
R̂ ≤ 1.1
R̂ > 1.1
R̂
N_eff/N ≤ 0.1
N_eff/N ≤ 0.5
N_eff/N > 0.5
N_eff/N
y
y_rep

Percent Flowery

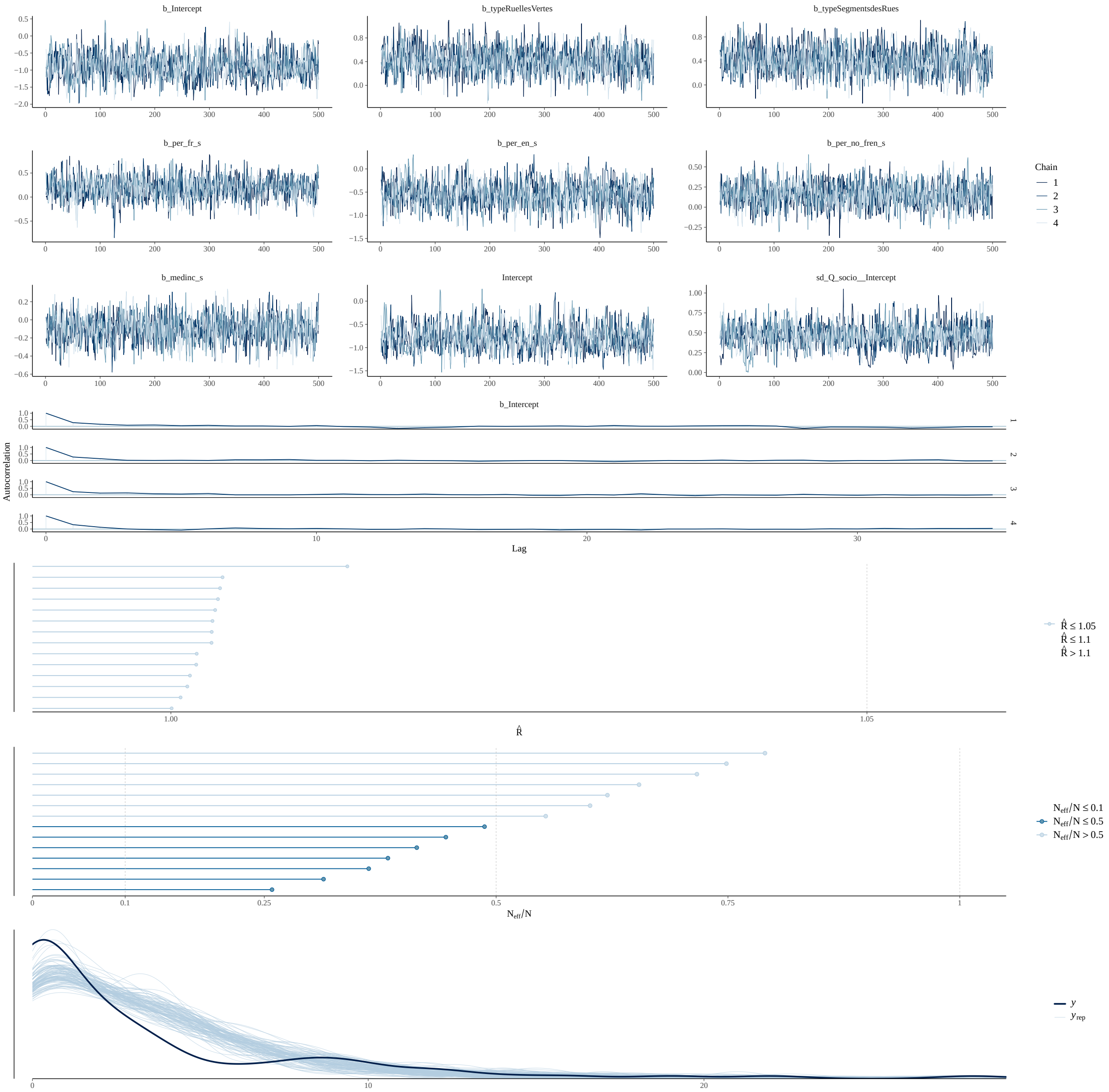

### Trois-Rivieres

#### Canopy Cover

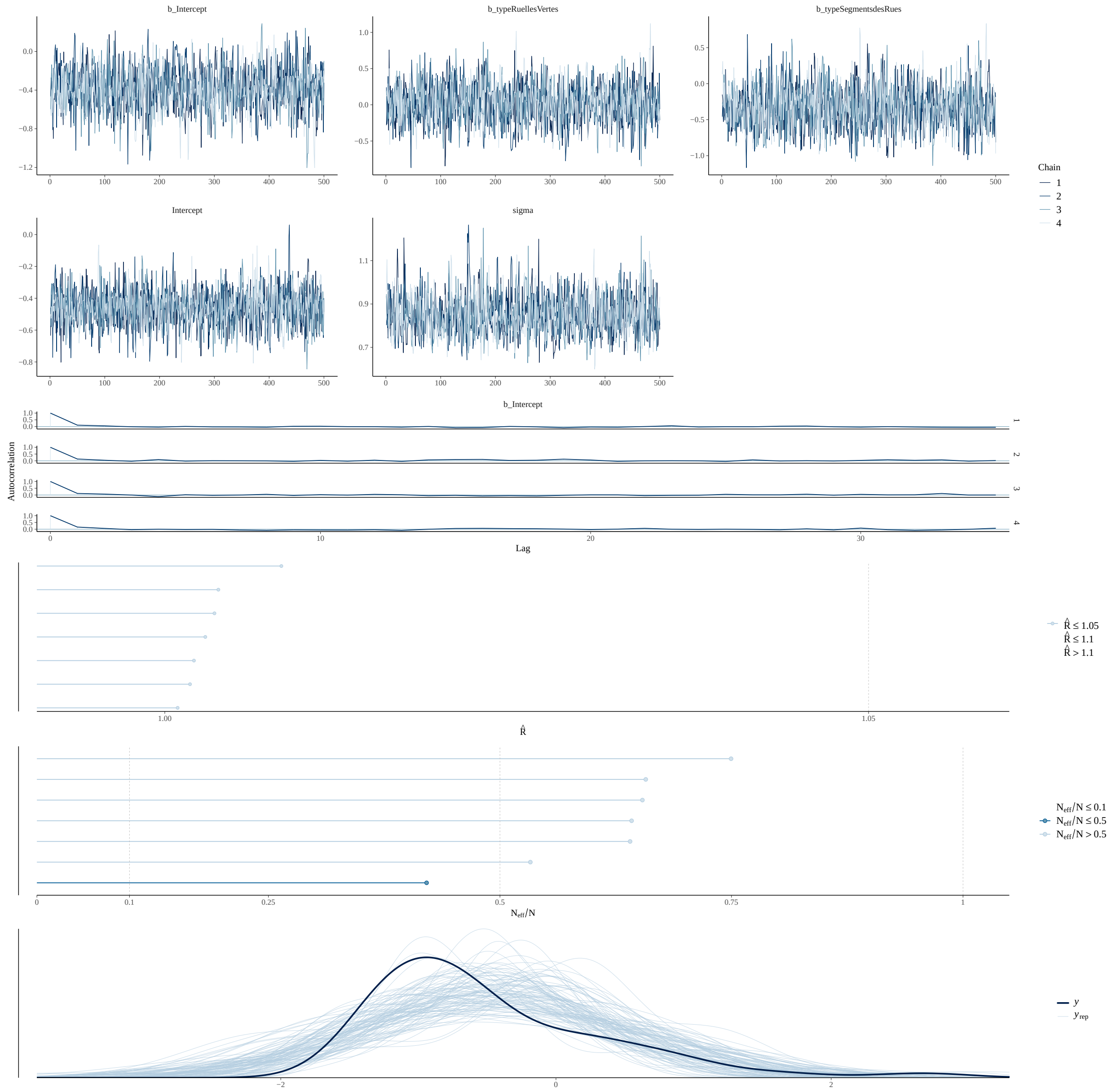

Species Richness

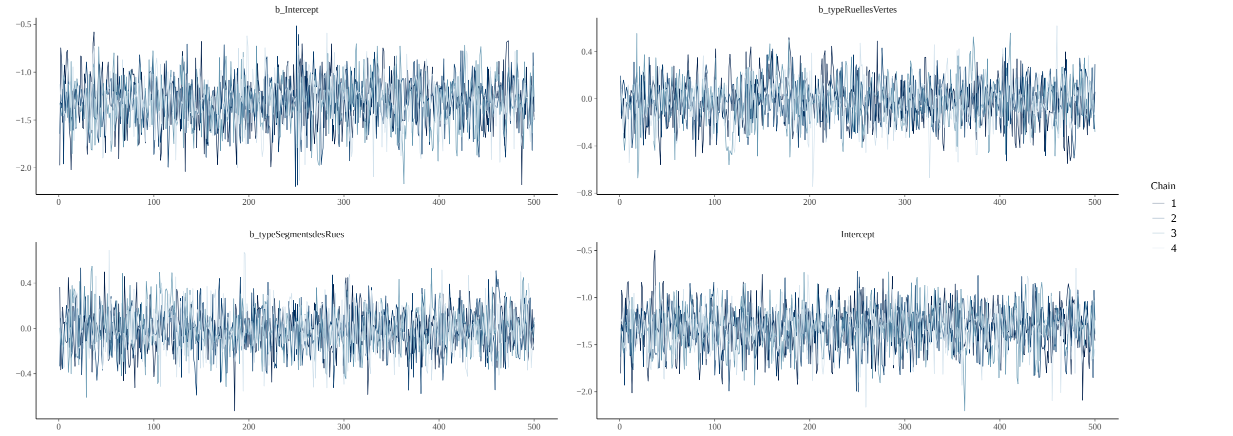

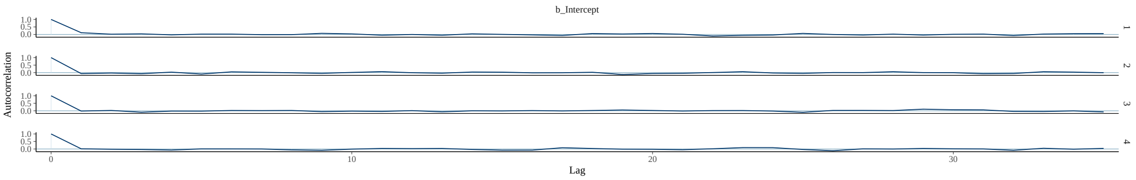

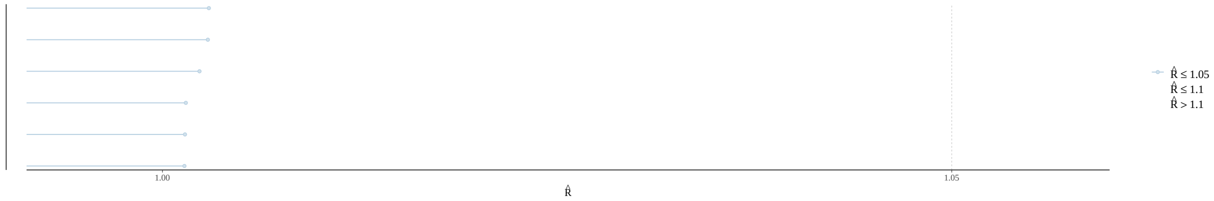

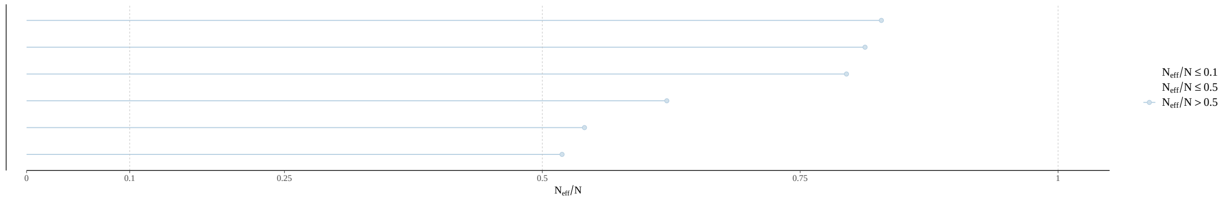

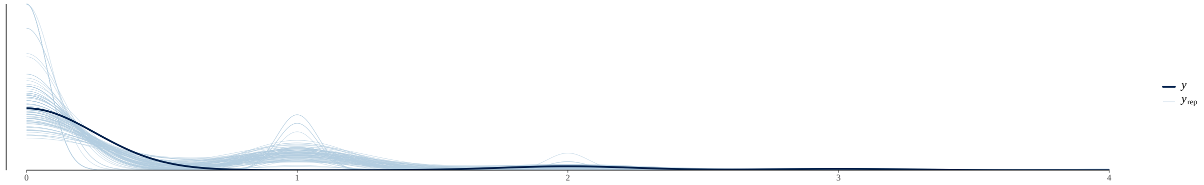

Functional Diversity

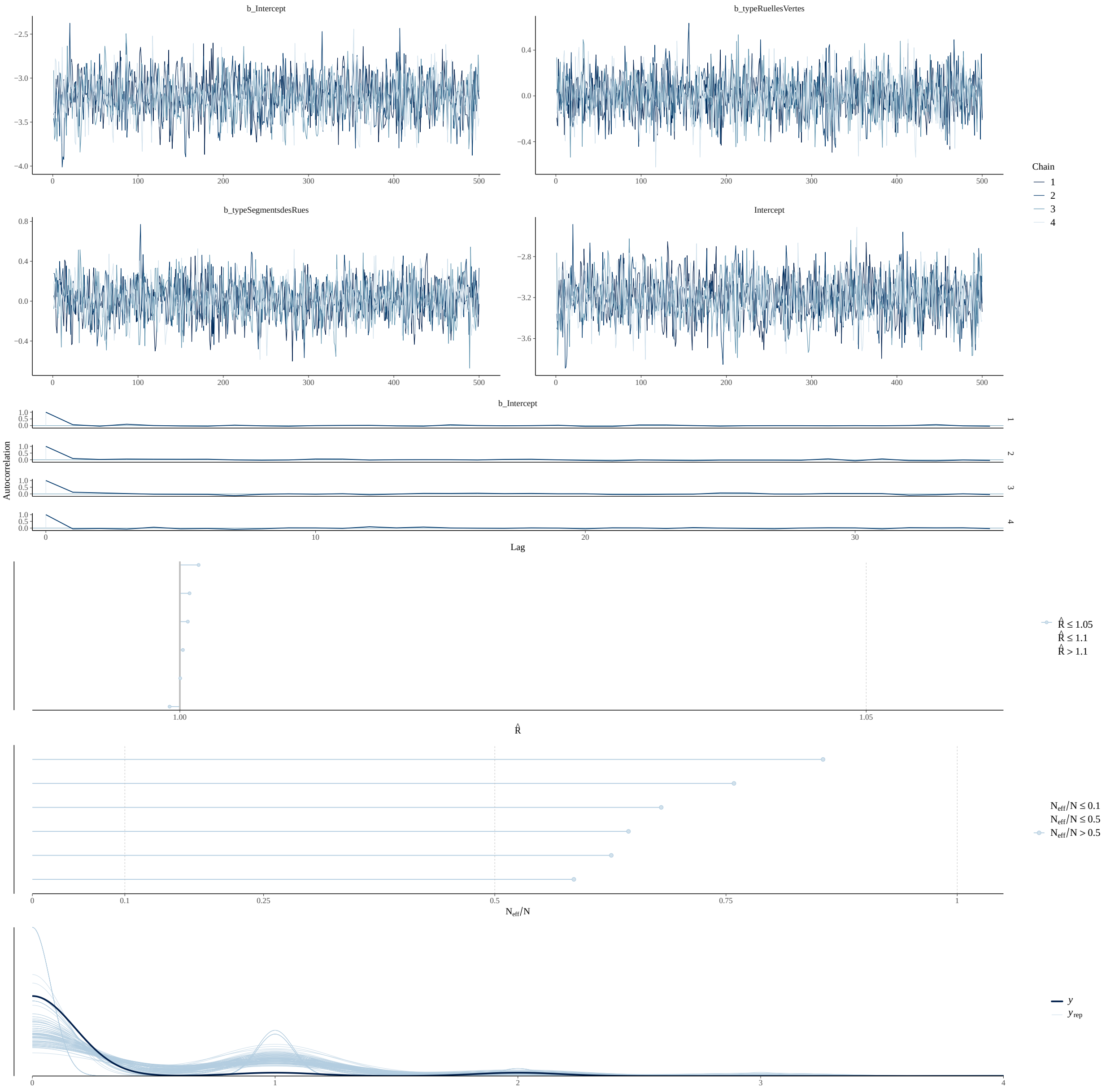

Vertical Complexity

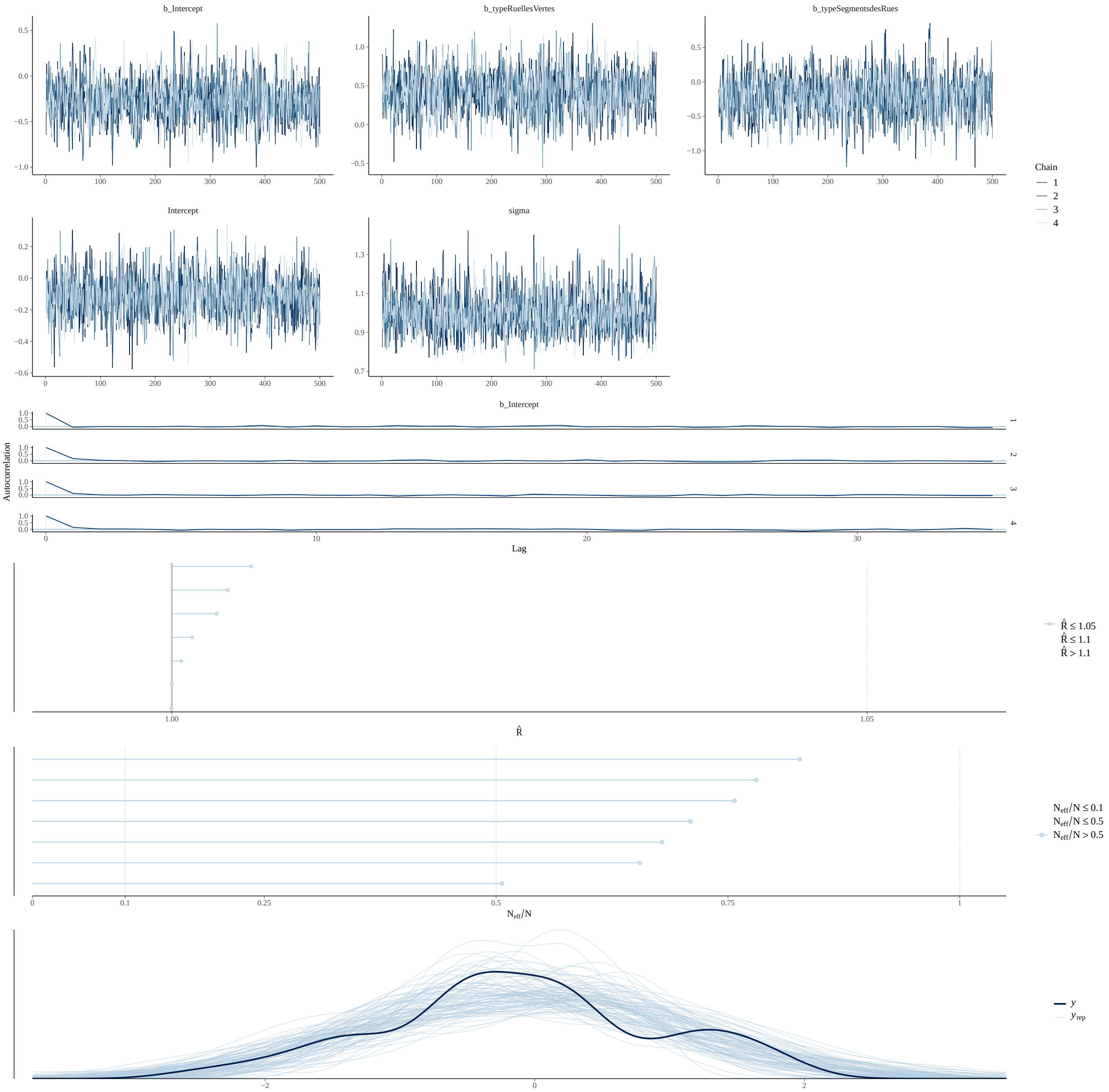

Percent Native

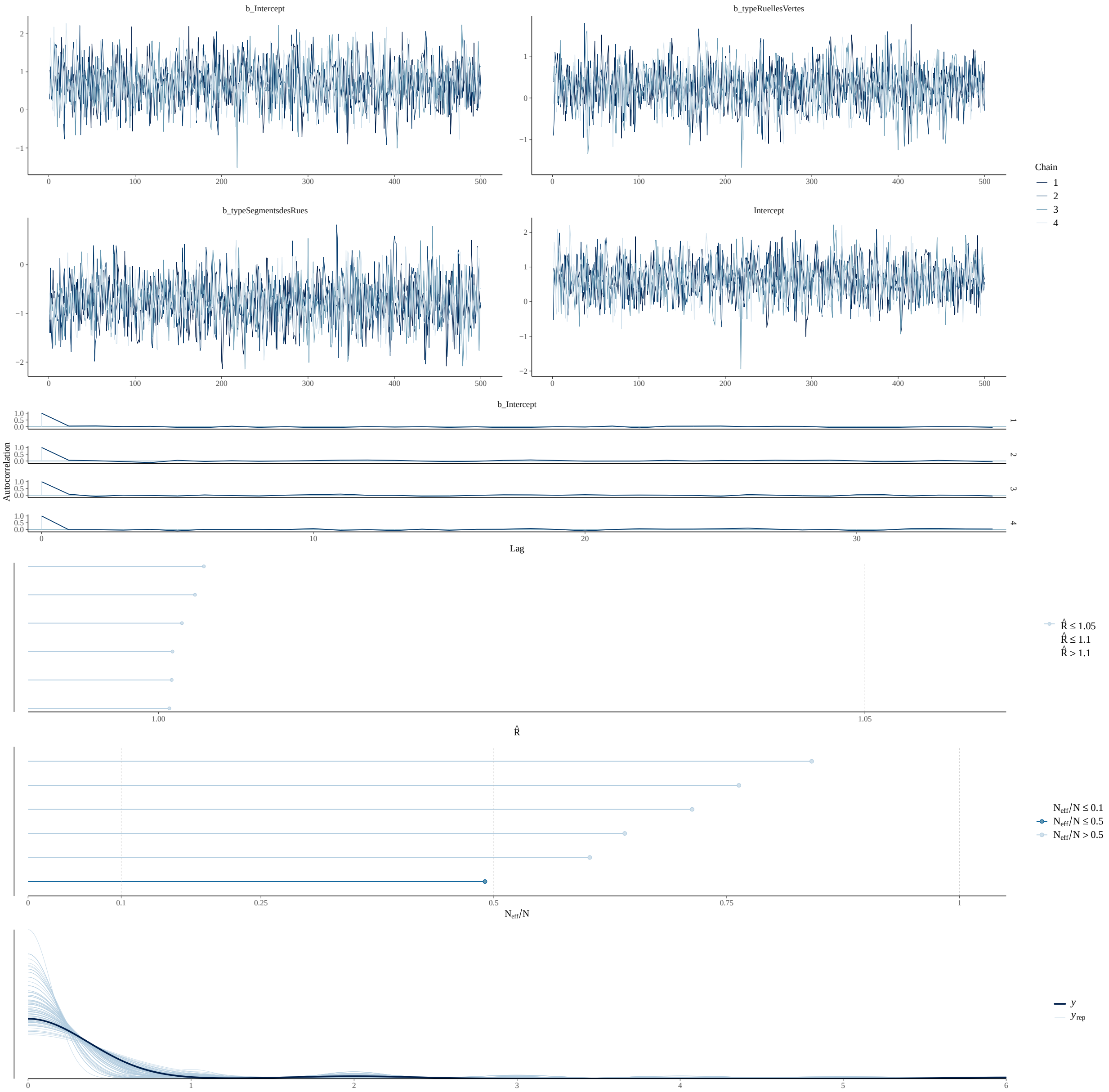

Percent Invasive

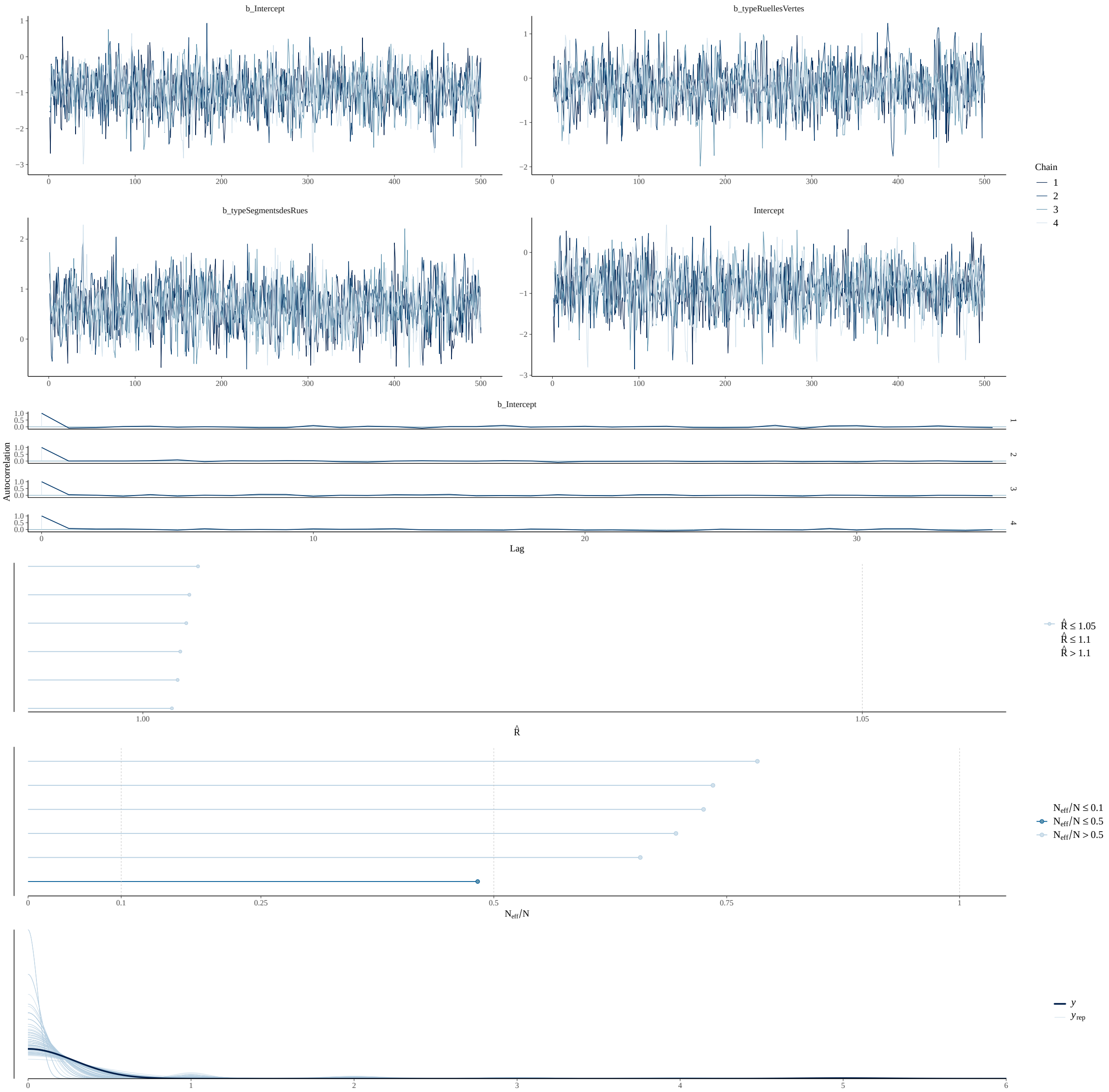

#### Temperature

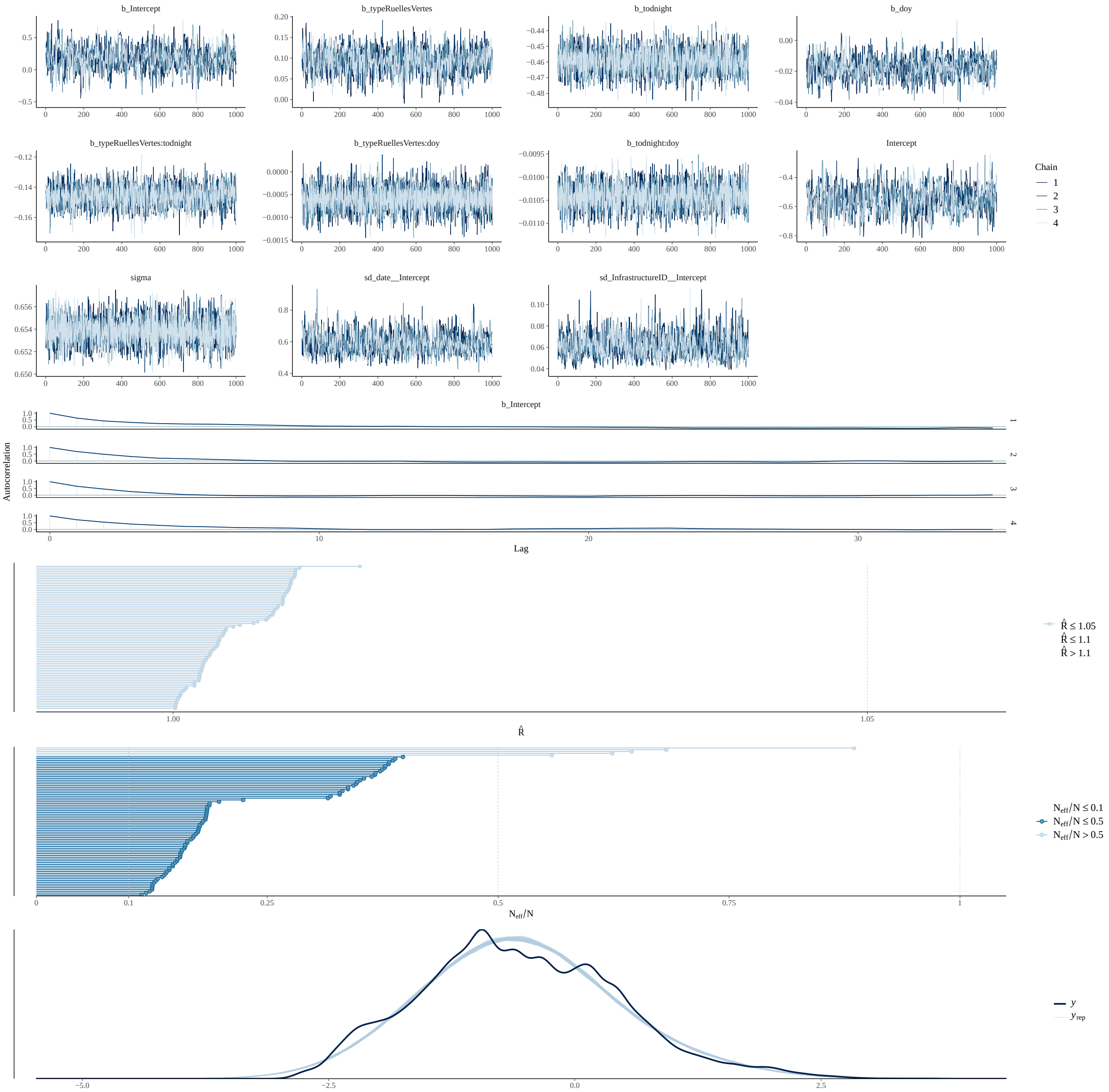

Tree Abundance

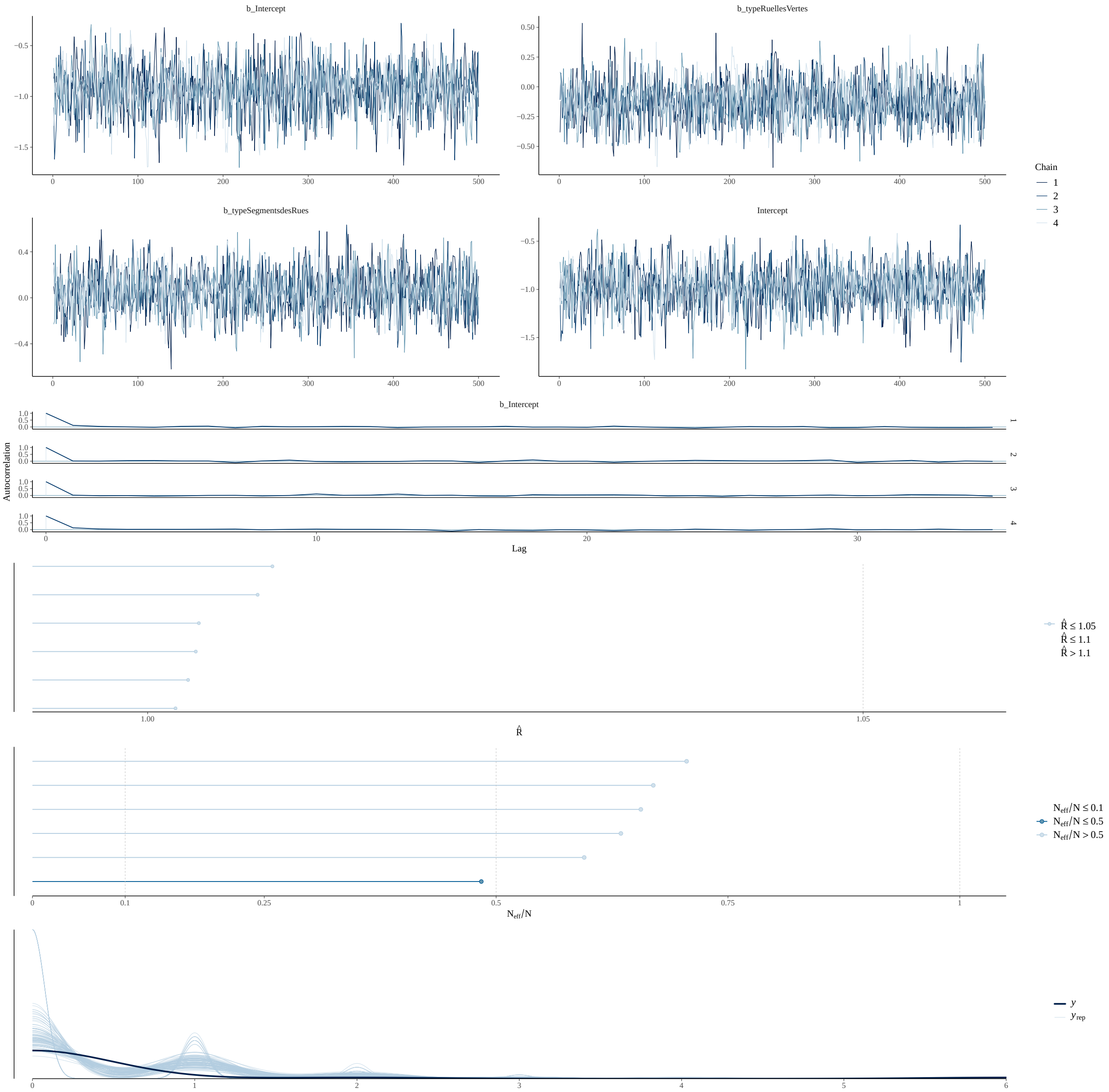

Tree Size (DBH)

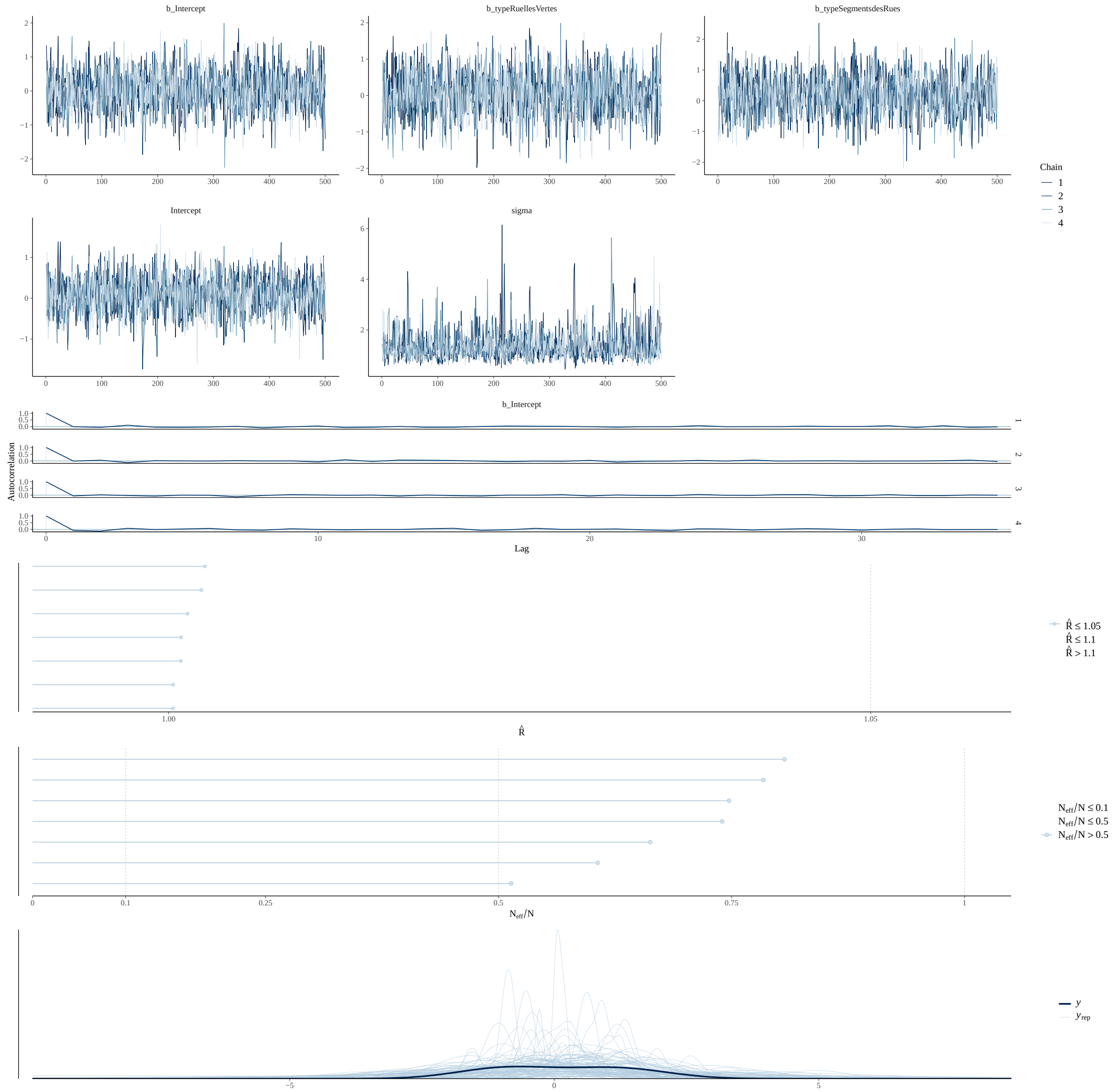

#### Tree Height

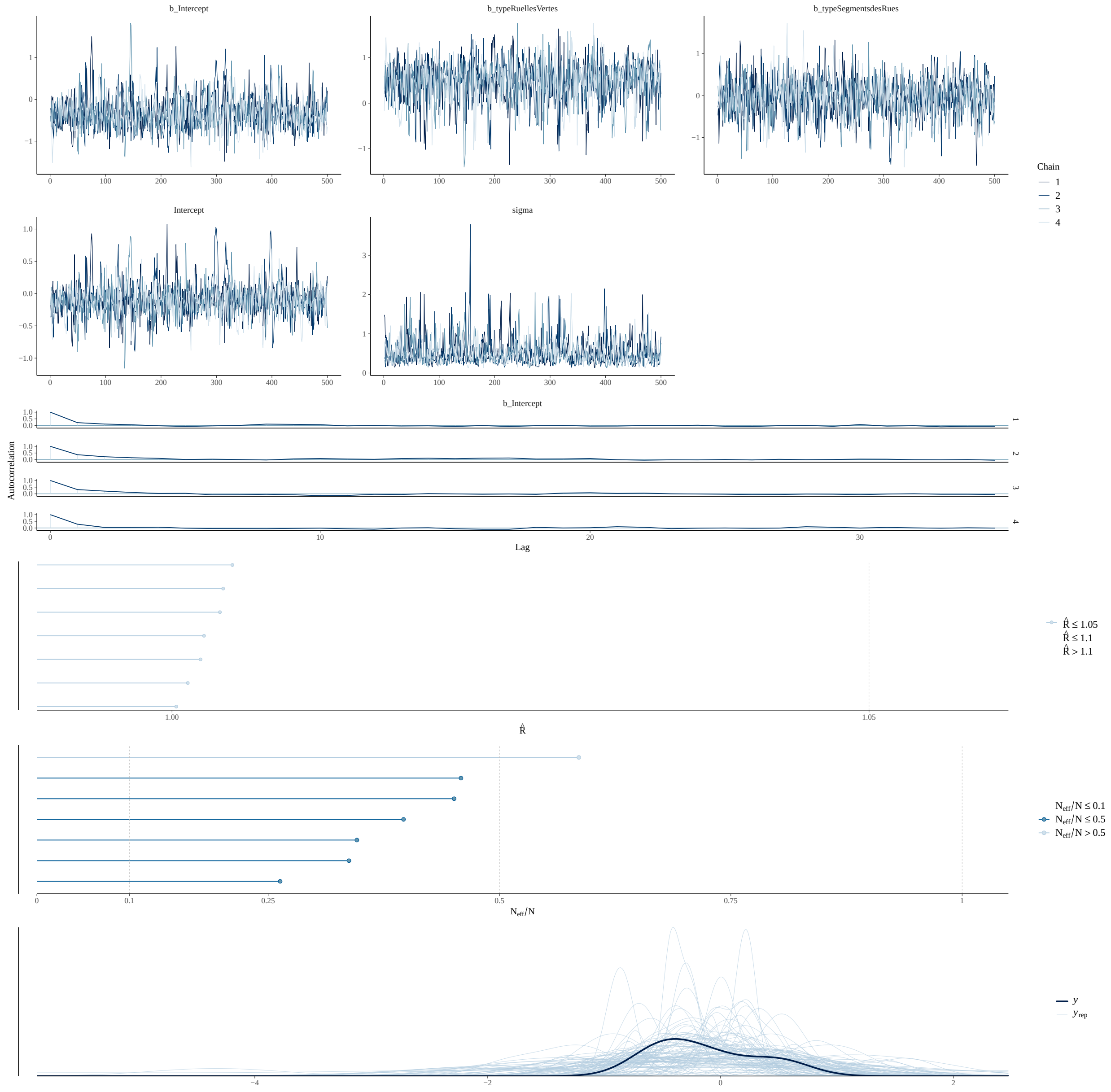
b_Intercept
b_typeRuellesVertes
b_typeSegmentsdesRues
Intercept
sigma
Chain
1
2
3
4
Autocorrelation
Lag
R̂ ≤ 1.05
R̂ ≤ 1.1
R̂ > 1.1
N_eff/N ≤ 0.1
N_eff/N ≤ 0.5
N_eff/N > 0.5
y
y_rep

Percent Flowery

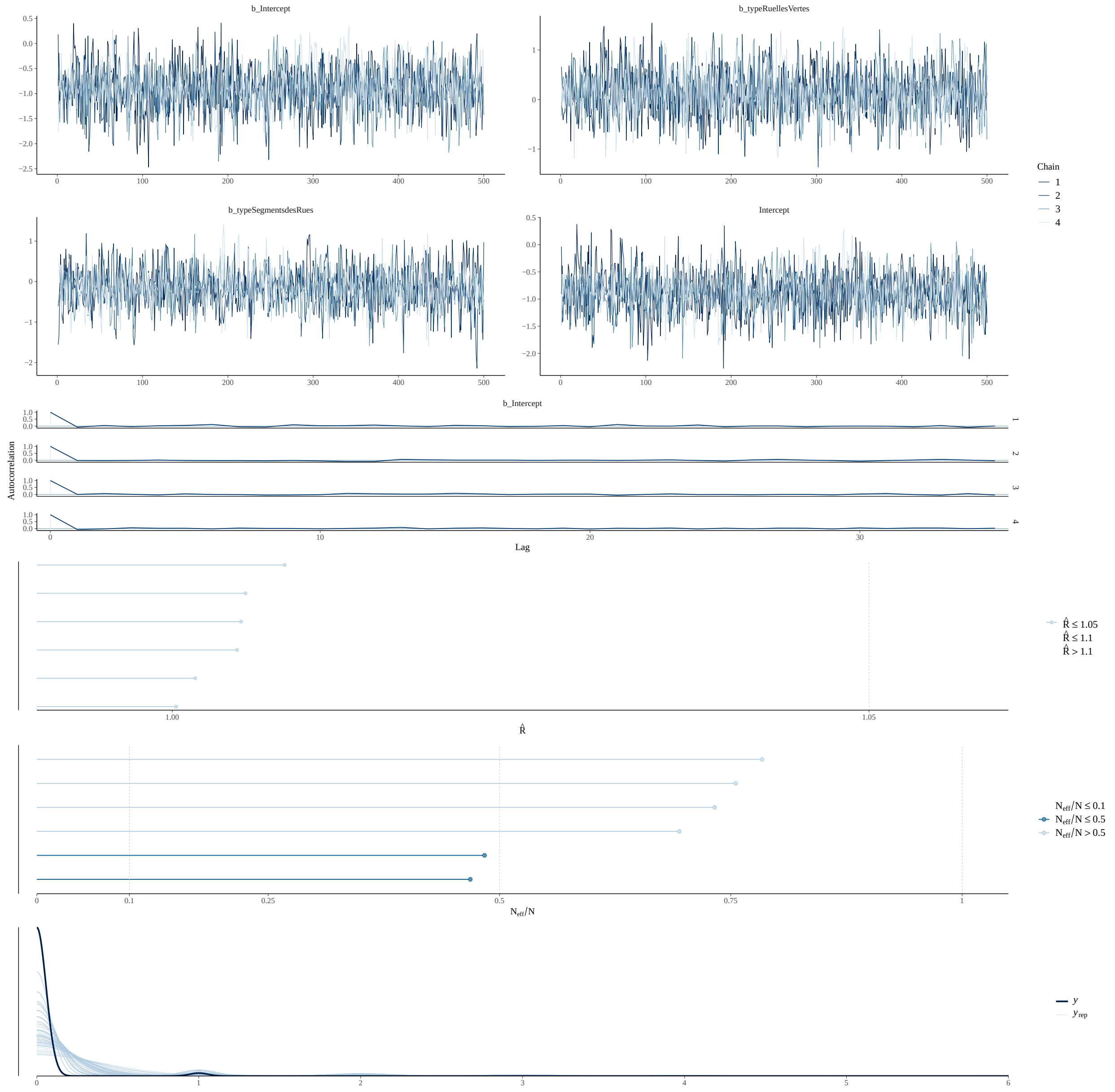

Force (3TF) - Tree Functional Trait Database. figshare. doi:10.6084/m9.figshare.14039504.v4.

bplant.org. 2024. Eastern Temperate Forests. *bplant.org.*

Canadian Wildlife Federation. 2024. Native Plant Encyclopedia. Encyclopedia.

Dirr, M. A., and K. S. Warren. 2019. *The Tree Book: Superior Selections for Landscapes, Streetscapes, and Gardens.* Portland, OR: Timber Press.

Farrar, J. L. 1995. *Trees in Canada.* Fitzhenry & Whiteside.

Firefly Atlas, C. Fallon, and R. Joyce. 2023. *Firefly Atlas Participant Handbook.*

Fryer, J. L. 2018. Tree species distribution maps from Little's "Atlas of United States trees" series. In: Fire Effects Information System.

Government of Canada, N. R. C. 2013. Natural Resources Canada: Trees, insects and diseases of Canada's forests - Index. December 31.

Institut National de Santé Publique du Québec, and Gouvernement du Québec. 2022. Canopée des six RMR du Québec 2022. Spatial. Partenariat Données Québec.

Little, E. L. 1980. *National Audubon Society Field Guide to Trees: Eastern Region.* North America. New York: Alfred A Knopf.

Magarik, Y. A. S., L. A. Roman, and J. G. Henning. 2020. How should we measure the DBH of multi-stemmed urban trees? *Urban Forestry & Urban Greening* 47: 126481. doi:10.1016/j.ufug.2019.126481.

Minister of Industry. 2010. EnviroStats. *EnviroStats* 4: 24.

Padvaiskas E, Richmond IC, Ziter CD. *In Review.* Forest structure but not tree diversity differs among urban woodlands with differing conservation status. *Ecoscience.*

Paquette, A., R. Sousa-Silva, F. Maure, E. Cameron, M. Belluau, and C. Messier. 2021. Praise for diversity: A functional approach to reduce risks in urban forests. *Urban Forestry & Urban Greening* 62: 127157. doi:10.1016/j.ufug.2021.127157.

Philp, K. 2024. zule-lab/katie-490: Completed (version v1.0.0). Zenodo. doi:10.5281/zenodo.10553246.

Picchi, M. S., L. Avolio, L. Azzani, O. Brombin, and G. Camerini. 2013. Fireflies and land use in an urban landscape: the case of Luciola italica L. (Coleoptera: Lampyridae) in the city of Turin. *Journal of Insect Conservation* 17: 797–805. doi:10.1007/s10841-013-9562-z.

QGIS Development Team. 2020. QGIS Geographic Information System (version 3.16). Hannover. QGIS Association.

The Morton Arboretum. 2024. Trees and Plants. *The Morton Arboretum*.

Tree Canada. 2019. *Canadian Urban Forest Strategy 2019 - 2024*.

United States Department of Agriculture, and Natural Resources Conservation Service. 2024. PLANTS Database.

Ville de Montréal. 2024. Actifs de voirie (Base de données complète - Chaussée, Îlot, Intersection, Trottoir, Zone). Spatial. Données Ouvertes Montréal.

Woody Invasives of the Great Lakes Collaborative. 2019. Woody Invasive Species. *Midwest Invasive Plant Network*. April 18.
